# Intermittent parathyroid hormone employs autonomous and non-autonomous mechanisms to drive osteogenesis from Ebf3-expressing skeletal progenitor cells

**DOI:** 10.64898/2026.05.21.726951

**Authors:** Byron S.K. Chan, Bingzi Dong, Majd George, Juwell Wu, Birol Ay, Daniel J. Brooks, Mary L. Bouxsein, Benjamin Z. Leder, Garyfallia Papaioannou, Karin Gustafsson, David T. Scadden, Takashi Nagasawa, Henry M. Kronenberg, Charles P. Lin, Marc N. Wein

## Abstract

How systemic hormonal signals coordinate stem cell fate decisions in adult tissues remains incompletely understood. In bone marrow, Cxcl12-abundant reticular (CAR) cells, marked by Early B-cell Factor 3 (Ebf3) expression, are multipotent mesenchymal progenitors that maintain the hematopoietic stem cell niche and serves as a major osteoblast progenitor source during adult bone remodeling. Using inducible lineage tracing coupled with single-cell transcriptomics and conditional genetics in mice, we show that intermittent parathyroid hormone (iPTH; teriparatide) drives osteogenesis from CAR cells by simultaneously engaging cell-intrinsic and cell-extrinsic mechanisms. Directly, iPTH suppresses lineage-enforcing transcription factors Ebf3, Ebf1, and Foxc1, thereby destabilizing progenitor identity and priming CAR cells for osteogenic commitment. Simultaneously, iPTH stimulates osteoclastic bone resorption, releasing TGFß which recruits these primed progenitors to bone surfaces, a process abolished by osteoclast depletion. Preventing CAR cell maturation via Sp7 deletion abrogates iPTH-induced bone gain, establishing these progenitors as essential mediators of bone anabolism. This coupled mechanism, in which intrinsic transcriptional priming converges with extrinsic niche remodeling, is conserved in human CAR cells from teriparatide-treated postmenopausal women, which show concordant suppression of EBF3 and FOXC1 and elevated TGFß-responsive gene signatures. These findings reveal a general principle by which a systemic hormone orchestrates tissue remodeling through simultaneous reprogramming of progenitor identity and remodeling of the niche microenvironment.

## Introduction

How systemic signals coordinate progenitor fate decisions in adult tissues is a fundamental question in stem cell biology. Progenitor differentiation depends on both cell-intrinsic transcriptional programs and extrinsic cues from the surrounding niche microenvironment^1,2^. How these two levels of regulation are integrated, and whether a single extracellular signal can simultaneously engage, both to drive and coordinate fate decisions, remains poorly understood. The bone marrow provides an experimentally tractable model for addressing this question in that it harbors progenitor cells, a complex multicellular niche, and is subject to regulation by systemic cues. Osteoblasts originate from early mesenchymal progenitor cells through multiple differentiation pathways^3–7^, and parathyroid hormone (PTH) acts on multiple cell types within the osteoblast lineage to regulate bone remodeling. Teriparatide, given once daily by subcutaneous injection as an intermittent parathyroid hormone (iPTH) therapy, is an effective and widely used medication to treat osteoporosis. The bone anabolic actions of iPTH are complex, but in part may be mediated by osteoblast progenitors^8^. However, osteoblast progenitors are heterogeneous, and multiple progenitor sources can contribute to the total osteoblast pool at different times and locations during skeletal growth and remodeling. Despite this, little is known about the effects of iPTH on specific subtypes of osteoblast progenitors.

CXC chemokine ligand 12 (Cxcl12)-abundant reticular (CAR) cells are a critical class of osteoblast progenitors during adult bone remodeling. These cells have a central role in supporting hematopoiesis, and also have the capacity to give rise to osteoblasts and adipocytes^9^. CAR cells are characterized by high expression of hematopoiesis-supportive paracrine factors including Cxcl12 and Stem cell factor (Kitl), as well as Leptin receptor (Lepr), and the transcription factors Forkhead box C1 (Foxc1)^10^ and Early B-cell Factor 3 (Ebf3). Ebf3, in particular, is a key transcription factor that maintains the ability of these cells to support hematopoiesis and inhibits their osteoblastic differentiation^11^.

CAR cells are a major source of osteoblast progenitors responsible for homeostatic bone remodeling at adulthood, compared to other progenitor sources that play more prominent roles during earlier postnatal skeletal growth^6,12,13^. Previously, we demonstrated that iPTH treatment in young/growing mice acts on a subset of growth-associated osteoblast progenitors marked by Sox9-CreERT2^14^. It is unclear whether these Sox9-expressing progenitors, versus other progenitor subtypes, respond to iPTH at skeletal maturity. Studies using the constitutive Lepr-Cre lineage tracing system^15–18^ have suggested that iPTH promotes osteoblastic differentiation of Lepr-expressing cells. However, the lack of inducible labelling of Lepr-expressing cells at skeletal maturity in this system raises the possibility that progeny of Lepr-expressing cells, rather than CAR cells themselves, may also contribute to iPTH-induced increases in osteoblasts.

The actions of iPTH concomitantly stimulate both bone formation and bone resorption. PTH directly targets cells within the osteoblast lineage^8,14,19^, but also exerts cell-extrinsic effects within the bone marrow microenvironment to enhance osteoblast differentiation. These non-cell-autonomous effects occur, in part, through iPTH-stimulated osteoclastic bone resorption, which liberates matrix-embedded growth factors such as insulin-like growth factor 1 (IGF1)^20^ and transforming growth factor ß-1 (TGFß1)^21^ that enhance osteoblastic differentiation. In addition to their resorptive activity, osteoclasts secrete anabolic coupling factors, or clastokines, that directly increase osteoblast numbers^22^. Additionally, PTH directly acts on matrix-embedded osteocytes to inhibit expression of the Wnt antagonist sclerostin^23^, thereby promoting osteoblast differentiation via Wnt signaling. While these cell-autonomous and non-cell-autonomous mechanisms are well established in mature osteoblast-lineage cells, it remains unclear whether and how iPTH employs similar mechanisms in early progenitors, such as CAR cells, to drive osteoblastic differentiation and bone formation.

Here, we used inducible Ebf3-CreERT2 and Sox9-CreERT2 lineage tracing systems, coupled with single-cell transcriptomics, to define the contributions of CAR cells and growth-associated osteoblast progenitors to iPTH-induced bone anabolism at skeletal maturity. We examined both cell-autonomous and non-cell-autonomous mechanisms through which iPTH regulates CAR cell fate, and investigated inter-cellular communication between osteoclasts and CAR cells during iPTH-stimulated bone remodeling. Finally, we tested whether the transcriptional programs governing iPTH action in murine CAR cells are conserved in human CAR cells from postmenopausal women receiving teriparatide.

## Results

### Intermittent PTH treatment recruits Ebf3-lineage cells to bone surfaces and induces their differentiation into mature osteoblasts

To assess the role of CAR cells and their descendants in response to intermittent PTH (iPTH), we performed lineage tracing of CAR cells using Ebf3-CreERT2; Rosa26-LSL-tdTomato; Bglap-GFP mice^11^. In this system, a single dose of tamoxifen treatment transiently activates Cre recombinase to induce the permanent expression of tdTomato in CAR cells and their descendants. Additionally, the Bglap-GFP (osteocalcin-GFP) transgene, which is highly expressed only by mature osteoblasts, affords visualization of labeled cell differentiation into mature osteoblasts. Ebf3-expressing cells were labeled with one dose of tamoxifen (2 mg) in 10-week-old mice. Thereafter, mice were treated with PTH amino acids 1-34 (100 µg/kg) or vehicle once per day and sacrificed 2 days, 10 days, and 2 months later (**Figure 1A**). No tdTomato+ cells were observed in the absence of tamoxifen treatment (**Supplemental Figure 1A, B)**.

**Figure 1.**
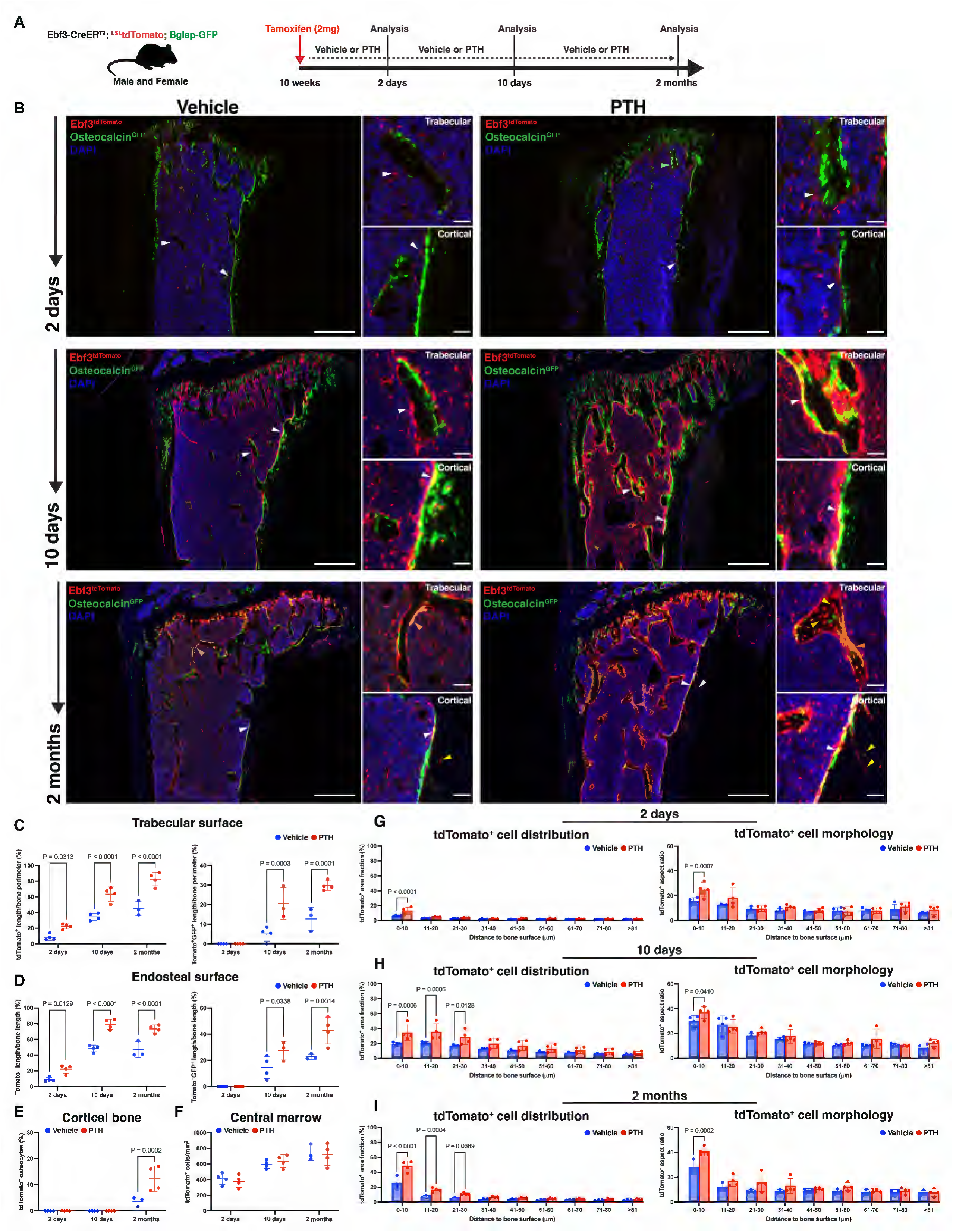
Intermittent parathyroid hormone treatment stimulates Ebf3-lineage cell recruitment to bone surfaces and promotes their osteoblastic differentiation. (**A**) Overview of lineage tracing strategy to study Ebf3-lineage cells using Ebf3-CreERT2; Rosa26-LSL-tdTomato; Bglap-GFP mice. Tamoxifen was administered at 10 weeks of age and sacrificed after 2 days, 10 days, and 2 months of iPTH or vehicle treatment. (**B**) Representative fluorescent images of the proximal tibia showing Ebf3-CreERT2-labeled tdTomato+ cells accumulate on bone surfaces (outlined by Bglap-GFP+ osteoblasts) and differentiate into mature osteoblasts (tdTomato+GFP+) and matrix-embedded osteocytes under steady state condition, and this process is enhanced with iPTH treatment. Red = Ebf3-CreERT2-tdTomato; green = Bglap-GFP; blue = DAPI. Scale bar = 500 µm (lower magnification) and 50 µm (higher magnification). (**C**) Quantification of tdTomato+ cells and tdTomato+GFP+ osteoblasts on metaphyseal trabecular and (**D**) endocortical bone surfaces, (**E**) tdTomato+ osteocytes in the cortical bone, and (**F**) tdTomato+ cells in the central bone marrow. Distribution (area fraction) and cell morphology (aspect ratio) analysis of tdTomato+ cells relative to defined distances from the endocortical bone surface in the proximal tibia following (**G**) 2 days, (**H**) 10 days, and (**I**) 2 months of iPTH or vehicle treatment. Statistical test: Two-way ANOVA with (**C-E**) treatment and time, or (**G-I**) treatment and specified distance bins as factors followed by Šidák’s multiple comparisons test. Data is expressed as mean ± SD, n = 3-4 mice/group.

In vehicle-treated mice at 2 days post-tamoxifen, labeled cells were distributed throughout the bone marrow and absent from bone surfaces (**Figure 1B**). There were also rare tdTomato+ cells present on periosteal surfaces. At 10 days post-tamoxifen, tdTomato+ cells were present on trabecular and endocortical bone surfaces and some differentiated into tdTomato+GFP+ osteoblasts (**Figure 1B, arrowheads**). Furthermore, labeled cells near and on bone surfaces appeared less reticular compared to those localized in the marrow. At 2 months post-tamoxifen in vehicle-treated mice, tdTomato+ and tdTomato+GFP+ cells continued to be present on bone surfaces. At this time point, but not earlier, tdTomato+ cells were embedded in cortical bone as terminally-differentiated osteocytes **(Figure 1B, yellow arrowheads**). In iPTH-treated mice, labeled cells were already localizing near trabecular and endocortical surfaces by 2 days post-tamoxifen and had adopted an elongated morphology. Following 10 days of iPTH treatment, there was an exuberant expansion of tdTomato+ cells on and near bone surfaces which was accompanied by enhanced differentiation into tdTomato+GFP+ osteoblasts. After 2 months of iPTH, the majority of trabecular and endocortical bone surfaces were occupied by tdTomato+ cells and tdTomato+GFP+ osteoblasts. Additionally, there was a greater density of tdTomato+ osteocytes in the trabecular and cortical bone upon iPTH treatment (**Figure 1B)**.

Quantification across skeletal compartments confirmed that iPTH progressively recruits Ebf3-lineage cells to bone surfaces and drives their osteoblastic differentiation over time **(Figures 1C–F, Supplementary Figures 2A–D)**. At 2 days, iPTH increased labeled cells on metaphyseal and diaphyseal surfaces without evidence of osteoblast differentiation. By 10 days, both progenitor and osteoblast numbers were significantly increased at trabecular and endocortical sites. After 2 months, labeled osteocytes accumulated in cortical bone, an effect seen at both metaphyseal and diaphyseal regions, while central marrow tdTomato+ cell numbers remained unchanged at all timepoints. These effects were comparable in male and female mice **(Supplementary Figure 2E)**.

To further assess the effects of time and iPTH on tdTomato+ cell distribution and morphology relative to bone surfaces, MATLAB-based image analysis (see Methods) was performed. After 2 days, iPTH increased the area fraction (proportion of tissue area occupied by tdTomato+ pixels) and aspect ratio (major axis length/minor axis length of tdTomato+ cell) within 10 µm of endocortical bone surfaces **(Figure 1G)**. After 10 days (**Figure 1H)** and 2 months **(Figure 1I)**, iPTH treatment increased the tdTomato+ area fraction within 30 µm distance from endocortical bone. Cell shape changes, however, were confined to within 10 µm of bone, indicating that PTH-induced morphological remodeling is spatially restricted to the endocortical surface. These results demonstrate that some Ebf3-lineage cells differentiate into osteoblasts and, eventually, osteocytes, over time. Moreover, Ebf3-expressing progenitors may respond to PTH via enhanced recruitment to trabecular and endocortical bone surfaces and subsequent differentiation into mature osteoblasts.

### Adipogenic differentiation of Ebf3-lineage cells is inhibited by PTH

As CAR cells can become bone marrow adipocytes^11,24^ and PTH inhibits progenitor adipogenesis^8,14^, we assessed whether iPTH alters the adipogenic fate of Ebf3-lineage cells **(Supplementary Figure 3A)**. 2 months after tamoxifen-induced labeling, the total number of perilipin+ and tdTomato+perilipin+ adipocytes in the proximal tibia metaphysis increases in vehicle-treated mice and was significantly reduced by iPTH **(Supplementary Figure 3B, C)**. Thus, iPTH may skew the fate of Ebf3-lineage cells from adipogenic to osteogenic commitment.

### Ebf3-lineage osteogenic/stromal subpopulations are increased by PTH

To characterize the phenotypic heterogeneity of Ebf3-lineage cells, we performed flow cytometric analysis on enzymatically-digested marrow plus bone fragments using a panel of cell surface markers that identifies early subpopulations of skeletal stem and progenitors (mSSC, pre-BCSP, and BCSP) that give rise to lineage-restricted osteochondrogenic (Thy1+) and HSC-supporting stroma (6C3+) subpopulations^25^ (**Supplementary Figure 4A**). First, we defined where initially labeled Ebf3-lineage CAR cells track in these FACS-defined subpopulations. 10-week-old Ebf3-CreERT2; Rosa26-LSL-tdTomato; Cxcl12-GFP transgenic mice were studied 2 days post-tamoxifen (**Supplementary Figure 5A**). Flow cytometry revealed Ebf3-CreERT2 marks the majority (∼85%) of Cxcl12-GFP expressing cells (**Supplementary Figure 5B)**. Moreover, the parental Ebf3-lineage CAR cells identified as Cxcl12-GFP+tdTomato+ cells predominantly occupy the 6C3+ FACS subpopulation (Thy1–6C3+; **Supplementary Figure 5C)**.

Next, we assessed the effects of iPTH on the distribution of Ebf3-lineage cells with respect to these skeletal stem and progenitor subpopulations. At 2 days post-tamoxifen (**Figure 2A)**, iPTH treatment led to modest but significant increases in CD45–Ter119–CD31– (hereafter Lineage–) tdTomato+ cells **(Figure 2B)** while the subsets of tdTomato+ skeletal progenitors did not change (**Figure 2C)**. Following 10 days of iPTH (**Figure 2D)**, there were no differences in overall numbers of Lineage–tdTomato+ cells (**Figure 2E)**. However, within tdTomato+ cells, iPTH decreased numbers of pre-BCSP cells while increasing 6C3+ cells (**Figure 2F**). Therefore, within the Ebf3-lineage, iPTH promotes the differentiation of early subsets of skeletal progenitors (Pre-BCSP) towards 6C3+ stromal cells.

**Figure 2.**
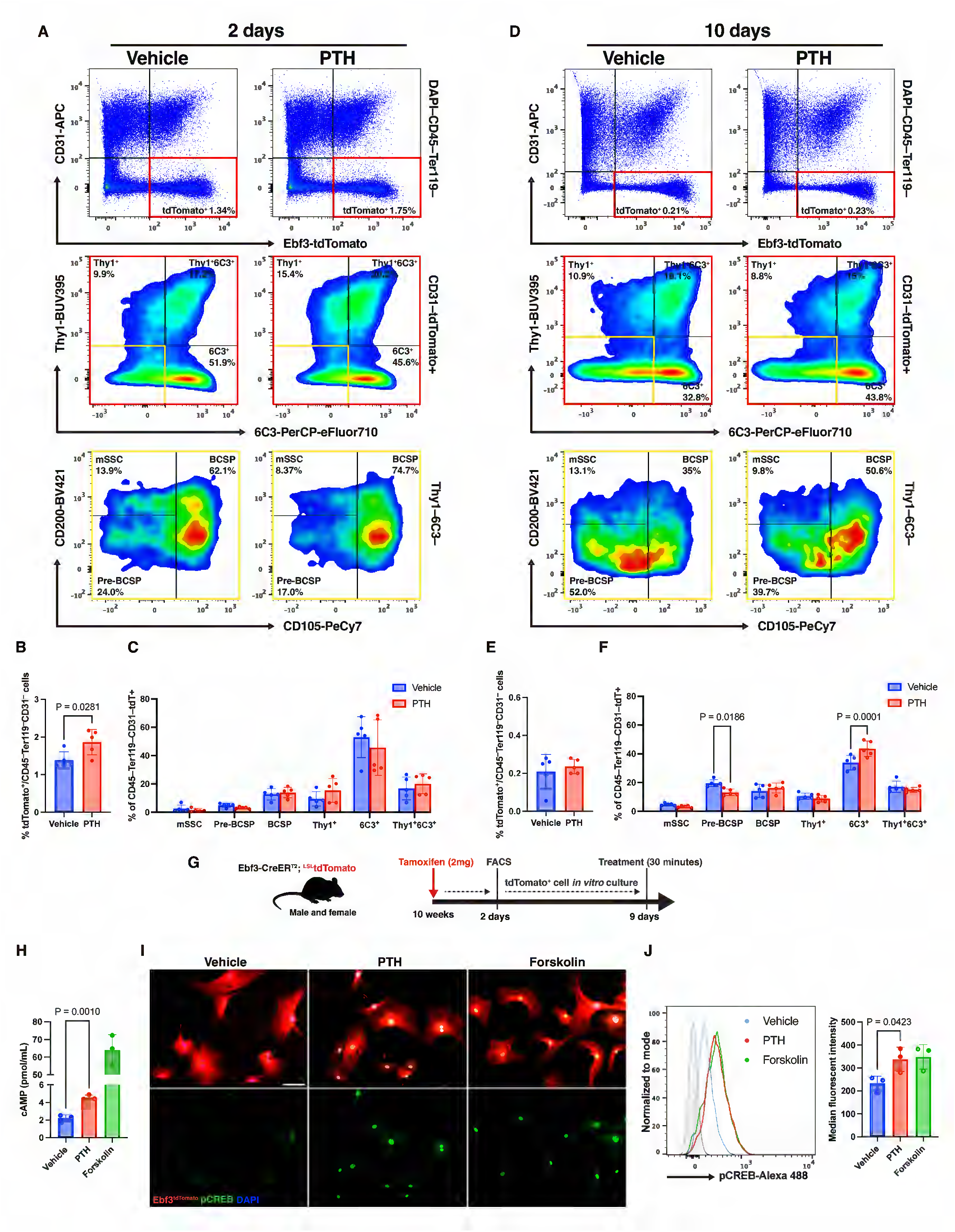
Ebf3-lineage cells can directly respond to iPTH. Representative flow cytometry plots and analysis of non-hematopoietic Ebf3-CreERT2-labeled tdTomato+ cells and their skeletal stem and progenitor cell surface markers following 2 (**A-C)** and 10 days (**D-F)** of iPTH or vehicle treatment. Red box: CD45–Ter119–CD31–tdTomato+ fraction. Yellow Box: CD45–Ter119–CD31–tdTomato+Thy1–6C3– fraction. mSSC: Mouse skeletal stem cell; Pre-BCSP: Pre-bone/cartilage/stromal progenitor; BCSP: Bone/cartilage/stromal progenitor. (**G**) Experimental overview for *in vitro* culturing of FACS-isolated non-hematopoietic Ebf3-CreERT2-labeled tdTomato+ cells. (**H**) Cyclic adenosine monophosphate (cAMP) analysis, (**I**) immunocytochemical staining and (**J**) flow cytometry analysis for phospho-CREB, of *in vitro* cultured tdTomato+ cells thirty minutes after PTH, forskolin, or vehicle treatment. Statistical test: Student’s T-test (**B, E, H, and J**) and Two-way ANOVA with treatment and subpopulations (**C and F**) as factors followed by Šidák’s multiple comparisons test. Data is expressed as mean ± SD, n = 3-5 mice/group.

Across multiple experiments (see below), we observed that increased tdTomato+ cells detected by histology were not consistently reflected by flow cytometry. We speculated that our serial enzymatic digestion inefficiently liberates strongly bone-adherent tdTomato+ cells (presumably pre-osteoblasts, osteoblasts, and osteocytes). RT-qPCR on bone fragments remaining post-serial digestion showed higher tdTomato mRNA with iPTH, suggesting a greater number of tdTomato+ cells remains associated with bone fragments following iPTH (**Supplementary Figure 5D**).

### Ebf3-lineage cells can respond directly to PTH

To test potential direct actions of PTH on Ebf3-lineage cells, Lineage–tdTomato+ cells were isolated by FACS 2 days after tamoxifen treatment in 10-week-old Ebf3-CreERT2; Rosa26-LSL-tdTomato mice. After short-term (7 day) *in vitro* culture (**Figure 2G**), cells were treated for 30 minutes with PTH which led to increases in cAMP (**Figure 2H**) and phospho-CREB (**Figure 2I, J**). As detailed below, *Pth1r* mRNA expression is also noted in CAR cells and their descendants. These results indicate that Ebf3-lineage cells can directly respond to PTH for their commitment towards the osteogenic differentiation. iPTH can increase osteoblast progenitor and osteoblast numbers via anti-apoptotic mechanisms^14,26^. Therefore, we assessed whether iPTH affected the proliferation and apoptosis of Ebf3-lineage cells by EdU and Annexin V staining, respectively. After 10, but not 2, days, iPTH decreased the percentage of Annexin V+tdTomato+ cells (**Supplemental Figure 6A)**. iPTH did not impact the percentage of EdU+tdTomato+ cells (**Supplemental Figure 6B)**.

### Transcriptomic effects of PTH on Ebf3-lineage cells

To understand the mechanisms by which iPTH promotes osteoblastic differentiation of Ebf3-lineage cells, we performed single cell RNA-sequencing on FACS-isolated CD45–Ter119–tdTomato+ cells following 2 and 10 days of vehicle or iPTH treatment. Mice were treated with PTH or vehicle 30 minutes prior to sacrifice to capture acute transcriptomic effects of PTH at both time points. After merging all libraries with Harmony^27^, an algorithm for integrating multiple single cell RNA-sequencing datasets, tdTomato mRNA was expressed in all cell clusters (**Supplemental Figure 7A**).

Unsupervised clustering revealed 15 distinct groups of Ebf3-lineage cells: 6 CAR, 1 pre-osteoblast, 1 mature osteoblast, 3 periosteal, 2 pericyte, 1 endothelial, and 1 myogenic (**Figure 3A**). Relative to other clusters, CAR subsets were identified based on the highest expression of *Cxcl12, Lepr, Ebf3, Adipoq*, and *Foxc1*. A pre-osteoblast cluster showed high expression of *Limch1, Kcnk2, Wif1, Dlx5, Lrp4, Spp1*, and *Alpl*. Notably, the expression of these pre-osteoblastic genes overlaps with genes that are expressed in the CAR6 cluster, suggesting that this may be an osteogenic-primed CAR subset (so-called OsteoCAR), as other groups have reported^28,29^. The mature osteoblast cluster was identified based on highest *Col1a1*, *Bglap*, *Dmp1*, and *Pth1r* expression and lower expression of the pre-osteoblast defining genes. Periosteal clusters were identified based on expression of *Prg4*, *Mfap5*, and *Clec3b*^28^. The remaining clusters represented pericytes, endothelial cells, and a rare myogenic population, identified by canonical markers **(Figure 3B)**.

**Figure 3.**
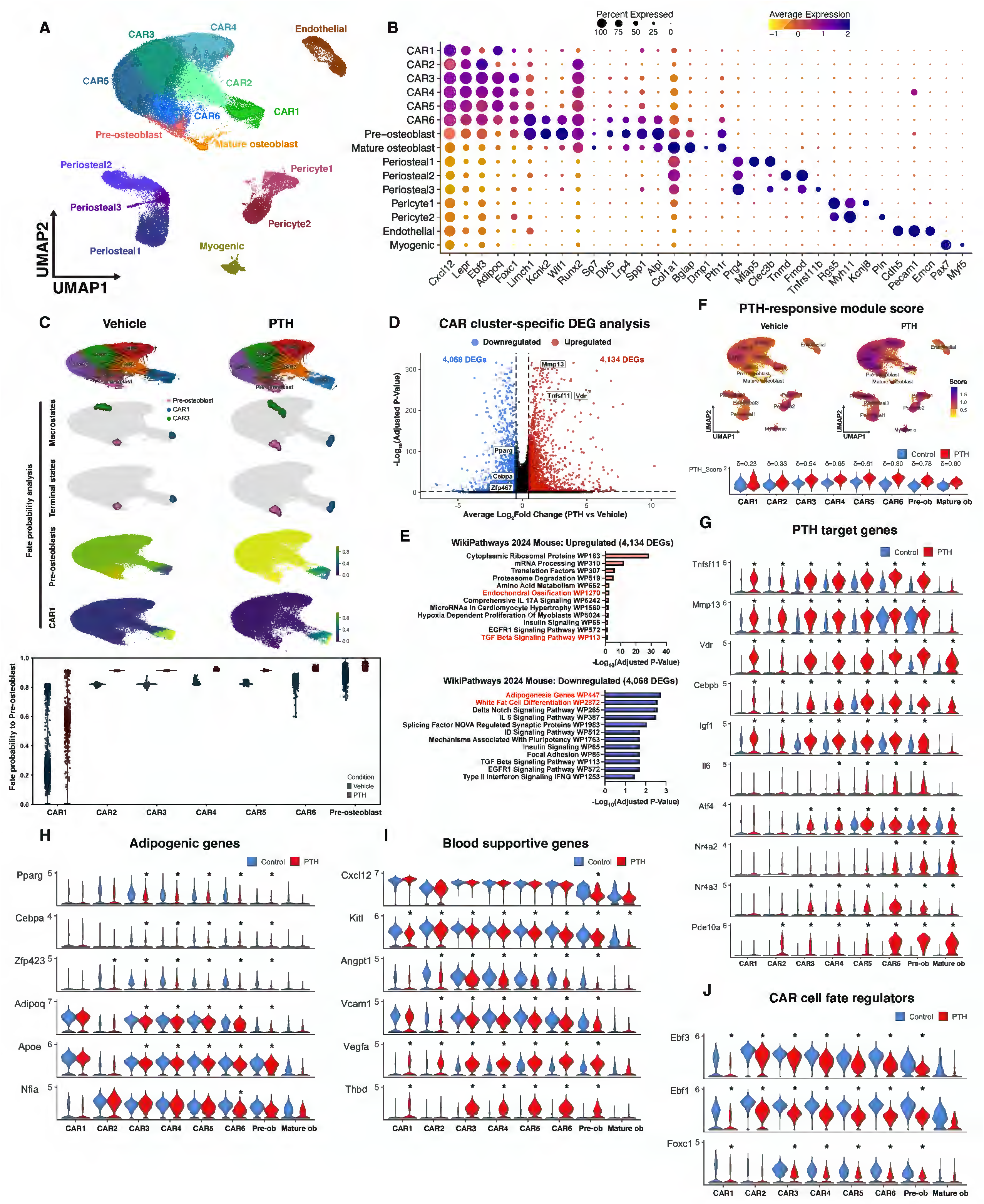
iPTH destabilizes the CAR cell phenotype. **(A)** Harmony-integrated UMAP of single cell RNA-sequencing data from FACS-isolated CD45–Ter119–tdTomato+ cells labeled by Ebf3-CreERT2 following 2 and 10 days of iPTH or vehicle treatment (2 days: vehicle = 13,058 cells and PTH = 12,214 cells; 10 days: vehicle = 34,735 cells and PTH = 19,809 cells; n = 4 mice/group). (**B**) Dot plot summarizing the expression levels of cell-type marker genes across subclusters within Ebf3-lineage cells. Dot size represents the fraction of positive cells within the cluster expressing the gene of interest, and color scale denotes average gene expression level. Purple = higher expression and yellow = lower expression. (**C**) RNA velocity-based fate probability analysis using the CellRank^42^ to infer Ebf3-lineage CAR cell fates at steady state or in response to PTH. Color scale represents fate probability of each cell. Violin plots show the fate probability of Ebf3-lineage CAR subclusters transitioning toward the pre-osteoblast cluster under steady state or PTH-treated conditions. (**D**) Volcano plot showing the total number of significant differentially expressed genes (DEGs) of all Ebf3-lineage CAR subclusters regulated by PTH (adjusted P-value < 0.05 and |log2FC| > 0.5). (**E**) Gene set enrichment analysis of significantly up- and down-regulated DEGs by PTH using the WikiPathways 2024 Mouse database. Top 12 significantly enriched gene sets are shown (adjusted P-value < 0.05). (**F**) PTH-responsive module scores visualized by Feature plots of each Ebf3-lineage cell (top) and violin plots (bottom) across Ebf3-lineage CAR and osteoblast clusters in response to iPTH or vehicle treatment. Purple = higher expression and yellow = lower expression. Cliff’s delta (δ) = non-parametric effect size used to measure differences between module score distributions. (**G**) Violin plots showing the expression levels of canonical PTH target genes, (**H**) adipogenic genes, (**I**) blood supportive genes, and (**J**) CAR cell fate regulating transcription factor genes, across Ebf3-lineage CAR and osteoblast clusters in response to iPTH or vehicle treatment. Differentially expressed genes between PTH and control cells within each cluster was assessed using the MAST hurdle model^74^. Asterisk (*) denotes significance (adjusted P-value < 0.05 and |log2FC| > 0.5).

To further resolve the heterogeneity within the 6 different CAR clusters, we performed DEG analysis of each CAR cluster versus all the other CAR subsets (**Supplemental Figure 7B**). While all CAR subsets express high levels of *Cxcl12*, its expression in CAR1 is the highest. The CAR1 subset is also enriched with genes such as *Adipoq*, *Sar1a*, *Srpr*, *P4hb*, *Os9*, and *Nfe2l1* that are involved with secretion^30,31^ (**Supplementary Figure 7C)** and may be most similar to previous reports of an adipogenic CAR subset^29,32^. CAR2 has the lowest *Cxcl12* and highest *Ebf3* expression and is enriched in genes (like *Dock1*, *Trio*, *Arhgap35*) which play roles in cytoskeletal remodeling and cell migration^33–35^. CAR3 is enriched in expression of *Foxc1*, *Snai2*, and *Abcg2*, all genes associated with stem and progenitor cells^10,36,37^. CAR4 is enriched in *Pecam1*, *Cx3cr1*, and *Sema4a* which may be involved with endothelial cell interactions^38^. CAR5 is enriched in expression of *Ier5* and *Nfkbia*, which are involved in early stress responses^39,40^. As noted above, CAR6 is enriched in expression of *Kcnk2*, *Alpl*, *Wif1*, *Limch1*, *Lrp4*, *Spp1*, and *Nav2*, a profile of enriched markers that is consistent with an osteogenic CAR subset^28,29,41^.

To study the relationship between CAR subsets and how they are regulated by iPTH, we performed RNA velocity-based fate probability analysis using CellRank^42^ (**Figure 3C**). Three putative macrostates, which represents cells that share similar transition probabilities, were predicted: CAR1, CAR3, and pre-osteoblast. CAR1 and pre-osteoblast were identified as terminal states with high self-transition probabilities suggesting a differentiated transcriptional state. Notably, when comparing between all CAR clusters in the vehicle treated group, CAR6 shows the highest fate probability towards pre-osteoblasts. Remarkably, iPTH treatment increases the fate probability of all CAR subsets towards pre-osteoblasts and decreases their fate probability towards adipocyte-primed CAR1 cells (**Figure 3C**).

To understand the transcriptional mechanisms associated with iPTH-induced osteoblastic CAR cell differentiation, we next performed cluster-specific DEG analysis comparing vehicle versus iPTH in all CAR and osteoblast clusters. A total of 4,134 upregulated and 4,068 downregulated DEGs were identified to be altered by PTH **(Figure 3D)**. Several canonical PTH target genes like *Tnfsf11* (RANKL), *Mmp13,* and *Vdr* were identified as upregulated DEGs, and adipogenesis-related genes *Pparg, Zfp467,* and *Cebpa* were identified as downregulated DEGs. Next, we performed gene set enrichment analysis (GSEA) of all the upregulated and downregulated DEGs (**Figure 3E)**. The top pathways with significant (p_adj_ < 0.05) enrichment amongst the upregulated DEGs are those associated with protein translation, endochondral ossification, and TGFß signaling. The top pathways amongst genes downregulated by PTH are those related to adipogenesis, Notch signaling, and mechanisms associated with pluripotency.

To assess overall PTH responsiveness, we computed a module score from 13 canonical PTH target genes (*Tnfsf11, Fos, Fosl1, Junb, Egr1, Crem, Nr4a2, Nr4a3, Nfkbia, Il6, Pde4d, Igf1,* and *Pde10a*). As expected, iPTH increased this module score in CAR subsets, pre-osteoblast, and mature osteoblast clusters. Of all CAR subsets, CAR6 showed the largest increase module score with PTH treatment (**Figure 3F)**.

Next, we queried the expression levels of canonical PTH target genes across the Ebf3-lineage CAR and osteoblast clusters. Expression levels of *Tnfsf11*, *Mmp13*, *Vdr*, and *Cebpb*, were induced by PTH in all CAR and osteoblast clusters. *Igf1* was increased in all CAR and pre-osteoblast clusters. Interestingly, other PTH target genes are uniquely increased in specific CAR subsets. For example, *Il6* expression is increased in CAR4/5/6 and pre-osteoblast; *Atf4* only in CAR6; *Nr4a2* and *Nr4a3* in CAR3-6 and osteoblast clusters; and *Pde10a* was increased in CAR2-6 and osteoblast clusters (**Figure 3G**).

As demonstrated earlier, iPTH exerts anti-apoptotic actions in Ebf3-lineage cells (**Supplementary Figure 6A)**. We queried the list of upregulated and downregulated DEGs that may be involved in apoptosis and observed that PTH downregulates a variety of pro-apoptotic genes (**Supplementary Figure 8A)**. An anti-apoptosis/survival module score was computed using a curated gene module (*Bcl2, Bcl2l1, Mcl1, Bcl2a1a, Bcl2a1b, Cflar, Xiap, Birc2, Birc3, Traf1, Traf2, Akt1. Akt2, Rel, Rela, Nfkb1, Nfkbia*) to further assess the anti-apoptotic effects of iPTH on Ebf3-lineage CAR cells and osteoblasts. In response to PTH, anti-apoptosis module scores increased, albeit to a more modest degree than the PTH-responsive score, across CAR2-6, pre-osteoblast, and mature osteoblast clusters, with CAR6 and mature osteoblast clusters exhibiting the strongest scores (**Supplementary Figure 8B)**.

iPTH suppressed the adipogenic transcription factors *Pparg*, *Cebpa*, and *Zfp423*^43,44^ across CAR3–6 and pre-osteoblast clusters **(Figure 3H)**. CAR1 cells were a notable exception: they expressed the highest levels of *Adipoq* and *Apoe*, and neither gene changed with PTH treatment, consistent with CAR1 representing an adipogenic-committed subset. The osteoblast differentiation factors *Sp7* and *Runx2*, along with their target *Bglap*, were acutely downregulated in CAR6 and osteoblast clusters **(Supplementary Figure 7D)**, a pattern previously described as part of the acute transcriptional response to PTH in osteoblast-lineage cells^45^. Despite unchanged *Col1a1*, iPTH upregulated *Col3a1*, *Col4a1*, and *Col6a1* in CAR3–6 and pre-osteoblasts, suggesting active remodeling of the extracellular matrix during CAR cell differentiation.

CAR cells are crucial components of the hematopoietic supportive niche. While short term iPTH treatment did not affect *Cxcl12* levels in any CAR subsets, *Kitl* was significantly decreased in all CAR and osteoblast clusters (**Figure 3I**). Other marrow niche supportive genes such as *Angpt1*^46^ and *Vcam1*^47^ were downregulated by PTH in CAR2-6 and pre-osteoblasts. In contrast, *Vegfa* and *Thbd* was increased in CAR and pre-osteoblast clusters. Thus, PTH actions on CAR subsets modulates the expression of paracrine-acting genes that participate in inter-cellular communication in the bone marrow microenvironment.

Given the effects of PTH on osteogenic differentiation of CAR cells, we queried expression levels of critical transcription factors (*Ebf1, Ebf3, and Foxc1*) that enforce the CAR cell phenotype^10,11^. *Ebf1* and *Ebf3* prevent osteoblastic differentiation of CAR cells^11^, and expression of both of these factors (which show lower expression in pre-osteoblasts and osteoblasts versus CAR subsets) is significantly reduced by PTH (**Figure 3J**). Expression of the essential CAR cell-enforcing TF, *Foxc1,* was also significantly downregulated in response to PTH. Taken together, these findings highlight the heterogeneity of CAR cells and their subset-specific responses to PTH. Most notably, these data suggest that PTH destabilizes the CAR cell state by suppressing Ebf3, Ebf1, and Foxc1 expression to facilitate their differentiation into osteoblasts.

### A subset of Ebf3-lineage endothelial cells does not change in numbers in response to PTH

Osteogenesis and angiogenesis are intricately linked during bone growth and response to parathyroid hormone^48,49^. Subsets of vasculature-associated CAR cells have been previously identified to support the hematopoietic niche^5^. Single-cell RNA-sequencing identified two distinct CD31-expressing populations within Ebf3-lineage cells: a true endothelial cluster expressing canonical markers (*Pecam1, Emcn, Flt1, Tek, Cdh5, Ptprb*) and CAR4, which expresses lower CD31 without other endothelial-specific transcripts (**Figures 3A, B; Supplementary Figures 7C, 9A**). Immunostaining for CD31 and endomucin confirmed that a subset of labeled cells displayed endothelial morphology; notably, these CD31+tdTomato+ cells were absent from bone surfaces (**Supplementary Figure 9B**). Flow cytometry showed iPTH did not affect the proportion of CD31+tdTomato+ cells within the non-hematopoietic labeled population (**Supplementary Figure 9C**).

### Sox9-lineage cells rarely differentiate into mature osteoblasts following PTH treatment in skeletally mature mice

Sox9-CreERT2 marks osteoblast progenitors in the growth plate and perichondrium that respond to iPTH in early postnatal mice^6,14^. To determine whether this progenitor source remains active at skeletal maturity, we performed Sox9-lineage tracing at two ages. When labeled at 3 weeks of age, Sox9-lineage cells responded to iPTH with increased progenitor and osteoblast numbers in the primary spongiosa and cortical bone **(Supplemental Figure 10A, B)**, and is consistent with previous findings^14^. Single-cell RNA-sequencing of these young-labeled cells identified mature osteoblast, pre-osteoblast, and CAR cell clusters **(Supplementary Figures 10C, D)**, the latter consistent with CAR cells being Sox9-lineage descendants during skeletal growth^6,50^. In contrast, Sox9-lineage cells labeled at 10 weeks of age, using an identical tamoxifen pulse-chase protocol, showed no iPTH-induced changes in cell numbers or osteoblast differentiation **(Supplementary Figure 10E–G)**. Single-cell RNA-sequencing revealed that these adult-labeled cells consist predominantly of chondrocytes (*Prg4*, *Acan*, *Col2a1*), with PTH target genes induced in Pth1r-expressing chondrocyte clusters but no osteoblast or CAR cell clusters present **(Supplementary Figure 10H–J)**. Thus, while Sox9-lineage progenitors contribute to osteoblasts and give rise to CAR cells during postnatal growth, they are supplanted by Ebf3-lineage CAR cells as the primary iPTH-responsive osteoprogenitor pool at skeletal maturity.

### PTH-induced osteoblastic differentiation of Ebf3-lineage cells is inhibited by Sp7 deletion

Osterix (*Sp7)* is an essential transcription factor for osteoblast differentiation^51^. In our scRNA-seq data of Ebf3-lineage cells we noted that *Sp7* is expressed only in CAR6, pre-osteoblasts, and osteoblasts, and not in CAR1-5 subsets **(Figure 4A).** We deleted Sp7 in Ebf3-lineage cells in Sp7 f/f mice (Ebf3-CreERT2; Sp7 f/f; Rosa26-LSL-tdTomato; Bglap-GFP). Wild type controls (Ebf3-CreERT2; Sp7 +/+; Rosa26-LSL-tdTomato; Bglap-GFP) and Sp7 f/f mice were pulsed with a single dose of tamoxifen to induce tdTomato reporter expression and deletion of Sp7 in Ebf3-lineage cells followed by vehicle or PTH treatment for 10 days **(Figure 4B).**

**Figure 4.**
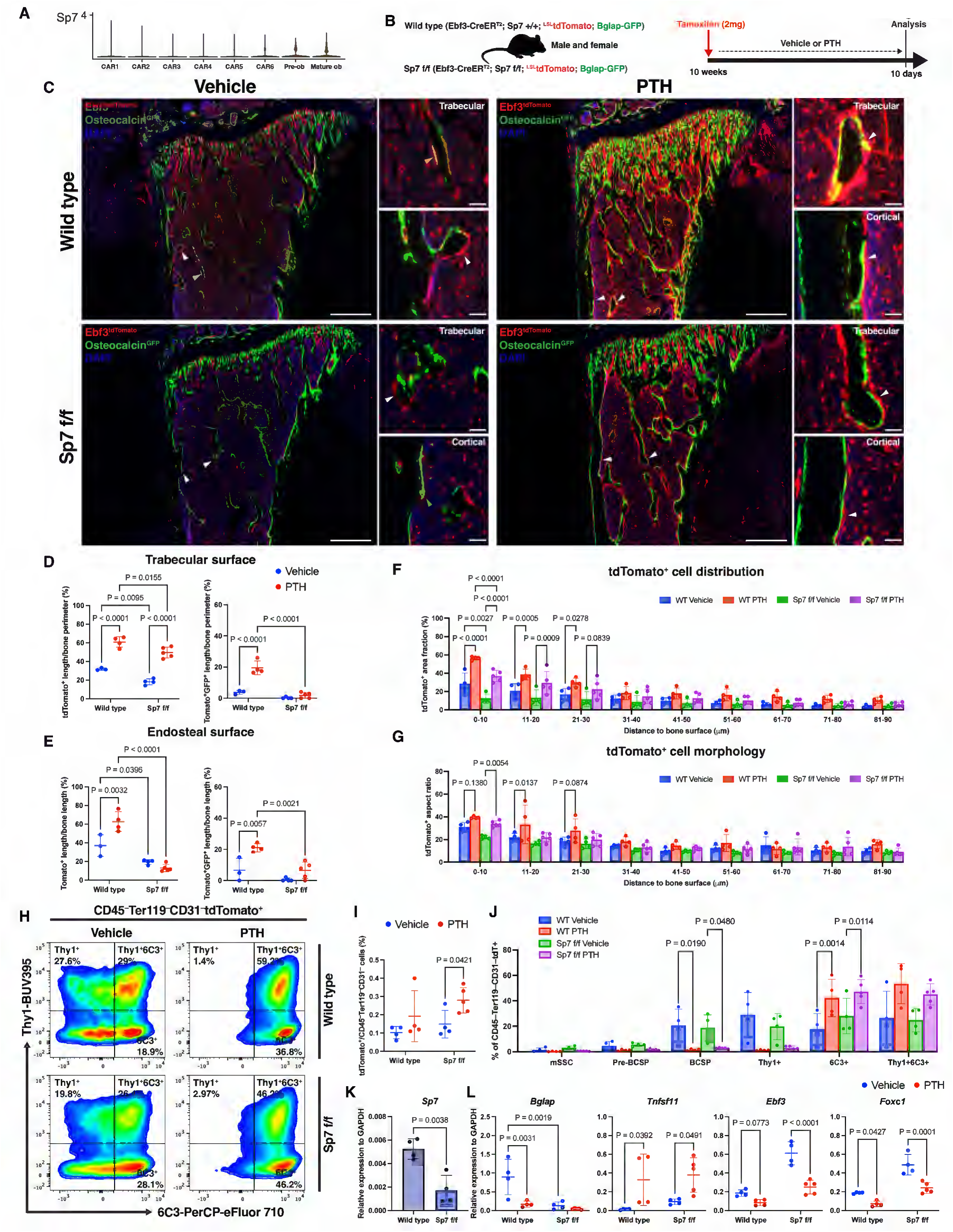
Sp7 is required for the iPTH-stimulated maturation of Ebf3-lineage cells into osteoblasts. **(A)** Violin plots showing Sp7 expression level across Ebf3-lineage CAR and osteoblast clusters. (**B**) Overview of lineage tracing strategy to study the role of Sp7 in Ebf3-lineage cells following 10 days of iPTH or vehicle treatment. (**C**) Representative fluorescent images of the proximal tibia from wild type and Sp7 f/f mice following 10 days of iPTH or vehicle treatment. Red = Ebf3-CreERT2-tdTomato; green = Bglap-GFP; blue = DAPI. Scale bar = 500 µm (lower magnification) and 50 µm (higher magnification). (**D**) Quantification of tdTomato+ cells and tdTomato+GFP+ osteoblasts on metaphyseal trabecular and (**E**) endocortical bone surfaces of wild type and Sp7 f/f mice following 10 days of iPTH or vehicle treatment. (**F**) Distribution (area fraction) and (**G**) cell morphology (aspect ratio) analysis of tdTomato+ cells relative to defined distances from bone surfaces in the proximal tibia of wild type and Sp7 f/f mice following 10 days of iPTH or vehicle treatment. (**H**) Representative flow cytometry plots and analysis of non-hematopoietic Ebf3-CreERT2-labeled tdTomato+ cells (**I**) and their skeletal stem/progenitor cell surface markers (**J**) from wild type and Sp7 f/f mice following 10 days of iPTH or vehicle treatment. (**K and L)** RT-qPCR analysis of FACS-isolated Lineage–tdTomato+ cells from wild type and Sp7 f/f mice following 10 days of iPTH and vehicle treatment. Statistical test: Two-way ANOVA with treatment and genotype (**D, E, I, and L**), or treatment and specified distance bins (**F and G**), or treatment and subpopulations (**J**) as factors followed by Šidák’s multiple comparisons test, and Student’s T-test (**K**). Data is expressed as mean ± SD, n = 4-5 mice/group.

At baseline, Sp7 f/f mice showed fewer labelled cells on metaphyseal trabecular and endocortical surfaces than wild type controls, with a similar trend at diaphyseal endocortical sites **(Figures 4C–E, Supplemental Figure 11A)**. In wild type mice, iPTH increased labeled progenitors and osteoblasts across all examined surfaces, consistent with prior results. In Sp7 f/f mice, iPTH still recruited labeled progenitors to trabecular surfaces but not endocortical surfaces, and, as expected, failed to increase tdTomato+GFP+ osteoblasts at any site. Central marrow numbers and periosteal progenitor responses were comparable between genotypes, and osteoblast differentiation was not observed on periosteal surfaces in either group **(Supplemental Figures 11B–D)**. Spatially, iPTH increased labeled cell density and elongation within 0–20 µm of endocortical bone in both genotypes; however, within the 0–10 µm zone where pre-osteoblasts and osteoblasts reside, Sp7 deletion specifically impaired the iPTH-stimulated increase in cell number **(Figures 4F, G)**. Flow cytometry showed no difference in overall Lineage–tdTomato+ cell numbers at baseline, and iPTH modestly increased these cells in both genotypes **(Figures 4H, I)**. The iPTH-induced shift from upstream BCSPs to downstream 6C3+ stromal cells was preserved across genotypes, indicating that early progenitor differentiation proceeds independently of Sp7 **(Figure 4J)**. RT-qPCR on sorted labeled cells confirmed reduced *Sp7* and *Bglap* mRNA expression in Sp7 f/f mice, while iPTH-induced changes in *Tnfsf11*, *Ebf3*, and *Foxc1* were comparable between genotypes **(Figures 4K, L)**.

Overall, these results indicate that the early stages of CAR cell differentiation towards osteoblasts, such as increases in stromal 6C3+ cells and their initial recruitment to bone surfaces, occur independently of Sp7. However, Sp7 plays a key role in controlling the final stages of osteoblast maturation (recruitment close to endocortical bone surfaces and expression of *Bglap*) at steady state and in response to iPTH.

### Non-cell-autonomous recruitment of Ebf3-lineage cells to bone surfaces

Since Ebf3-lineage cells can directly respond to PTH *in vitro* (**Figure 2**), and scRNA-seq indicates that its receptor, Pth1r, expression is present in CAR3-6 and osteoblasts (**Figure 5A)**, we next tested whether iPTH induces osteoblastic differentiation of CAR cells in a cell-autonomous manner *in vivo* by studying Pth1r f/f mice (Ebf3-CreERT2; Pth1r f/f; Rosa26-LSL-tdTomato; Bglap-GFP) and wild type control animals (Ebf3-CreERT2; Pth1r +/+; Rosa26-LSL-tdTomato; Bglap-GFP) (**Figure 5B)**.

**Figure 5.**
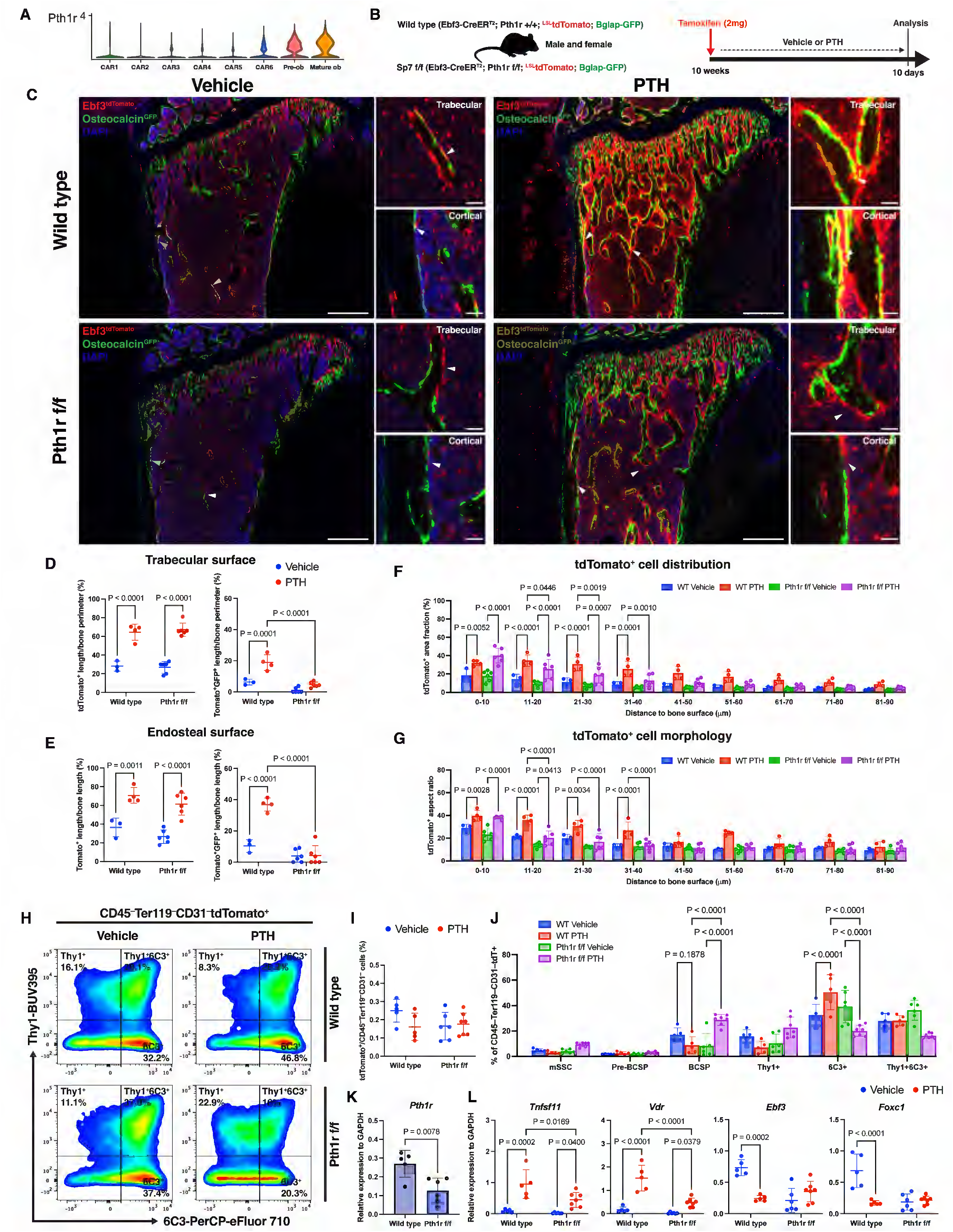
Pth1r is required for iPTH-mediated suppression of CAR cell-enforcing transcription factors in Ebf3-lineage cells and for their maturation into osteoblasts. **(A)** Violin plots showing Pth1r expression level across Ebf3-lineage CAR and osteoblast clusters. (**B**) Overview of lineage tracing strategy to study the role of Pth1r in Ebf3-lineage cells following 10 days of iPTH or vehicle treatment. (**C**) Representative fluorescent images of the proximal tibia from wild type and Pth1r f/f mice following 10 days of iPTH or vehicle treatment. Red = Ebf3-CreERT2-tdTomato; green = Bglap-GFP; blue = DAPI. Scale bar = 500 µm (lower magnification) and 50 µm (higher magnification). (**D**) Quantification of tdTomato+ cells and tdTomato+GFP+ osteoblasts on metaphyseal trabecular and (**E**) endocortical bone surfaces of wild type and Pth1r f/f mice following 10 days of iPTH or vehicle treatment. (**F**) Distribution (area fraction) and (**G**) cell morphology (aspect ratio) analysis of tdTomato+ cells relative to defined distances from bone surfaces in the proximal tibia of wild type and Pth1r f/f mice following 10 days of iPTH or vehicle treatment. (**H**) Representative flow cytometry plots and analysis of non-hematopoietic Ebf3-CreERT2-tdTomato+ cells (**I**) and their skeletal stem/progenitor cell surface markers (**J**) from wild type and Sp7 f/f mice following 10 days of iPTH or vehicle treatment. (**K and L)** RT-qPCR analysis of FACS-isolated Lineage–tdTomato+ cells from wild type and Sp7 f/f mice following 10 days of iPTH and vehicle treatment. Statistical test: Two-way ANOVA with treatment and genotype (**D, E, I, and L**), or treatment and specified distance bins (**F and G**), or treatment and subpopulations (**J**) as factors followed by Šidák’s multiple comparisons test, and Student’s T-test (**K**). Data is expressed as mean ± SD, n = 4-7 mice/group.

At baseline, labeled cell numbers and osteoblast differentiation were comparable between Pth1r f/f and wild type mice across all examined surfaces **(Figures 5C–E, Supplementary Figure 12A)**. In response to iPTH, progenitor recruitment to trabecular and endocortical surfaces was similarly increased in both genotypes; however, Pth1r deletion blocked iPTH-induced osteoblast differentiation at all sites. Central marrow numbers were unaffected by genotype or treatment, and periosteal osteoblast differentiation was not observed in either group, though iPTH expanded labeled progenitors on periosteal surfaces in wild type but not Pth1r f/f mice **(Supplementary Figures 12B–D)**. Spatially, both genotypes showed comparable iPTH-induced increases in labeled cell density and elongation within 10 µm of bone; at 11–40 µm from bone surfaces, however, Pth1r deletion attenuated these effects **(Figures 5F, G)**.

Flow cytometry revealed no iPTH-induced change in overall Lineage–tdTomato+ cell numbers in either genotype **(Figures 5H, I)**. Within this population, Pth1r deletion reversed the direction of the iPTH-induced subset shift: whereas iPTH promoted BCSP contraction and 6C3+ expansion in wild type mice, Pth1r f/f mice showed increased BCSPs and decreased 6C3+ cells **(Figure 5J)**. RT-qPCR confirmed reduced Pth1r mRNA expression and attenuated iPTH-induced upregulation of *Tnfsf11* and *Vdr* in Pth1r f/f mice **(Figures 5K, L)**. Critically, iPTH-induced downregulation of the CAR cell-enforcing transcription factors *Ebf3* and *Foxc1* was abolished by Pth1r deletion.

Pth1r expression in Ebf3-lineage cells is required for iPTH-induced suppression of CAR cell-enforcing transcription factors, expansion of the 6C3+ subset, and terminal differentiation into Bglap+ osteoblasts, but is dispensable for progenitor recruitment to bone surfaces. This dissociation prompted us to consider non-cell-autonomous mechanisms of intercellular communication that govern this early step in the progenitor response to iPTH.

### PTH-stimulated recruitment of Ebf3-lineage cells to bone surfaces requires osteoclasts and is associated with TGFß-responsiveness

Given the key roles of osteoclasts in PTH-regulated bone remodeling^52^ and our focus on inter-cellular communication between CAR cells and other cell types in the bone marrow microenvironment, we performed TRAP staining to define the relationship between Ebf3-lineage cells and osteoclasts. In vehicle-treated mice, labeled cells occupied trabecular surfaces that harbored TRAP+ osteoclasts but lacked Bglap-GFP+ osteoblasts, and iPTH further enhanced this co-localization (**Figure 6A**). Quantification confirmed that labeled cells were on average closer to TRAP+ osteoclasts than to Bglap-GFP+ osteoblasts, with more cells found within a 50 µm radius of osteoclasts than osteoblasts (**Figures 6B, C**). Therefore, Ebf3-lineage progenitors preferentially associate with osteoclasts at steady state, an association amplified by iPTH.

**Figure 6.**
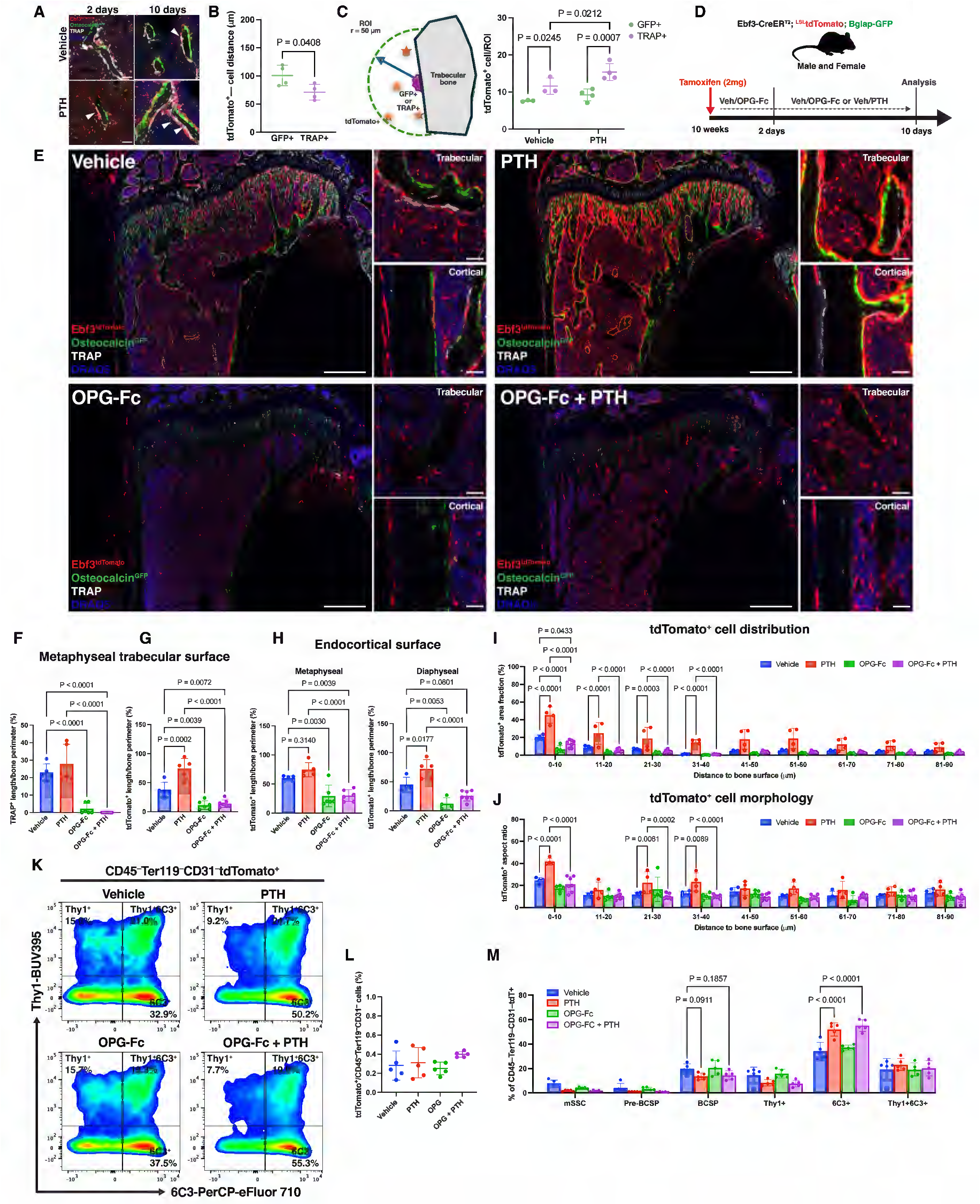
Osteoclasts are required for iPTH-stimulated recruitment of Ebf3-lineage cells to bone surfaces. **(A)** Representative fluorescent images of TRAP-stained osteoclasts in proximal tibia tissue sections from 10-week-old Ebf3-CreERT2; Rosa26-LSL-tdTomato; Bglap-GFP mice following 2 and 10 days of iPTH or vehicle treatment. White = TRAP; Red = Ebf3-CreERT2-tdTomato; green = Bglap-GFP; blue = DRAQ5. Scale bar = 50 µm. **(B)** Quantification of Ebf3-CreERT2-labeled tdTomato+ cell distance relative to Bglap-GFP+ osteoblast or TRAP+ osteoclast following 10 days of vehicle treatment. (**C**) Quantification of Ebf3-CreERT2-labeled tdTomato+ cell numbers within 50 µm radius of Bglap-GFP+ osteoblast or TRAP+ osteoclast following 10 days of iPTH or vehicle treatment. (**D**) Experimental overview to selectively deplete osteoclasts by OPG-Fc in the Ebf3-CreERT2 lineage tracing iPTH system. (**E**) Representative fluorescent images of the proximal tibia from Ebf3-CreERT2; Rosa26-LSL-tdTomato; Bglap-GFP mice following 10 days of iPTH, OPG-Fc, combination or vehicle treatment. Red = Ebf3-CreERT2-tdTomato; green = Bglap-GFP; blue = DAPI. Scale bar = 500 µm (lower magnification) and 50 µm (higher magnification). (**F**) Quantification of TRAP+ cells on metaphyseal trabecular bone surfaces, and (**G**) tdTomato+ cells on metaphyseal trabecular and (**H**) endocortical bone surfaces following 10 days of iPTH, OPG-Fc, combination or vehicle treatment. (**I**) Distribution (area fraction) and (**J**) cell morphology (aspect ratio) analysis of tdTomato+ cells relative to defined distances from bone surfaces in the proximal tibia of Ebf3-CreERT2; Rosa26-LSL-tdTomato; Bglap-GFP mice following 10 days of iPTH, OPG-Fc, combination or vehicle treatment. (**K**) Representative flow cytometry plots and analysis of (**L**) non-hematopoietic Ebf3-CreERT2-labeled tdTomato+ cells and (**M**) their skeletal stem/progenitor cell surface markers following 10 days of iPTH, OPG-Fc, combination or vehicle treatment. Statistical test: Student’s T-test (**B**), Two-way ANOVA with treatment and cell-type (**C**), or treatment and specified distance bins (**I and J**), or treatment and subpopulations (**M**) as factors followed by Šidák’s multiple comparisons test, and one-way ANOVA followed by by Šidák’s multiple comparisons test (**F, G, H, and L**). Data is expressed as mean ± SD, n = 5-6 mice/group.

To test whether osteoclasts mediate non-cell-autonomous recruitment of Ebf3-lineage cells, we depleted osteoclasts using OPG-Fc (**Figure 6D**). OPG-Fc reduced serum P1NP and CTX and prevented iPTH-induced increases in both markers, confirming effective osteoclast suppression (**Supplemental Figure 13A**). Consistent with this, OPG-Fc ablated TRAP+ osteoclasts on bone surfaces and, remarkably, reduced labeled cell numbers on trabecular and endocortical surfaces (**Figures 6E–G**). While iPTH alone again increased labeled cell recruitment, combined iPTH and OPG-Fc abolished this effect (**Figure 6H**). Spatially, OPG-Fc reduced labeled cell density within 10 µm of endocortical bone, and co-treatment with iPTH failed to restore the iPTH-stimulated increases in cell density and elongation at 11–40 µm from bone surfaces (**Figures 6I, J**).

Flow cytometry showed no difference in overall Lineage–tdTomato+ cell numbers across treatment groups, and OPG-Fc did not alter progenitor subset composition at baseline (**Figures 6K, L**). Notably, despite blocking iPTH-stimulated recruitment to bone surfaces, OPG-Fc did not impair iPTH-induced 6C3+ subset expansion. Thus, osteoclast activity is selectively required for Ebf3-lineage cell recruitment to bone surfaces, while the cell-autonomous effects of iPTH (transcriptional priming and 6C3+ expansion) proceed independently.

MicroCT of L5 vertebrae confirmed that OPG-Fc, iPTH, and combined treatment each increased trabecular bone parameters compared to vehicle (**Supplemental Figures 13B, C**), and spine histology showed iPTH-stimulated labeled cell recruitment and osteoblast differentiation at vertebral surfaces (**Supplemental Figure 13D**). Thus, Ebf3-lineage progenitors contribute to osteoblasts in response to iPTH in both the axial and appendicular skeleton.

To understand the cell-extrinsic mechanisms used by osteoclasts to promote iPTH-stimulated recruitment of Ebf3-lineage cells to bone surfaces, we performed single cell RNA-sequencing on FACS-isolated CD45–Ter119–tdTomato+ cells following 10 days of vehicle, iPTH, OPG-Fc, or OPG-Fc plus iPTH treatment (**Figure 7A)**. Unsupervised clustering again identified 6 conserved CAR subsets, 1 pre-osteoblast, and 1 mature osteoblast with similar transcriptomic profiles as above (**Figure 7B**). For each CAR and pre-osteoblast cluster, we performed cluster-specific DEG analysis comparing vehicle versus PTH, OPG-Fc, or OPG-Fc + PTH treatment (**Figure 7C)**. Notably, OPG-Fc treatment downregulated osteoblastic genes *Alpl, Spp1,* and *Col1a1* and upregulated the skeletal progenitor associated gene *Grem1*^7^. As expected, PTH-responsive module scores were higher in both PTH and OPG-Fc + PTH conditions relative to vehicle in CAR and pre-osteoblast clusters (**Figure 7D)**.

**Figure 7.**
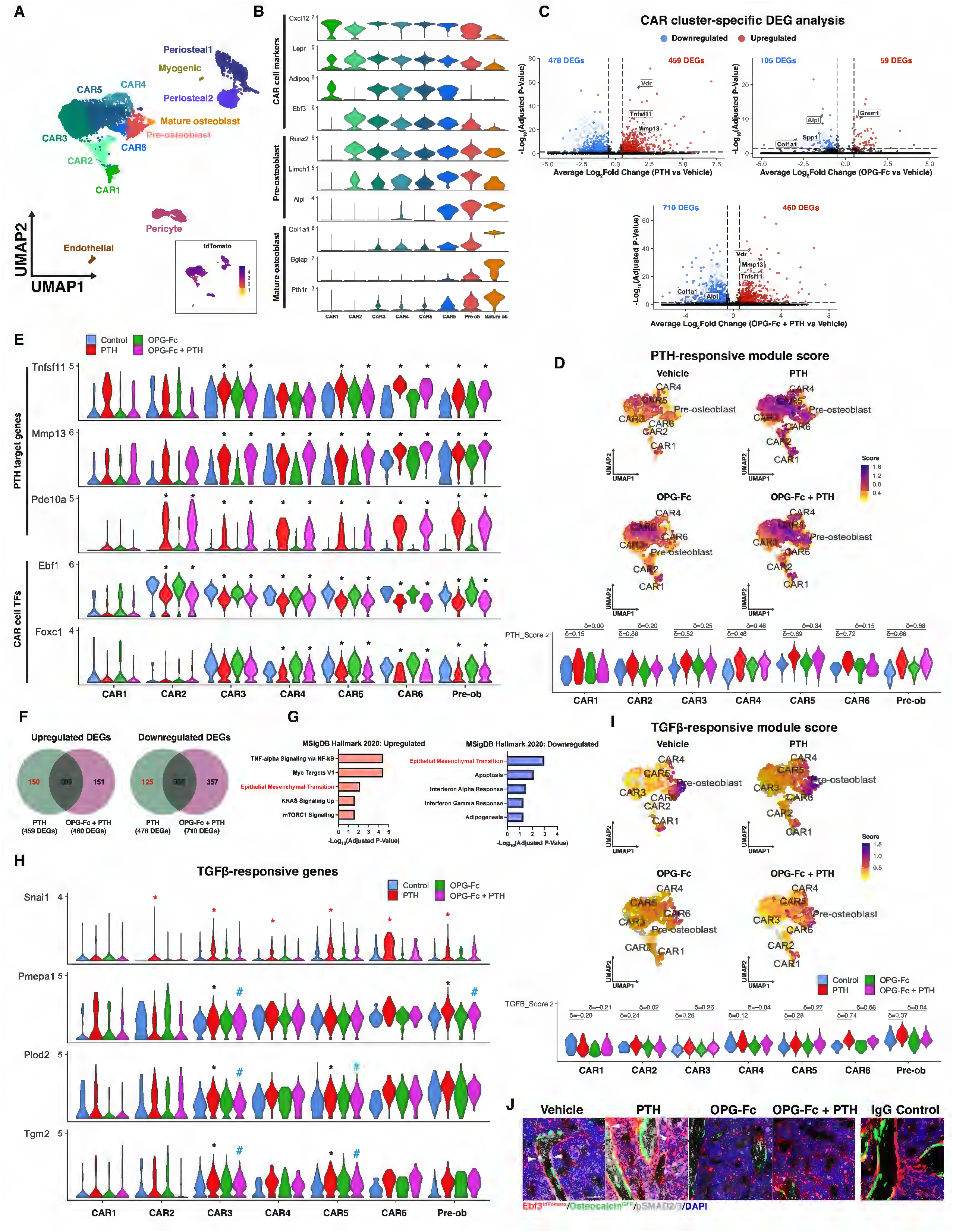
Recruitment of Ebf3-lineage cells to bone surfaces in response to iPTH is associated with TGFß-response. **(A)** Harmony-integrated UMAP of single cell RNA-sequencing data from FACS-isolated CD45–Ter119–tdTomato+ cells labeled by Ebf3-CreERT2 following 10 days of iPTH, OPG-Fc, combination or vehicle treatment (Vehicle = 1,155 cells, PTH = 1,638 cells, OPG-Fc = 1,720, and OPG-Fc + PTH = 1,773; n = 5-6 mice/group). Inset shows feature plot of tdTomato mRNA expression across clusters. (**B**) Violin plots showing mRNA expression levels of CAR cell, pre-osteoblast, and mature osteoblast marker genes in Ebf3-lineage cell clusters. (**C**) Volcano plots showing the total number of significant differentially expressed genes (DEGs) of all Ebf3-lineage CAR subclusters regulated by PTH, OPG-Fc, or OPG-Fc + PTH (adjusted P-value < 0.05 and |log2FC| > 0.5). (**D**) PTH-responsive module scores visualized by Feature plots of each Ebf3-lineage CAR cell and pre-osteoblast (top) and violin plots (bottom) across Ebf3-lineage CAR and osteoblast clusters in response to iPTH, OPG-Fc, combination, or vehicle treatment. (**E**) Violin plots showing the expression levels of canonical PTH target genes and CAR cell fate regulating transcription factor genes across Ebf3-lineage CAR and pre-osteoblast clusters in response to iPTH, OPG-Fc, combination, or vehicle treatment. (**F**) Venn-diagrams comparing differentially expressed genes altered by PTH versus OPG-Fc + PTH treatment. (**G**) Gene set enrichment analysis of all significantly up- and down-regulated DEGs by PTH but not OPG-Fc + PTH in Ebf3-lineage CAR clusters using the MSigDB Hallmark 2020 database. The top 5 significantly enriched gene sets are shown (adjusted P-value < 0.05). (**H**) Violin plots showing the expression levels of TGFß-responsive genes across Ebf3-lineage CAR and pre-osteoblast clusters in response to iPTH, OPG-Fc, combination, or vehicle treatment. (**I**) TGFß-responsive module scores visualized by Feature plots of each Ebf3-lineage CAR cell and pre-osteoblast (top) and violin plots (bottom) across Ebf3-lineage CAR and osteoblast clusters in response to iPTH, OPG-Fc, combination, or vehicle treatment. Cliff’s delta (δ) = non-parametric effect size used to measure differences between module score distributions. Differential gene expression between treatment and control cells, and treatment interactions within each cluster was assessed using the MAST hurdle model. Black asterisks (*) denotes genes that are significantly differentially expressed in current dataset (adjusted P-value < 0.05 and |log2FC| > 0.5). Red asterisks indicate genes that were significantly regulated by iPTH in the single cell RNA-sequencing dataset from Figure 3, but did not reach statistical significance in the current experiment, likely due to limited power. Blue hashtags (#) indicates a significant antagonistic treatment interaction between OPG-Fc and PTH. (**J**) Representative confocal images of immunohistochemical staining for phospho-SMAD2/3 in proximal tibia tissue sections from Ebf3-CreERT2; Rosa26-LSL-tdTomato; Bglap-GFP mice following 10 days of iPTH, OPG-Fc, combination or vehicle treatment. Red = Ebf3-CreERT2-tdTomato; green = Bglap-GFP; white = phospho-SMAD2/3; blue = DAPI. Scale bar = 50 µm.

Concurrent PTH and OPG-Fc treatment did not affect the PTH-induced expression of canonical PTH target genes (*Tnfsf11, Mmp13,* and *Pde10a*) and CAR cell transcription factors *Ebf1* and *Foxc1* (**Figure 7E)**. To identify genes that are involved with PTH-stimulated recruitment of Ebf3-lineage cells to bone surfaces, we performed GSEA on DEGs that were altered only by PTH, but not OPG-Fc + PTH (**Figure 7F**), and identified ‘Epithelial Mesenchymal Transition’ (**Figure 7G)**. Within this gene set, the expression of several downstream TGFß-responsive genes (*Snai1*^53^*, Pmepa1*^54^*, Plod2*^55,56^ and *Tgm2*^57^) was significantly increased by PTH and blunted by PTH/OPG-Fc cotreatment (**Figure 7H**). To assess whether OPG-Fc was attenuating the TGFß-responsiveness of Ebf3-lineage cells to PTH, a TGFß-responsive module score was computed using a literature-curated gene set (*Serpine1, Ccn2, Postn, Fn1, Col1a1, Col1a2, Tagln, Timp1, Vim, Itga5, Mmp2, Mmp14, Plod2, Tgm2*). In response to PTH, CAR6 and pre-osteoblast clusters showed the largest increase in TGFß-responsive module scores (**Figure 7I)**. Notably, in pre-osteoblasts, OPG-Fc cotreatment inhibited PTH-stimulated induction of the TGFß-responsive module despite having an increase in canonical PTH-target genes (**Figure 7D**). To confirm whether TGFß-responsiveness of Ebf3-lineage cells was blunted by OPG-Fc co-treatment, we stained for phospho-SMAD2/3 and observed fewer phospho-SMAD2/3+ cells **(Figure 7J)** with OPG-Fc alone or combined with iPTH treatment.

Together, these findings indicate that osteoclasts are required for the iPTH-stimulated recruitment of Ebf3-lineage cells to bone surfaces and their subsequent differentiation into mature osteoblasts. Osteoclastic bone resorption releases latent TGFß from bone matrix into the local microenvironment^58^. In contrast, early steps in PTH-induced Ebf3-lineage cell differentiation (6C3+ cell expansion, Ebf3 downregulation, reduction of adipogenic factors) occurs independently of osteoclastic bone resorption.

### Ebf3-lineage cells are important contributors to overall bone anabolism of iPTH

To assess the relative contribution of Ebf3-lineage cells to iPTH-induced bone anabolism, we returned to Ebf3-CreERT2; Sp7 f/f mice, a model that selectively blocks CAR cell maturation into osteoblasts without affecting other progenitor sources. Wild type and Sp7 f/f mice received three tamoxifen doses at 10 weeks of age, followed by vehicle or iPTH for 4 weeks (**Figure 8A**).

**Figure 8.**
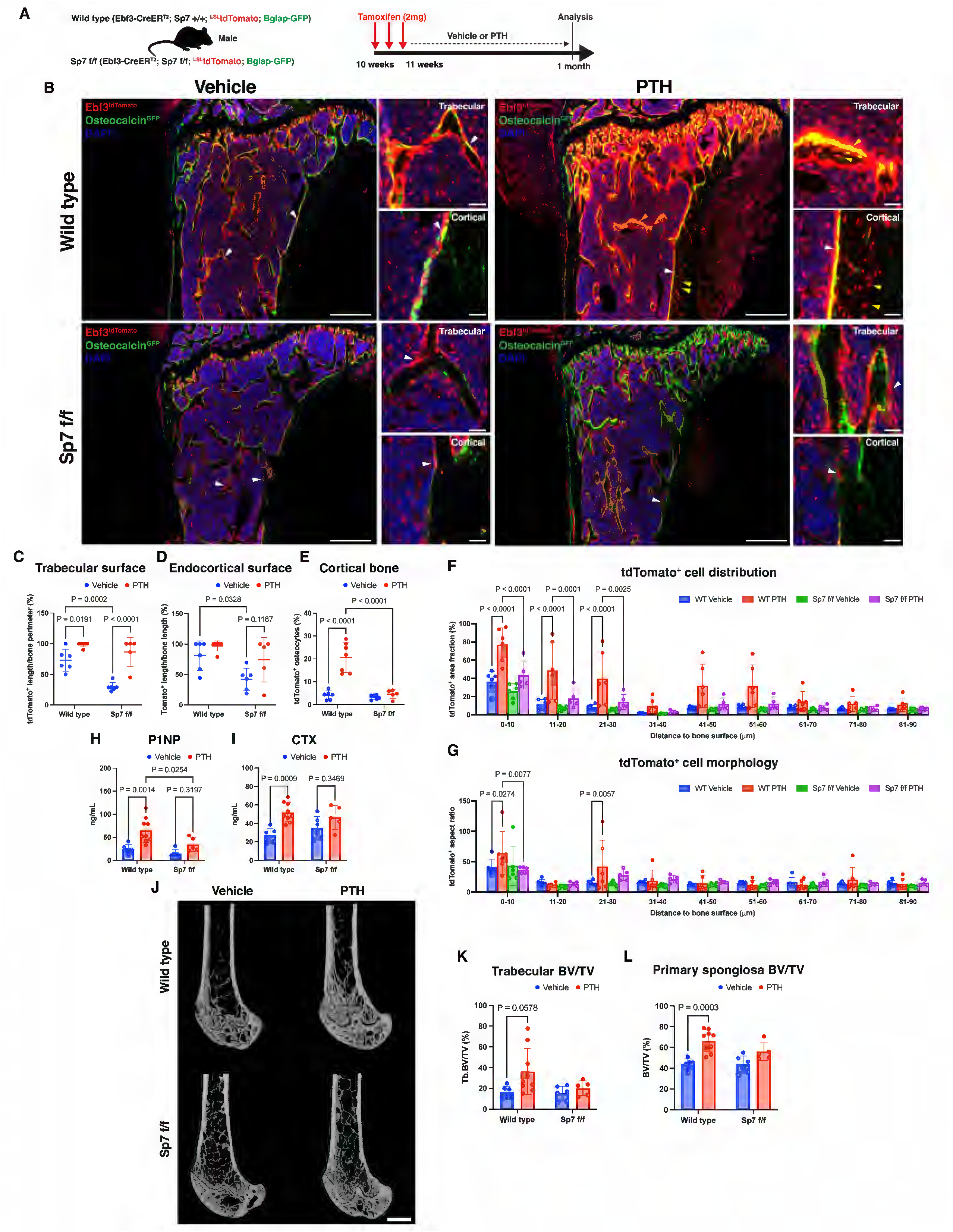
Ebf3-lineage cells are substantial contributors to the overall bone anabolism of iPTH. (**A**) Overview of lineage tracing strategy to study the relative contribution of Ebf3-lineage cells following 1 month of iPTH or vehicle treatment in male mice. (**C**) Representative fluorescent images of the proximal tibia from wild type and Sp7 f/f mice following 1 month of iPTH or vehicle treatment. Red = Ebf3-CreERT2-tdTomato; green = Bglap-GFP; blue = DAPI. White arrow: tdTomato+ cell on bone surface; yellow arrow: tdTomato+ osteocyte. Scale bar = 500 µm (lower magnification) and 50 µm (higher magnification). (**C**) Quantification of tdTomato+ cells on metaphyseal trabecular and (**D**) endocortical bone surfaces, (**E**) tdTomato+ osteocytes in the cortical bone of wild type or Sp7 f/f mice following 1 month of iPTH or vehicle treatment. (**F**) Distribution (area fraction) and (**G**) cell morphology (aspect ratio) analysis of tdTomato+ cells relative to defined distances from endocortical bone surface in the proximal tibia of wild type and Sp7 f/f mice following 1 month of iPTH or vehicle treatment. Levels of serum bone formation marker (**H**) P1NP and resorption marker (**I**) CTX from wild type and Sp7 f/f mice following 1 month of iPTH or vehicle treatment. (**J**) Representative sagittal femur microCT images and quantification of (**K**) trabecular and (**L**) primary spongiosa bone volume/total volume (BV/TV) in wild type and Sp7 f/f mice following 1 month of iPTH or vehicle treatment. Scale bar = 1 mm.Statistical test: Two-way ANOVA with treatment and genotype (**C, D, E, H, I, K and L**), or treatment and specified distance bins (**F and G**) as factors followed by Šidák’s multiple comparisons test. Data is expressed as mean ± SD, n = 6-8 mice/group.

Sp7 f/f mice showed fewer labeled cells on trabecular and endocortical surfaces at both metaphyseal and diaphyseal sites compared to wild type controls (**Figures 8B–D, Supplementary Figure 14A**). iPTH increased labeled progenitors on metaphyseal trabecular but not endocortical surfaces in Sp7 f/f mice, and Sp7 deletion blocked iPTH-induced labeled osteocyte accumulation in cortical bone (**Figure 8E, Supplementary Figure 14B**). Central marrow and periosteal cell numbers were unaffected by genotype or treatment (**Supplementary Figures 14C–F**). Spatially, the iPTH-induced increase in labeled cell density within 30 µm of endocortical bone was impaired in Sp7 f/f mice, and iPTH-stimulated cell elongation within 0–10 µm was abolished. Flow cytometry showed iPTH increased overall labeled stromal cell numbers in both genotypes; however, the iPTH-stimulated 6C3+ expansion seen in wild type mice was absent in Sp7 f/f mice (**Supplementary Figures 14G, H**).

As expected, iPTH elevated serum P1NP and CTX in wild type mice; Sp7 deletion blocked the P1NP increase and prevented a significant rise in CTX (**Figures 8H, I**). Femur microCT showed iPTH-induced gains in BV/TV in the secondary and primary spongiosa of wild type mice that were absent in Sp7 f/f mice (**Figures 8J, K**), and iPTH-stimulated increases in trabecular thickness and primary spongiosa BMD were similarly blocked (**Supplementary Figures 15A, B**). Cortical bone parameters were unaffected by genotype or treatment (**Supplementary Figures 15C, D**). Together, these findings establish Ebf3-lineage cells as major contributors to iPTH-induced bone anabolism, accounting for the bulk of trabecular bone gains at this treatment duration.

### Ebf3-lineage cells continue to participate in the anabolic action of PTH in the setting of ovariectomy

To determine whether Ebf3-lineage progenitors contribute to iPTH-induced bone anabolism under estrogen-deficient conditions, we performed OVX in Ebf3-lineage tracing mice. All CAR and osteoblast clusters express *Esr1* (estrogen receptor alpha), with highest levels in CAR6 and pre-osteoblasts (**Supplementary Figure 16A**), supporting the relevance of this model. OVX was performed at 12 weeks of age; 8 weeks later, mice received tamoxifen followed by vehicle, iPTH, OPG-Fc, or combined iPTH and OPG-Fc (**Supplementary Figure 16B**).

Consistent with findings in eugonadal mice, iPTH increased serum P1NP and CTX, OPG-Fc suppressed both markers, and all treatments increased vertebral trabecular bone mass (**Supplemental Figures 17A-C**). iPTH-stimulated recruitment of labeled cells to bone surfaces was preserved in older OVX mice, and osteoclast depletion with OPG-Fc again blocked this recruitment and prevented iPTH-induced cell morphology changes (**Supplementary Figures 16C-G**).

Flow cytometry revealed no difference in overall Lineage–tdTomato+ cell numbers across treatment groups (**Supplementary Figures 16H, I**). Within this population, however, iPTH-induced subset dynamics differed markedly from young eugonadal mice: rather than expanding the 6C3+ subset, iPTH decreased 6C3+ cell numbers in older OVX mice; OPG-Fc co-treatment also reduced 6C3+ cells, though to a lesser degree (**Supplementary Figure 16J**). Thus, while iPTH-induced labeled cell recruitment and osteoblastic differentiation are broadly shared between young eugonadal and older OVX mice, age- and/or estrogen-dependent differences may impact CAR cell subset composition and its response to iPTH.

### CAR cells in teriparatide-treated postmenopausal women show reduced expression of lineage-enforcing transcription factors and increased TGFß-responsive genes

We next investigated whether the transcriptomic changes by PTH on Ebf3-lineage cells also occur in human CAR cells. We performed single cell RNA sequencing on FACS-isolated non-hematopoietic (CD45–CD235a–CD14–) stromal cells obtained from bone marrow aspirates of postmenopausal women receiving teriparatide, a PTH analogue, for 3 months^59^. Using Harmony, we integrated these data with a single cell atlas composed of multiple recently published datasets derived from drug-naïve human bone marrow (**Figure 9A**)^60^. After unsupervised clustering on this integrated dataset, 3 CAR and 1 pre-osteoblast clusters were identified (**Figure 9B)**. CAR clusters were identified based on the highest relative expression of *CXCL12, LEPR, EBF3, FOXC1,* and *KITLG*. Pre-osteoblasts were identified based on the highest levels of *LIMCH1, SP7, RUNX2, SPP1, IBSP, COL1A1, BGLAP,* and *PTH1R* (**Figure 9C and D**).

**Figure 9.**
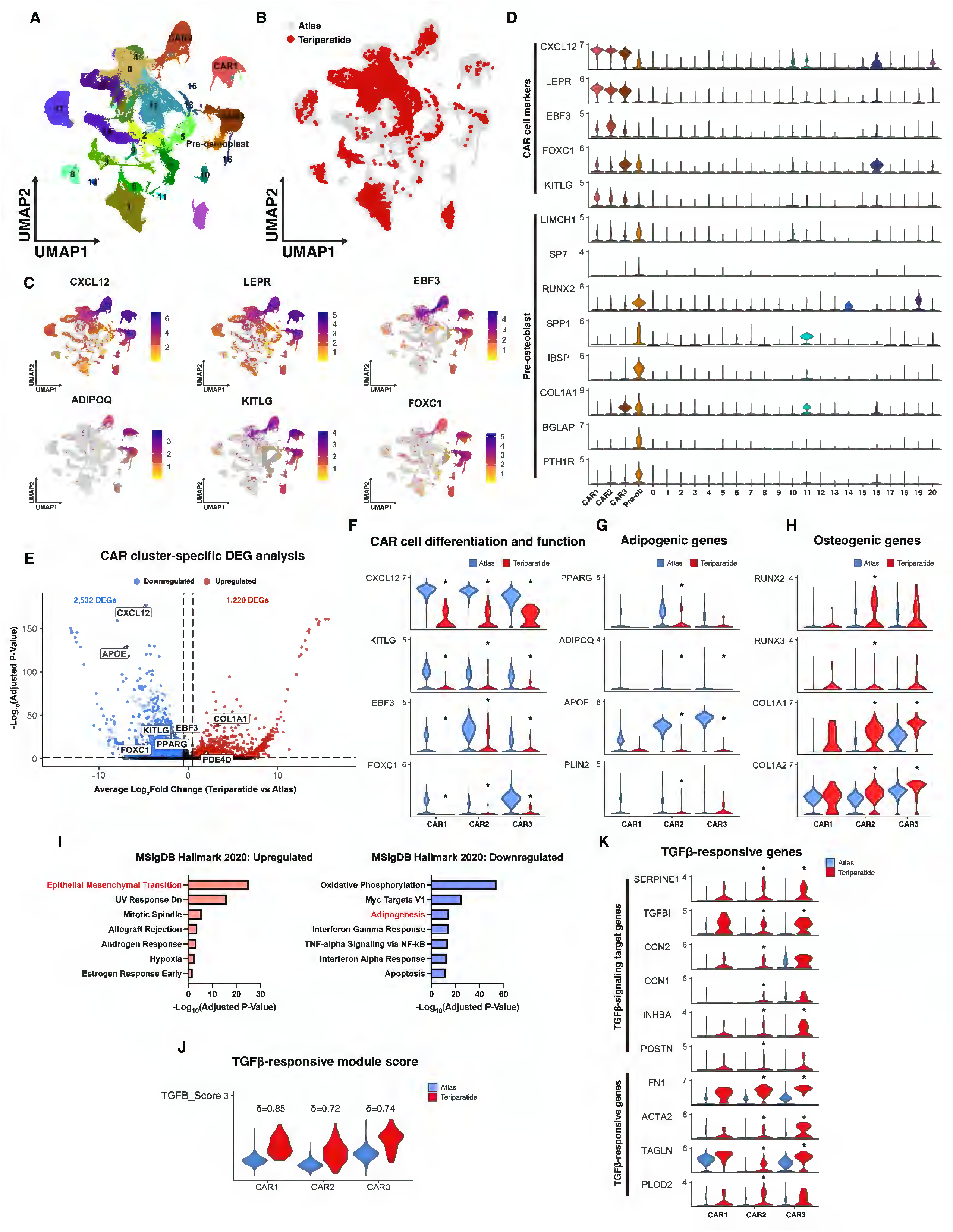
Destabilization of the CAR cell state by PTH occurs in human CAR cells. **(A)** UMAP visualization of human bone marrow single cell RNA-sequencing data following (**B**) Harmony-integration of bone marrow aspirate samples from postmenopausal women treated with teriparatide for 3 months (11,090 cells; n = 2 patient samples) and a naïve human bone marrow atlas. (**C**) Feature plots showing the expression CAR cell marker genes. (**D**) Violin plots showing mRNA expression levels of CAR cell and pre-osteoblast genes. (**E**) Volcano plot showing the total number of significant differentially expressed genes between each teriparatide-treated vs atlas CAR clusters (adjusted P-value < 0.05 and |log2FC| > 0.5). (**F**) Violin plots showing the expression levels of CAR cell genes, (**G**) adipogenic genes, (**H**) and osteogenic and bone matrix genes, across each teriparatide-treated versus atlas CAR clusters. (**I**) Gene set enrichment analysis of all up- and down-regulated differentially expressed genes identified in teriparatide-treated CAR clusters using the MSigDB Hallmark 2020 database. (**J**) Violin plots of TGFß-responsive module scores of teriparatide-treated versus atlas CAR clusters. Cliff’s delta (δ) = non-parametric effect size used to measure differences between module score distributions. (K) Violin plots showing the expression levels of TGFß-responsive genes between teriparatide-treated vs atlas CAR clusters. Differential gene expression between teriparatide-treated and atlas cells within each CAR cluster was assessed using the MAST hurdle model. Asterisk (*) denotes significance (adjusted P-value < 0.05 and |log2FC| > 0.5).

To assess the transcriptomic effects of teriparatide treatment in CAR cells, we performed cluster-specific DEG analysis (teriparatide versus atlas) in each CAR cluster (**Figure 9E**). Like the effects of iPTH in Ebf3 lineage cells in mice, teriparatide treatment in women led to marked decreases in the expression of hematopoiesis supportive genes *KITLG* and *CXCL12*, and CAR cell-enforcing transcription factors *EBF3* and *FOXC1* (**Figure 9F**). Furthermore, the expression of adipogenesis-related genes such as *PPARG, ADIPOQ, APOE,* and *PLIN2* were reduced **(Figure 9G)** while osteogenic genes *RUNX2, RUNX3, COL1A1,* and *COL1A2* were increased in teriparatide-treated CAR cells (**Figure 9H)**. GSEA on all the teriparatide-upregulated DEGs identified the ‘Epithelial Mesenchymal Transition’ gene set as most enriched **(Figure 9I)**. Within this gene set, several canonical TGFß-signaling target genes such as *SERPINE1, TGFBI, CCN2, CCN1, INHBA, POSTN,* and TGFß-responsive genes *FN1, ACTA2, TAGLN, and PLOD2* were increased in teriparatide-treated CAR cells **(Figure 9J)**. Consistent with this finding, increased TGFß-responsive module scores were noted in teriparatide-treated CAR cells versus naïve atlas populations (**Figure 9K)**.

Taken together, these results establish CAR cells as a critical cellular target of iPTH. iPTH promotes osteoblastic differentiation of CAR cells through coupled cell-intrinsic and cell-extrinsic mechanisms. PTH directly acts on CAR cells to suppress the expression of critical transcription factors responsible for suppressing osteogenic differentiation, thereby priming these cells for osteogenic differentiation. Cell-extrinsically, PTH stimulates osteoclastic bone resorption, likely driven by RANKL induction by mature osteoblasts and osteocytes, resulting in enhanced TGFß signaling that may promote recruitment of these primed progenitors to bone surfaces. Once localized to bone surfaces, PTH again acts directly on CAR cell-derived pre-osteoblasts to drive their maturation into osteoblasts and osteocytes **(Figure 10).**

**Figure 10.**
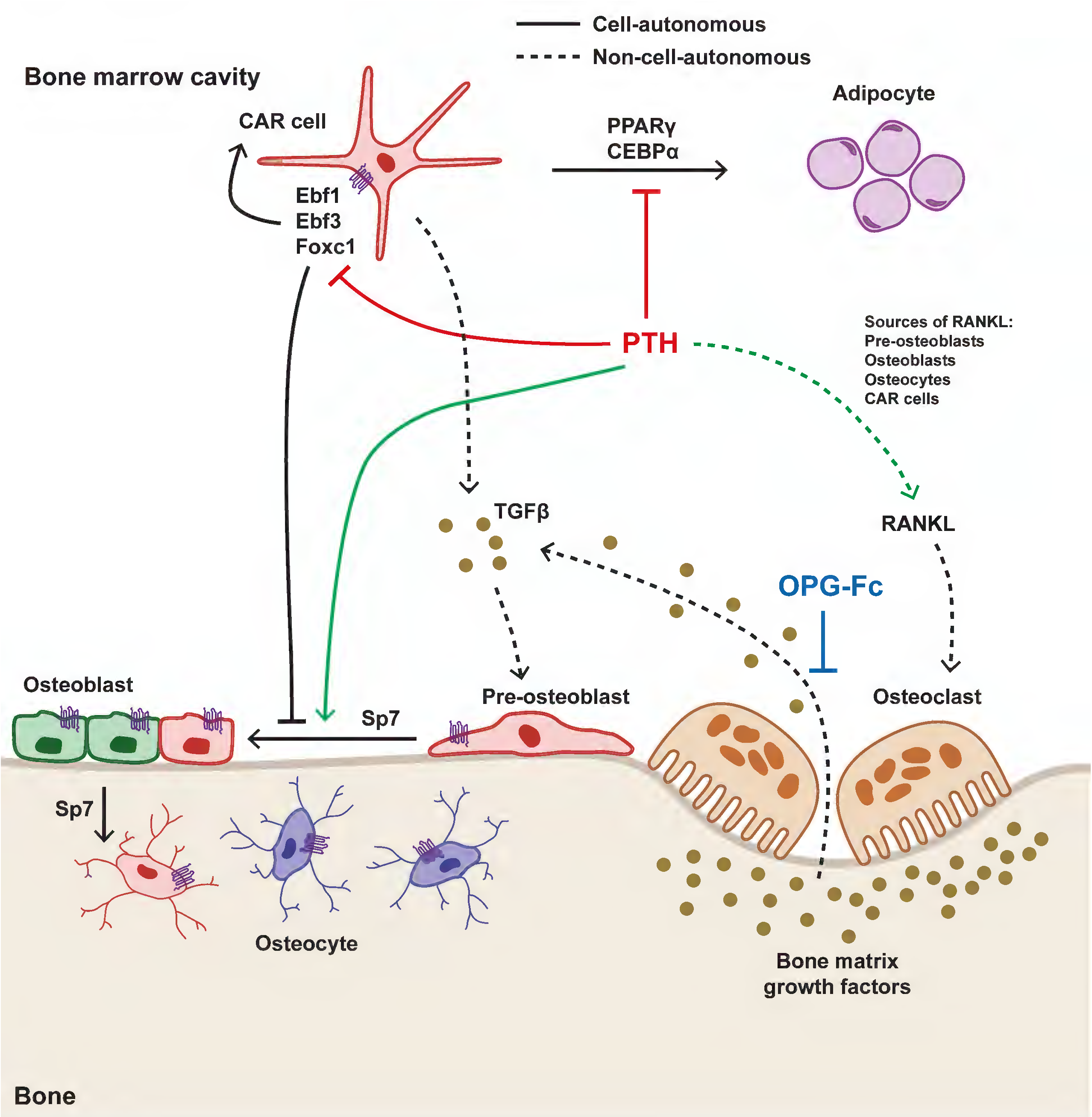
Model of CAR cell autonomous and non-cell-autonomous mechanisms by which iPTH promotes CAR cell-mediated osteoblastogenesis. CAR cells express the transcription factors Ebf1, Ebf3, and Foxc1, which maintain their hematopoiesis-supportive phenotype. iPTH directly acts on CAR cells via Pth1r to suppress CAR cell-maintaining transcriptional programs, thereby priming osteoblastic differentiation and inhibiting adipogenic fate (cell-autonomous effect). In parallel, iPTH stimulates osteoclastic bone resorption, through induction of RANKL in multiple cell types including mature osteoblasts, osteocytes, and CAR cells. Bone resorption releases matrix-derived growth factors, such as TGFβ, which recruit CAR cells to bone surfaces (non-cell-autonomous effect). Once localized to bone surfaces, iPTH again directly targets CAR cell-derived pre-osteoblasts via Pth1r to promote Sp7-dependent maturation into osteoblasts and terminally differentiated, matrix-embedded osteocytes (cell-autonomous effect). Solid lines indicate cell-autonomous mechanisms used by PTH to drive Ebf3 lineage cell osteoblast differentiation, and dashed lines indicate non-cell-autonomous mechanisms employed by PTH to recruit CAR cells to bone surfaces.

## Discussion

This study identifies Ebf3-lineage CAR cells as major cellular contributor in the bone anabolic actions of iPTH at skeletal maturity. In contrast to growth-associated progenitors residing near the growth plate, Ebf3-lineage cells are distributed throughout bone and can give rise to osteoblasts on trabecular and endocortical surfaces. Our studies demonstrate that iPTH promotes the osteoblastic differentiation of CAR cells. As early response to PTH, the CAR cell phenotype is destabilized in a cell-autonomous manner through loss of the expression of crucial CAR-cell maintaining transcription factors *Ebf3*, *Ebf1*, and *Foxc1*, thereby permitting their osteogenic-commitment while reducing adipogenesis. iPTH also acts directly on Ebf3-lineage CAR cells to expand the 6C3+ CAR cell subpopulation to maintain the bone marrow niche. A key event in PTH-induced osteoblastic differentiation of CAR cells involves their recruitment to bone surfaces and morphologic change from a reticular to more elongated shape, a process that requires osteoclasts and may be mediated by TGFß signaling. Preventing Ebf3-lineage CAR cells from contributing to the osteoblast pool via Sp7 deletion severely blunts the PTH-induced gain in trabecular bone mass. Critically, the gene regulatory network controlled by iPTH in murine Ebf3-lineage CAR cells is conserved in human CAR cells from teriparatide-treated patients.

### CAR cell-autonomous effects of PTH

Pth1r mediates the cell-autonomous effects of iPTH in Ebf3-lineage cells, driving 6C3+ CAR cell expansion, suppressing CAR cell-maintaining transcriptional programs, and promoting osteoblastic differentiation, but is dispensable for their recruitment onto bone surfaces. The long-term consequences of Pth1r deficiency in Ebf3-lineage on the bone marrow microenvironment, both at steady state and in response to iPTH, remain to be determined. CAR cells play a crucial role in supporting hematopoiesis. Deletion of Pth1r in Lepr-expressing cells and their progeny appeared to have only a modest effect on hematopoiesis^61^; however, the efficiency of Cre-dependent recombination of the Pth1r floxed allele in these cells was not assessed. Pth1r ablation in mesenchymal progenitors using Prx1-Cre^62^ resulted in a marked expansion of bone marrow adipocytes and increased bone resorption^8^. Given that iPTH-induced osteoblastic differentiation of CAR cells is blocked in the absence of Pth1r, whether this alters their adipogenic commitment or long-term hematopoietic supportive function following long-term iPTH administration warrants further study.

### CAR cell-extrinsic effects of PTH

PTH stimulates osteoclastic bone resorption, leading to enhanced TGFß signaling which may promote the recruitment of CAR cells to bone surfaces. This bone resorptive effect is likely driven by RANKL induction by osteoblasts and osteocytes, which are sources that are spatially localized at bone surfaces and therefore optimally positioned to promote osteoclast activity. CAR cells have been shown to be major producers of RANKL for osteoclast development^24,63,64^, and our data indicate that PTH also directly induces RANKL expression in CAR cells in a Pth1r-dependent manner (**Figure 5L**).

However, ablation of Pth1r in CAR cells did not impair their recruitment to bone surfaces in response to PTH, suggesting that CAR cell-derived RANKL is not the predominant driver of this osteoclast-dependent recruitment process. Bone matrix is abundant in growth factors including TGFß, IGFs, BMPs, WNTs which can be released during osteoclastic resorption^65^. Out of these factors, TGFß1 has been implicated as a chemoattractant involved in mesenchymal stromal progenitor cell recruitment to bone surfaces during physiologic and stimulated bone remodeling^21,66,67^. Defining specific TGFß ligand and receptor isoforms responsible for osteogenic CAR cell recruitment to bone surfaces represents an important priority for future research. Furthermore, whether this migratory process is a mediated directly by osteoclasts or indirectly through their bone resorptive activity requires further investigation. Our data suggests that modulating the TGFß-responsiveness of CAR cells to enhance their recruitment to bone surfaces may be a potential therapeutic strategy to promote the recruitment of these critical osteoblast progenitors to bone surfaces.

### Relative contribution of CAR cells to the bone anabolism of iPTH

CAR cells, and not growth plate-associated progenitors, are the major osteoblast progenitor population that contributes to the bone anabolic actions of iPTH. Despite blocking the CAR cell contribution to the osteoblast pool via Sp7 deletion with Ebf3-CreERT2, we note that Bglap-GFP+ osteoblasts persist on trabecular and endocortical bone surfaces and are increased with iPTH treatment (**Figure 8B),** thus indicating that other sources of osteoblast progenitors are involved. Quiescent bone lining cells can be stimulated by iPTH to become active osteoblasts^68^. However, from these studies it appears that CAR cell-derived osteoblasts are critical to iPTH-induced bone gains at this treatment duration **(Figure 8H-L)**.

### Functional heterogeneity among CAR cells

At present, we do not know why some CAR cells in the central marrow remain unrecruited to bone surfaces in response to PTH. CAR cell proliferation is low (approximately 5% EdU+; **Supplementary Figure 6B**), consistent with previous reports^5,11^, and is unaffected by iPTH. Through single cell RNA-sequencing analysis of Ebf3-lineage cells, we identified transcriptomic heterogeneity amongst Ebf3-lineage CAR cells, despite all subsets expressing high levels of *Cxcl12* and other canonical markers such as *Lepr* and *Ebf3*. The most notable contrast is between CAR1 and CAR6 which have transcriptomic profiles that are consistent with previous reports of an adipogenic and osteogenic-primed subsets, respectively^28,29,41^. Even though all subsets exhibit PTH-induced increases in target genes *Tnfsf11, Mmp13,* and *Vdr*, CAR1 showed the lowest increase in PTH-response module score and no changes in adipogenic genes (*Adipoq* and *Apoe*) compared to all other CAR subsets (**Figure 3)**. Thus, it may be that CAR1 cells represents a CAR subset that is not recruited to bone surfaces following iPTH. In addition, CAR1 is the least enriched in the expression of actin cytoskeleton remodeling or migration-related genes (*Dock1, Trio, Arhgap35* (**Supplementary Figure 8C**)), further supporting the notion that this CAR subset may not be migratory. Unfortunately, differential gene expression analysis between the 6 CAR subsets did not reveal specific/exclusive single markers that can be used to identify spatially where these CAR subsets reside in the marrow. Further investigation using spatial transcriptomic approaches may shed additional light onto anatomic and functional differences between CAR cell subsets.

### Clinical data for OPG-Fc + PTH

Clinical data from post-menopausal women showed that concomitant intermittent teriparatide and denosumab (akin to OPG-Fc) therapy is more effective at increasing bone mineral density than either medication alone^69^. Our *in vivo* results are consistent with this **(Supplementary Figure 13 and 17)**. iPTH acts to stimulate both modeling-(osteoclast-independent) and remodeling-based (osteoclast-dependent) bone formation, whereas OPG-Fc inhibits osteoclastic bone resorption. Therefore, changes in osteoblastic activity observed with OPG-Fc plus iPTH combination treatment likely reflect modeling-based bone formation. Clinical data showing that serum osteocalcin was less suppressed in combination group compared to denosumab monotherapy^69^ support this notion. Notably, we observe that some tdTomato+ cells persist on endocortical surfaces at the mid-diaphysis with OPG-Fc treatment and that tdTomato+ cell numbers increase modestly with concurrent iPTH treatment compared to OPG-Fc alone (**Figure 6H and Supplementary Figure 16E)**. Future studies are needed to address the mechanisms dictating how CAR cells are recruited to bone surfaces that are undergoing modeling versus remodeling.

### Future Directions

This present study focuses primarily on the bone anabolic actions of iPTH on CAR cell dynamics, given its translational relevance to understanding the mechanisms of PTH-based osteoporosis therapy. The response of CAR cells to continuous (rather than intermittent) hyperparathyroidism remains to be determined. The transcription factors *Runx1* and *Runx2* play crucial regulatory roles in preventing the fibrotic conversion of CAR cells^70^. The CAR cell-intrinsic or extrinsic mechanism underlying this phenomenon are unclear. Given that bone marrow fibrosis occurs in the setting of chronic hyperparathyroidism and the potential fibrotic fate of CAR cells, the role of CAR cells in this disease warrants further investigation.

Ebf3-CreERT2 also marks a PTH-responsive subset of periosteal cells (**Figure 3F)**. Periosteal Ebf3-lineage cells do not differentiate into Bglap+ osteoblasts during the length (2 months) of tracing these cells at baseline or in response to iPTH. Osteocytes descended from Ebf3-lineage cells are preferentially localized on the endosteal half of cortical bone rather than periosteal side, further supporting the notion that Ebf3-lineage periosteal cells are not osteogenic-committed. The role of these periosteal Ebf3-expressing cells remains to be determined. Ebf3-lineage periosteal cells were identified based on expression of *Prg4, Mfap5,* and *Clec3b,* the latter two identify periosteal cells^28^. Given the established role of the periosteum in bone repair and regeneration, whether these periosteal cells respond to perturbations, such as bone fracture or mechanical stress, warrants future investigation.

### Study Limitations

First, although PTH acts directly on CAR cells to suppress expression of critical transcription factors (Ebf3, Ebf1, and Foxc1), the specific signaling mechanisms that mediate these gene expression changes^71^ are not known. Second, we employed two different lineage tracing systems, Ebf3- and Sox9-CreERT2, to study the effects of iPTH on two major osteoblast progenitor populations. In comparable tamoxifen-pulse chase time points in 10-week-old mice, we observed that Ebf3-CreERT2 labeled cells more efficiently gave rise to osteoblasts (**Figure 1B**) than Sox9-CreERT2 (**Supplementary Figure 11F**) after 10 days of iPTH. However, we were unable to directly assess the relative contribution between these two osteoblast progenitor populations in the same animal due to lack of appropriate dual recombinase-driven lineage tracing systems.

Third, estrogen deficiency (induced by ovariectomy) led to a reduction of the 6C3+ subpopulation within Ebf3-lineage cells following iPTH treatment. This finding lies in contrast to our results in younger/eugonadal mice in which iPTH expands this subpopulation. The 6C3+ subpopulation represents the parental CAR cell population (Cxcl12-GFP^+^Ebf3-tdTomato^+^; **Supplementary Figure 5 and 8)**; this finding suggests that the CAR cell pool may become depleted over time with prolonged iPTH treatment. If true, this may be a potential mechanism that could explain why the bone anabolic efficacy of iPTH-based therapies slowly wane over time^72,73^. To rigorously test this hypothesis, future experiments controlled with sham-operated animals should be included to examine the effects of ovariectomy on CAR cell dynamics. Finally, due to the clinical study (DATA-Biopsy) design in which marrow aspirates were collected from teriparatide-treated post-menopausal women with osteoporosis, a control (placebo) treatment group was not included. Thus, we employed a reference “atlas” dataset of similarly-processed marrow cells as an aggregate untreated control population. Future prospective studies are needed to examine the effects of teriparatide (versus baseline or no treatment) on human CAR cells over time.

### Summary

We establish that CAR cells are a critical subset of osteoblast progenitors that drive iPTH-induced bone anabolism through both cell-autonomous and non-autonomous mechanisms. We report key transcriptional mechanisms through which iPTH directs murine and human CAR cell fate, suppressing adipogenic programming in favor of osteogenic commitment. We demonstrate that osteoclastic bone resorption is required for iPTH-stimulated recruitment of CAR cells to bone surfaces, suggesting that TGFß-responsiveness may have an important role in this process. Together, these findings define a unified mechanism of bone anabolic PTH action in which intrinsic transcriptional priming of CAR cells towards osteogenic fate converges with osteoclast-mediated ‘niche activation’ to drive osteoblast differentiation. Targeting CAR cell TGFß-responsiveness may represent a therapeutic strategy to enhance recruitment of this critical class of osteoblast progenitors to bone surfaces and improve the efficacy of anabolic treatment for osteoporosis.

## Figure Legends

**Supplementary Figure 1.**
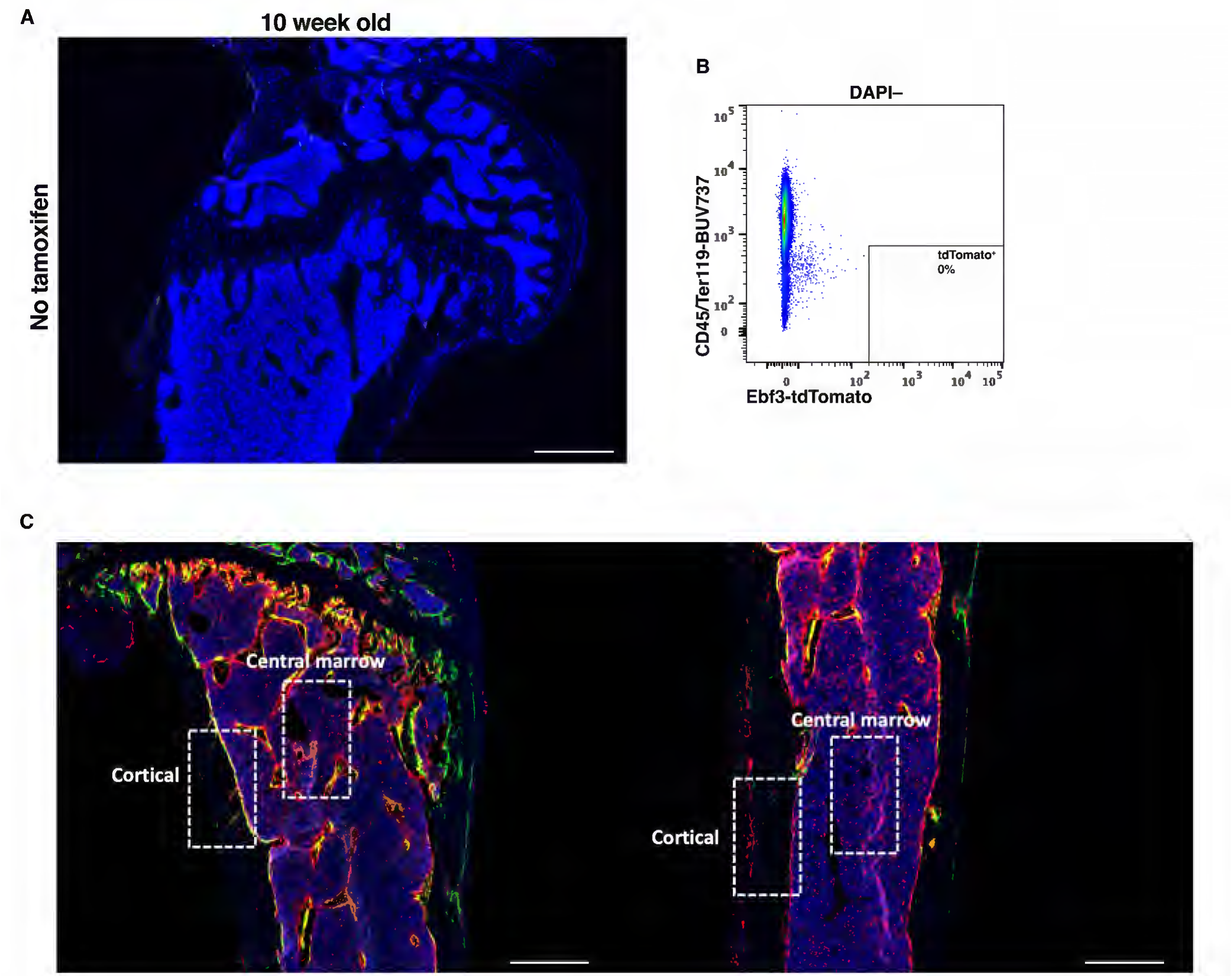
(**A)** Representative fluorescent images of the humerus and (**B**) flow cytometry analysis shows tdTomato+ cells are not present in 10-week-old Ebf3-CreERT2; Rosa26-LSL-tdTomato mice when tamoxifen is not administered. **(C)** Region of interest (500 µm x 750 µm box; 400 µm below the growth plate) used to quantify tdTomato+ cells in the central marrow and cortical bone at the metaphyseal and diaphyseal region of the tibia. Scale bar = 500 µm.

**Supplementary Figure 2.**
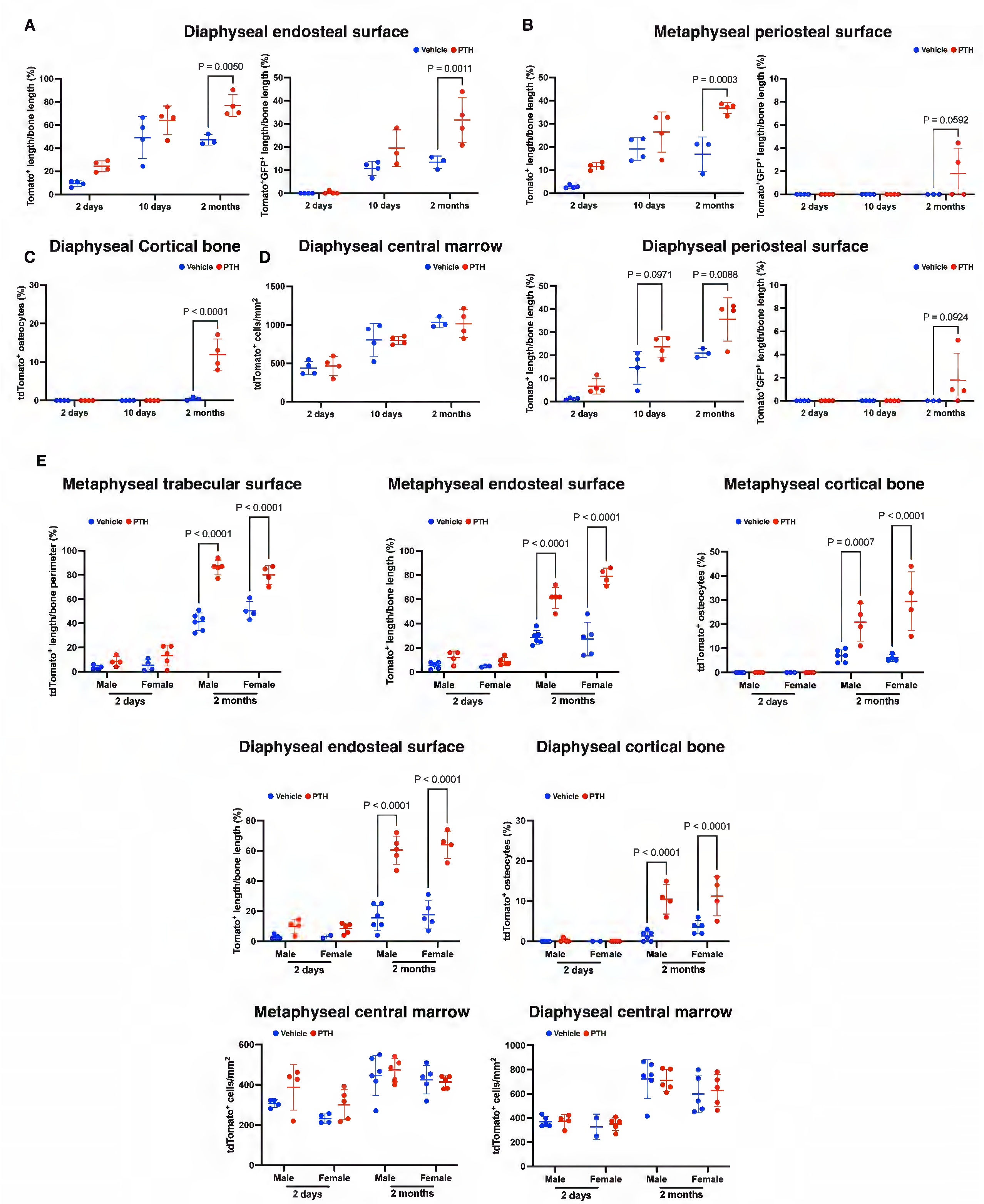
Fluorescent image quantification of Ebf3-CreERT2-labeled tdTomato+ and tdTomato+Bglap-GFP+ cells in the (**A**) diaphyseal endocortical, (**B)** metaphyseal and diaphyseal periosteal, (**C**) diaphyseal cortical bone, and (**D**) diaphyseal central marrow skeletal regions following 2 days, 10 days, and 2 months of iPTH or vehicle treatment. **(E)** Fluorescent image quantification of Ebf3-CreERT2-labeled tdTomato+ cells from male and female mice at distinct skeletal sites of the humerus following 2 days and 2 months of iPTH or vehicle treatment. Statistical test: Two-way ANOVA with treatment and time (**A-E**) as factors followed by Šidák’s multiple comparisons test. Data is expressed as mean ± SD, n = 3-6 mice/group.

**Supplementary Figure 3.**
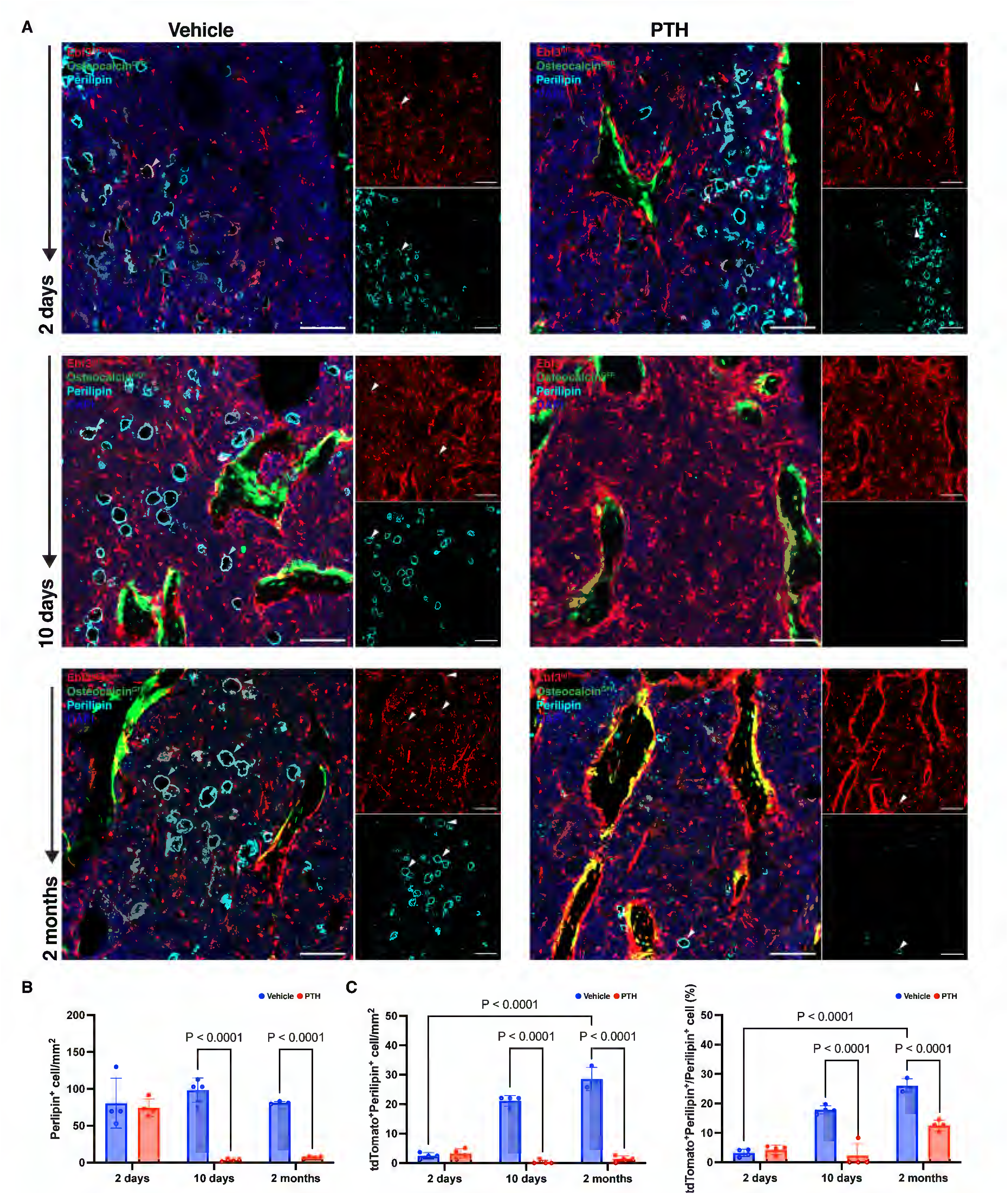
(**A**) Representative confocal images of immunohistochemical staining for perilipin in proximal metaphysis tibia tissue sections from Ebf3-CreERT2; Rosa26-LSL-tdTomato; Bglap-GFP mice following 2 days, 10 days, and 2 months of iPTH or vehicle treatment. Cyan = Perilipin; Red = Ebf3-CreERT2-tdTomato; green = Bglap-GFP. Scale bar = 100 µm. (**B**) Quantification of Perilipin+ cells, (**C**) tdTomato+Perilipin+ cells, (**D**) tdTomato+Perilipin+ cells relative to total Perilipin+ cells were performed on confocal images (640 µm x 640 µm) acquired at the proximal metaphysis (500 µm below the growth plate) of the tibia. Statistical test: Two-way ANOVA with treatment and time (**B and C**) as factors followed by Šidák’s multiple comparisons test. Data is expressed as mean ± SD, n = 3-4 mice/group.

**Supplementary Figure 4.**
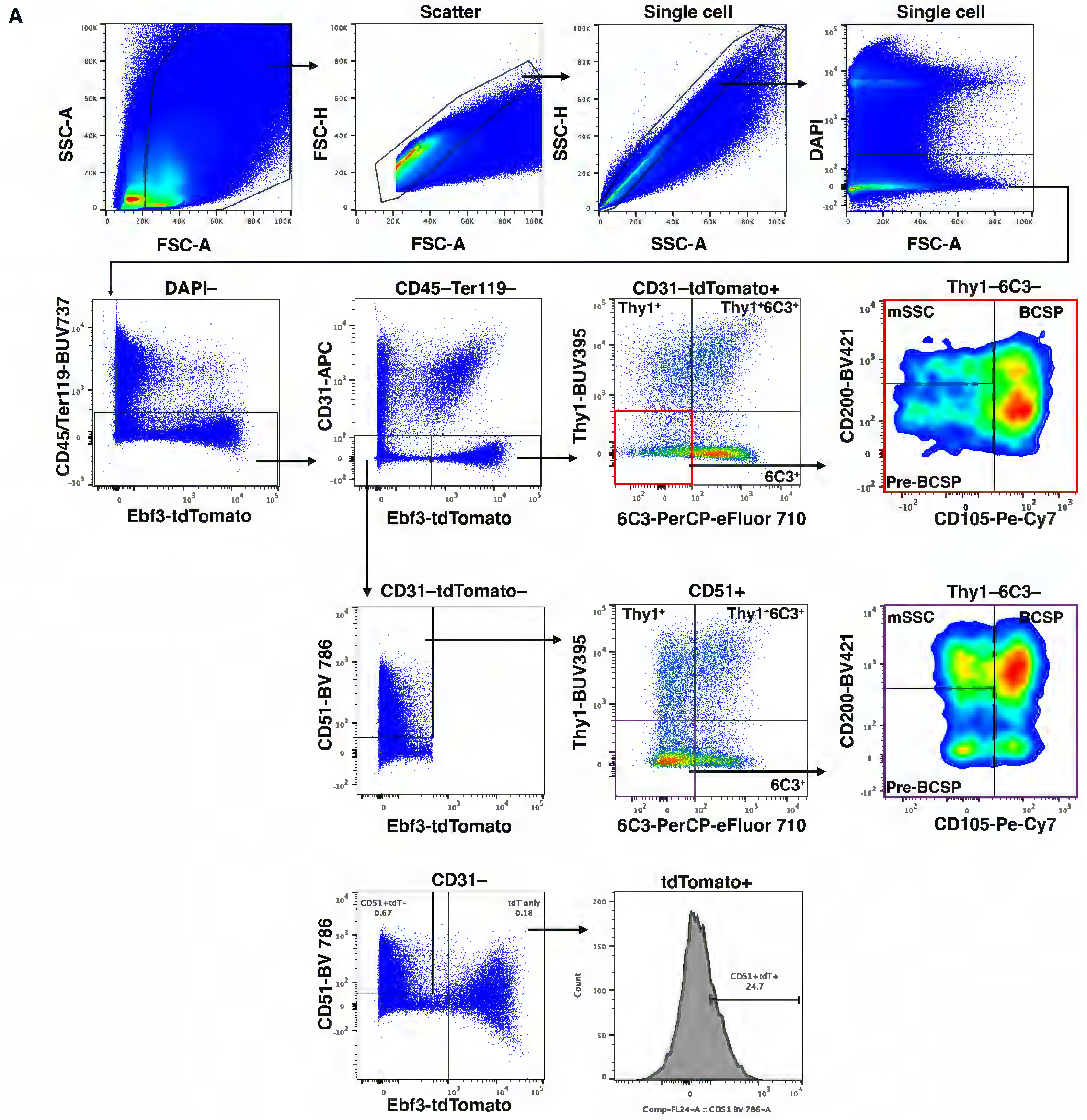
(**A**) Multicolor flow cytometry gating strategy used to analyze non-hematopoietic Ebf3-lineage tdTomato+ and tdTomato– skeletal stem and progenitor subsets. mSSC: Mouse skeletal stem cell; Pre-BCSP: Pre-bone/cartilage/stromal progenitor; BCSP: Bone/cartilage/stromal progenitor.

**Supplementary Figure 5.**
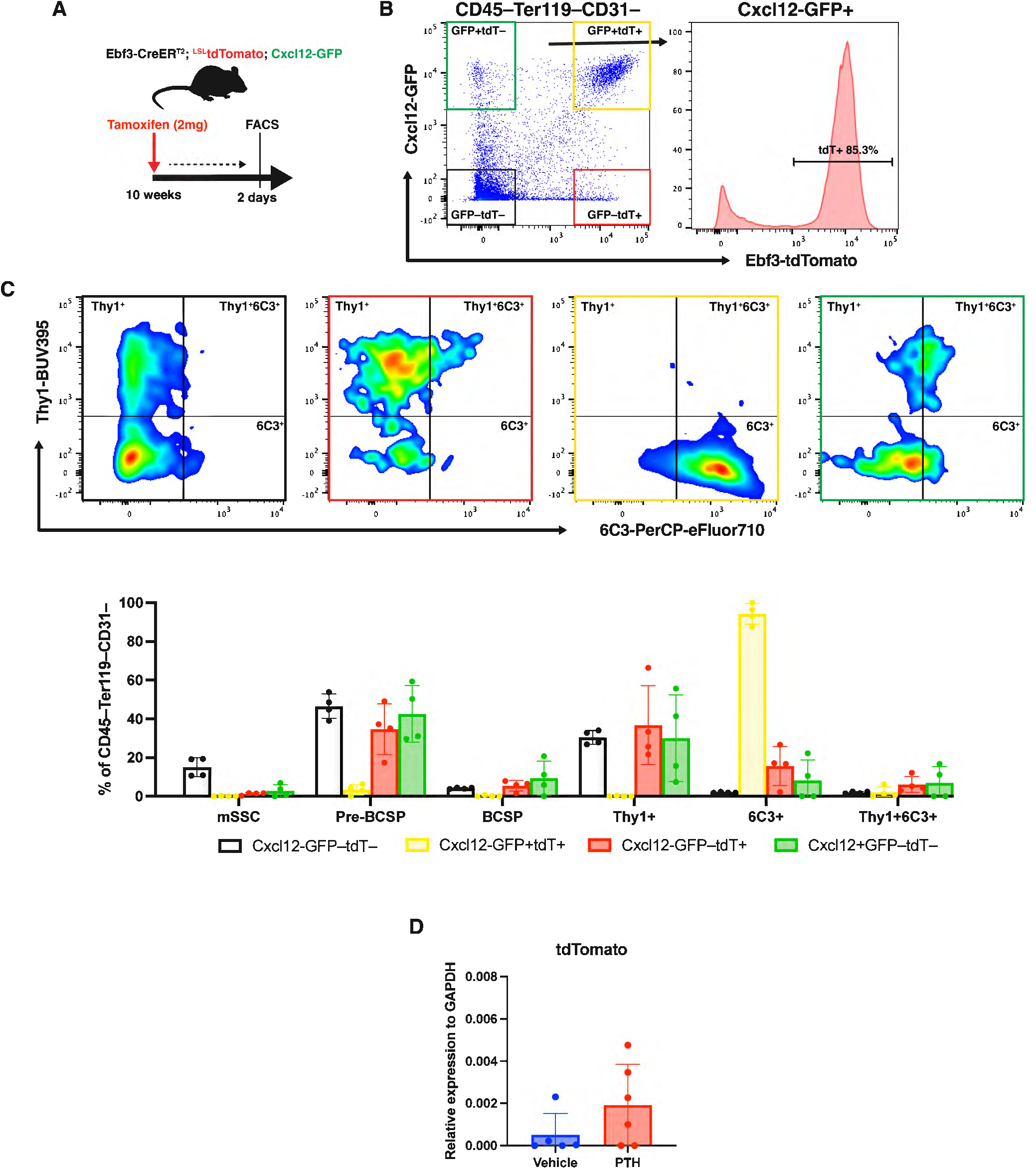
(**A**) Experimental overview for labelling parental CAR cells using Ebf3-CreERT2; Rosa26-LSL-tdTomato; Cxcl12-GFP mice treated with tamoxifen at 10-weeks-old and sacrificed 2 days later. (**B**) Flow cytometry analysis showing Ebf3-CreERT2 labels approximately 85% of Cxcl12-GFP+ cells in enzymatically digested marrow plus bone fragments. (**C**) Flow cytometry analysis indicates parental CAR (tdTomato+Cxcl12-GFP+) cells predominately lie within the 6C3+ stromal progenitor subset. (**D**) RT-qPCR analysis of tdTomato mRNA of the remaining bone fragments post-serial digestion of tibia and femur from Ebf3-CreERT2; Rosa26-LSL-tdTomato mice following 2 weeks of iPTH or vehicle treatment. Statistical test: Student’s T-test (**D**). Data is expressed as mean ± SD, n = 5-6 mice/group.

**Supplementary Figure 6.**
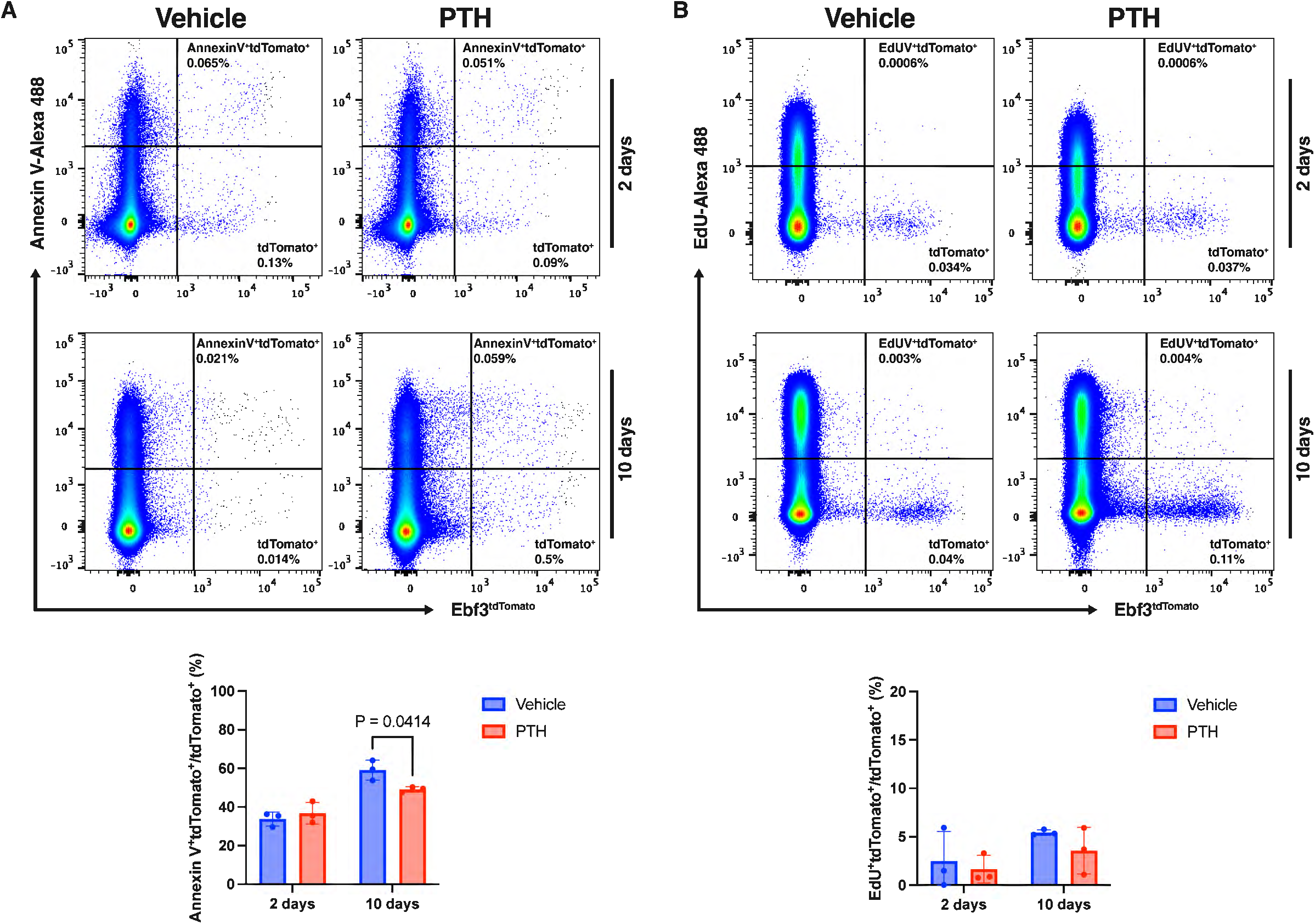
Flow cytometry analysis for (**A**) Annexin V (apoptosis) and (**B**) EdU (proliferation) staining of Ebf3-CreERT2-tdTomato+ cells labeled at 10 weeks of age, following 2 and 10 days of iPTH or vehicle treatment.

**Supplementary Figure 7.**
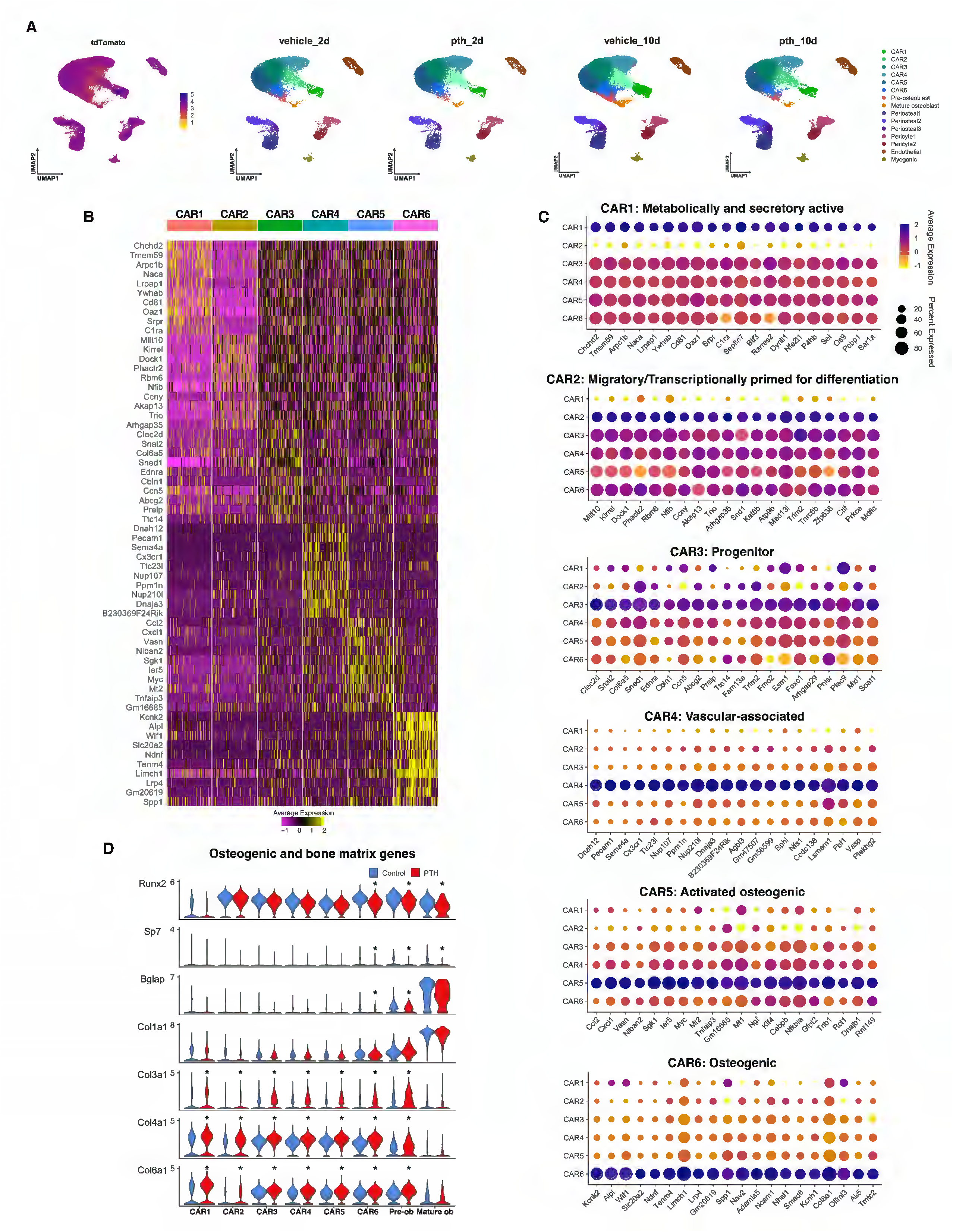
(**A**) Feature plot of tdTomato mRNA expression of Ebf3-lineage cell clusters following Harmony-integration of libraries from 2 and 10 days of iPTH or vehicle treatment. (**B**) Heatmap of the top 10 differentially expressed genes of each CAR cluster that were identified from unsupervised clustering. (**C**) Dot plots of the top 20 differentially expressed genes of each CAR cluster relative to all other CAR clusters. (**D**) Violin plots showing the expression levels of osteogenic and bone matrix genes of each CAR and osteoblast clusters with PTH versus vehicle treatment.

**Supplementary Figure 8.**
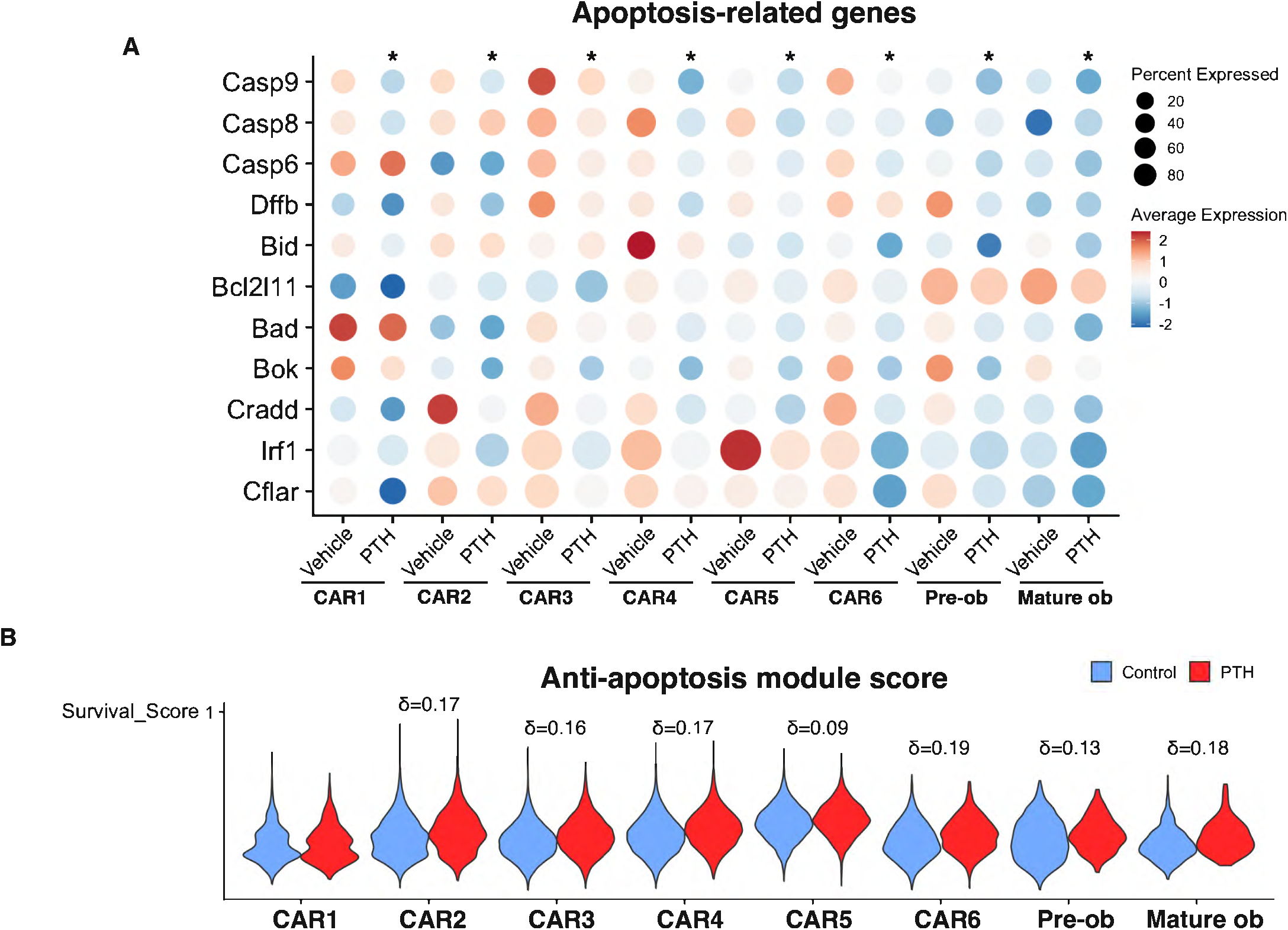
Single cell RNA-sequencing analysis of (**A**) canonical apoptosis-related genes regulated by PTH across Ebf3-lineage CAR and osteoblast clusters. (**B)** Anti-apoptosis and survival gene module scoring of Ebf3-lineage CAR and osteoblast clusters in response to PTH or vehicle treatment. Asterisk (*) denotes significance (adjusted P-value < 0.05 and |log2FC| > 0.5). Cliff’s delta (δ) = non-parametric effect size used to measure differences between module score distributions.

**Supplementary Figure 9.**
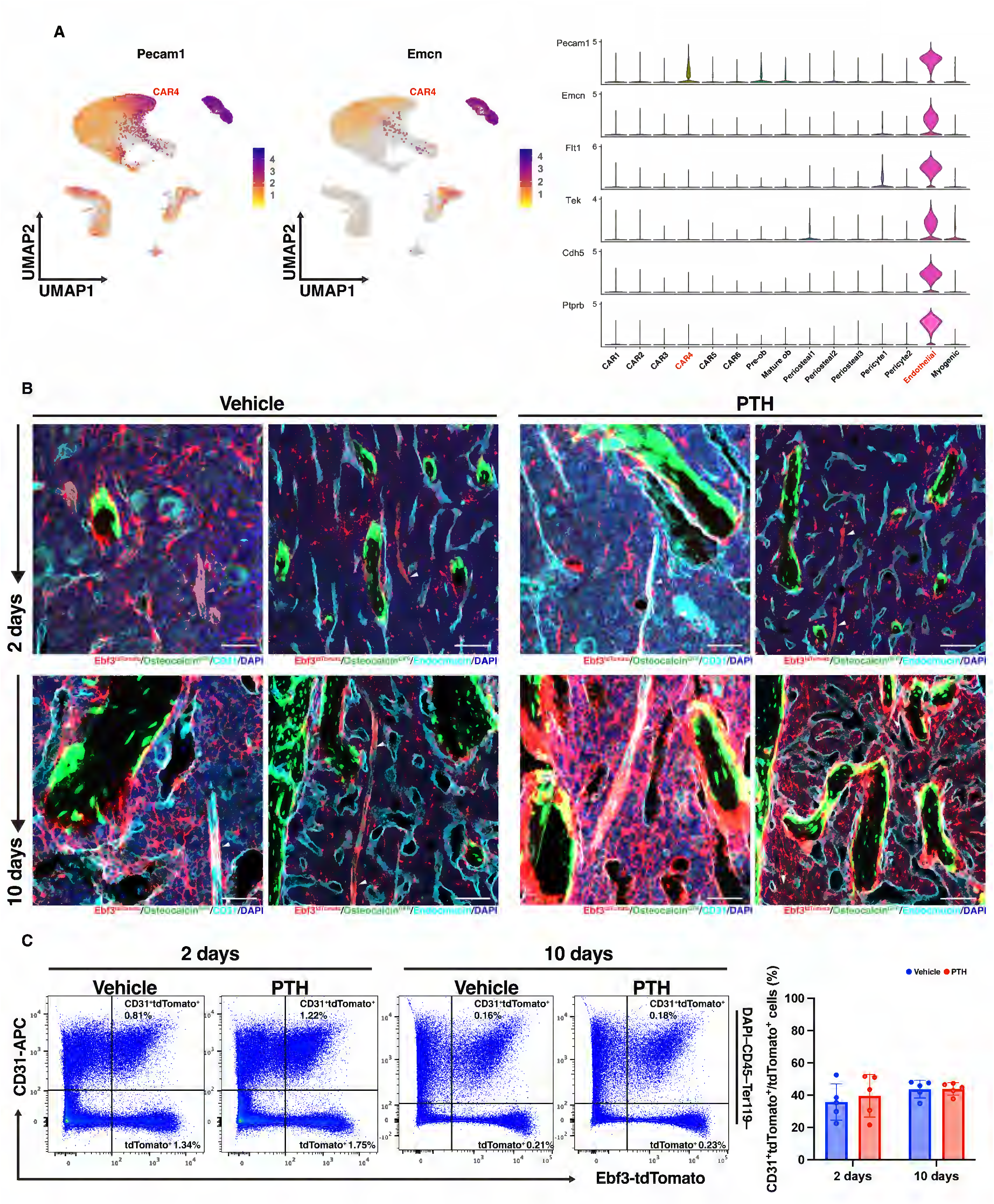
(**A**) Feature and violin plots of endothelial marker genes Pecam1 (CD31) and Emcn (Endomucin) in Ebf3-lineage cell clusters. (**B**) Immunohistochemical staining for CD31 and Endomucin in proximal tibia tissue sections from Ebf3-CreERT2; Rosa26-LSL-tdTomato; Bglap-GFP mice labeled at 10 weeks of age, following 2 and 10 days of iPTH or vehicle treatment. Cyan = CD31 or Endocmucin; red = Ebf3-CreERT2-tdTomato; green = Bglap-GFP. Scale bar = 100 µm. (**C**) Flow cytometry analysis of CD31 from Ebf3-CreERT2-tdTomato+ cells following 2 and 10 days of iPTH or vehicle treatment.

**Supplementary Figure 10.**
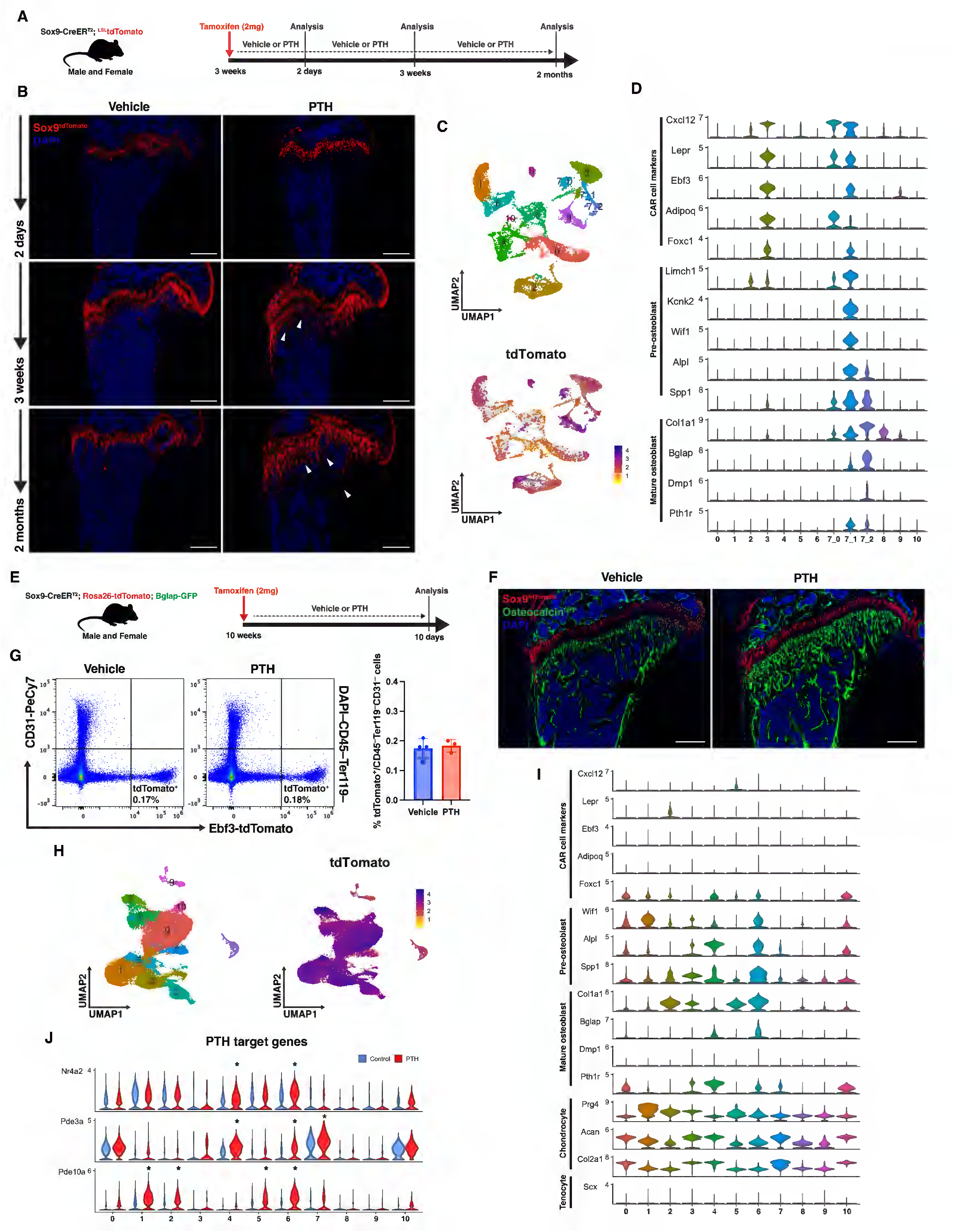
(**A**) Overview of lineage tracing strategy to study Sox9-lineage cells using Sox9-CreERT2; Rosa26-LSL-tdTomato mice. Lineage tracing Sox9-lineage cells in young mice: tamoxifen was given at 3 weeks of age and sacrificed after 2 days, 3 weeks, and 2 months of iPTH or vehicle treatment. (**B**) Representative fluorescent images of the proximal humerus showing Sox9-CreERT2-tdTomato+ cells labeled at 3 weeks of age, following 2 days, 3 weeks, and 2 months of iPTH or vehicle treatment. Scale bar = 500 µm. (**C**) UMAP and feature plot of tdTomato mRNA expression of Sox9-lineage cell clusters following Harmony-integration of libraries from 2 days, 3 weeks, and 2 months of iPTH or vehicle treatment. (**D)** Violin plots showing mRNA expression levels of CAR cell, pre-osteoblast, and mature osteoblast marker genes in Sox9-lineage cell clusters when tamoxifen is given to 3-week-old Sox9-CreERT2; Rosa26-LSL-tdTomato mice. (**E**) Overview of lineage tracing strategy to study Sox9-lineage cells using Sox9-CreERT2; Rosa26-LSL-tdTomato mice. Lineage tracing Sox9-lineage cells in adult mice: tamoxifen was given at 10 weeks of age and sacrificed after 10 days of iPTH or vehicle treatment. (**F**) Fluorescent images of the proximal tibia showing Sox9-CreERT2-tdTomato+ cells following 10 days of iPTH or vehicle treatment. Scale bar = 500 µm. (**G**) Flow cytometry analysis of Sox9-CreERT2-tdTomato+ cell numbers after 10 days of iPTH or vehicle treatment. (**H**) UMAP and feature plot of tdTomato mRNA expression of Sox9-lineage cell clusters following Harmony-integration of libraries from 10 days of vehicle and iPTH treatment. (**I**) Violin plots showing mRNA expression levels of CAR cell, pre-osteoblast, mature osteoblast, and tenocyte marker genes in Sox9-lineage cell clusters when tamoxifen is given to 10-week-old Sox9-CreERT2; Rosa26-LSL-tdTomato mice. (**J**) Violin plots showing the expression levels of canonical PTH target genes of each Sox9-lineage cell clusters with PTH versus vehicle treatment.

**Supplementary Figure 11.**
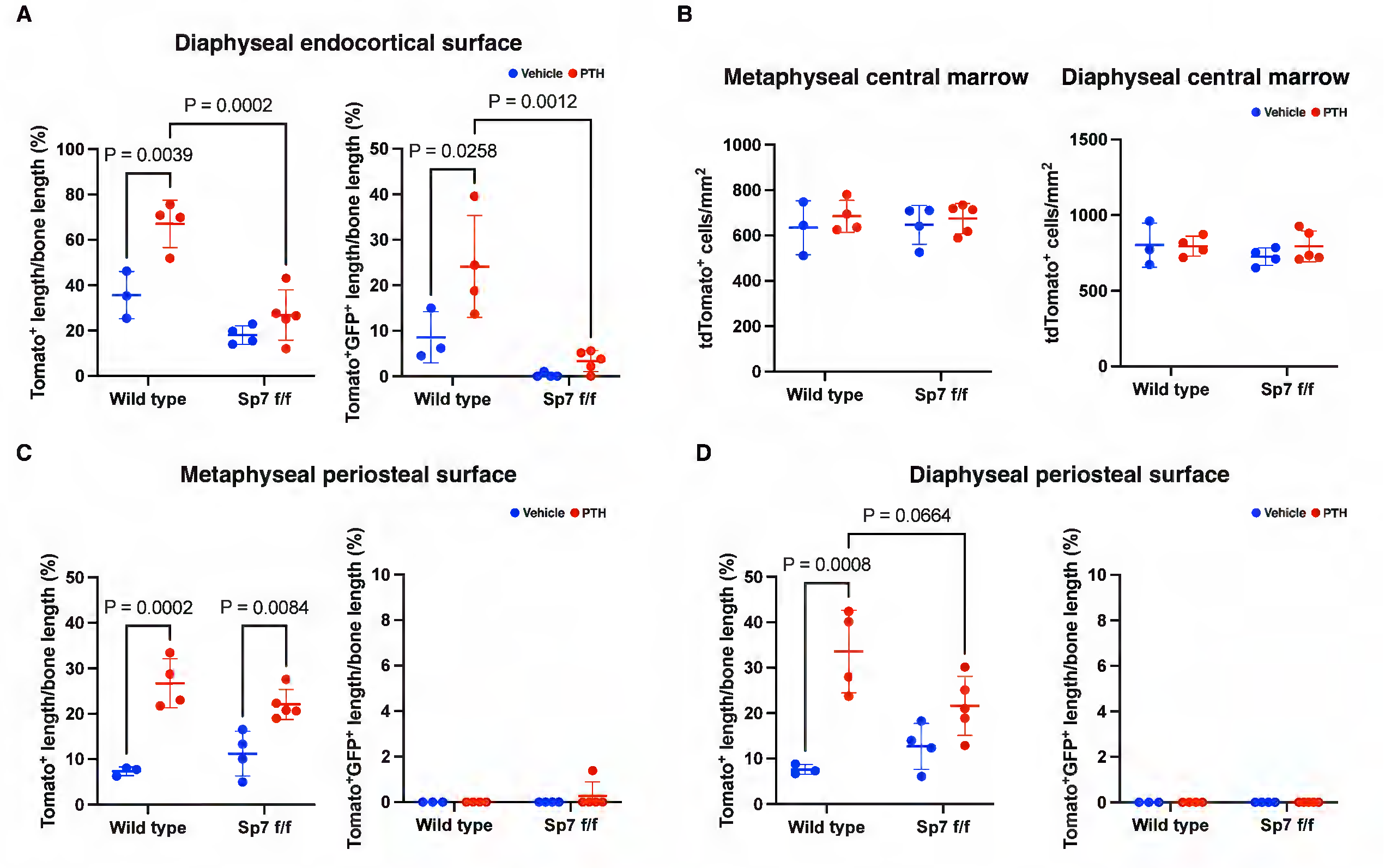
Fluorescent image quantification of Ebf3-labeled tdTomato+ and tdTomato+Bglap-GFP+ cells from wild type and Sp7 f/f mice labeled at 10 weeks old, in the (**A**) diaphyseal endocortical, (**B)** metaphyseal and diaphyseal central marrow, (**C**) metaphyseal and (**D**) diaphyseal periosteal skeletal regions following 10 days of iPTH or vehicle treatment. Statistical test: Two-way ANOVA with treatment and genotype (**A-D**) as factors followed by Šidák’s multiple comparisons test. Data is expressed as mean ± SD, n = 3-5 mice/group.

**Supplementary Figure 12.**
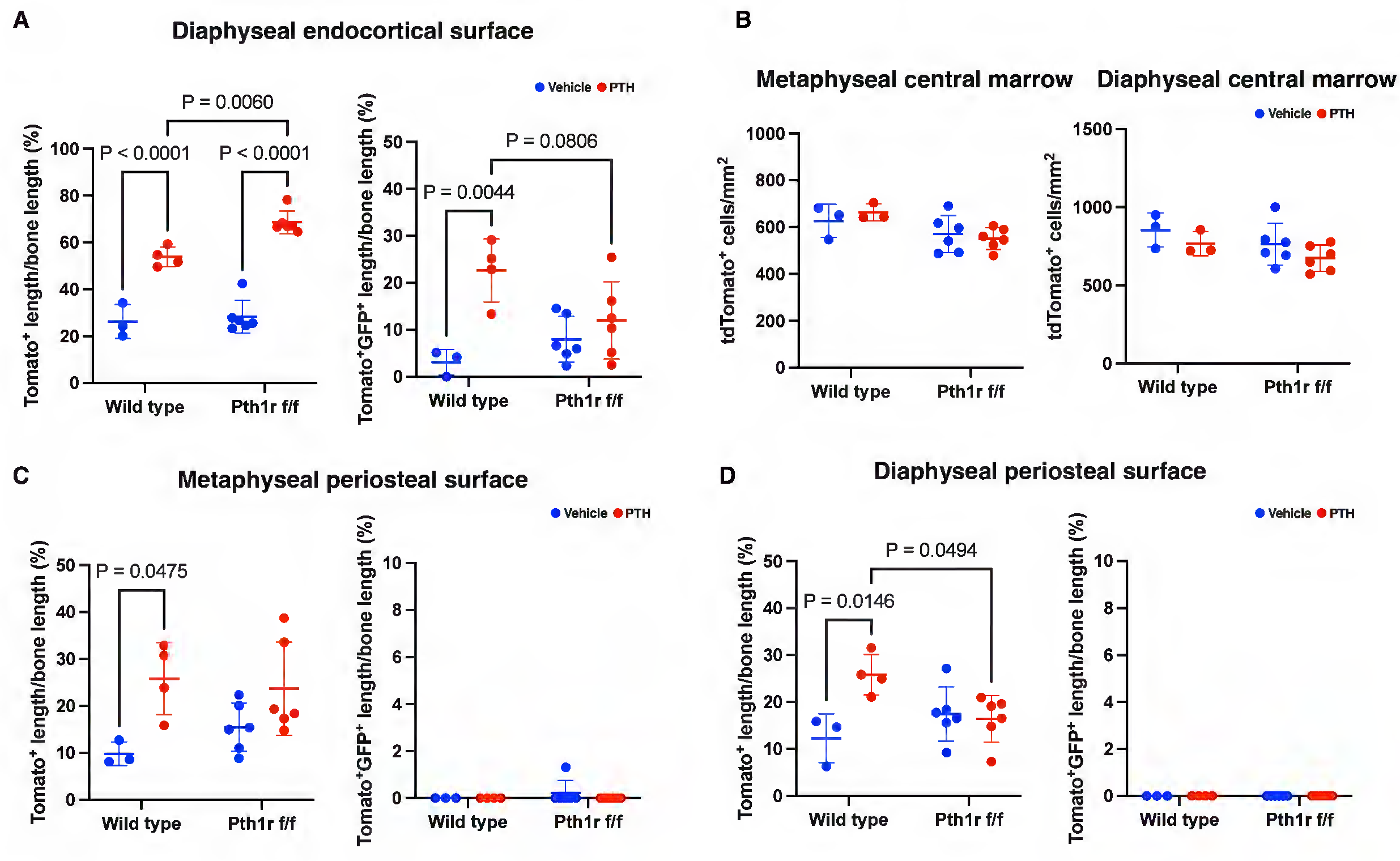
Fluorescent image quantification of Ebf3-labeled tdTomato+ and tdTomato+Bglap-GFP+ cells from wild type and Pth1r f/f mice labeled at 10 weeks old, in the (**A**) diaphyseal endocortical, (**B)** metaphyseal and diaphyseal central marrow, (**C**) metaphyseal and (**D**) diaphyseal periosteal skeletal regions following 10 days of iPTH treatment or vehicle. Statistical test: Two-way ANOVA with treatment and genotype (**A-D**) as factors followed by Šidák’s multiple comparisons test. Data is expressed as mean ± SD, n = 3-6 mice/group.

**Supplementary Figure 13.**
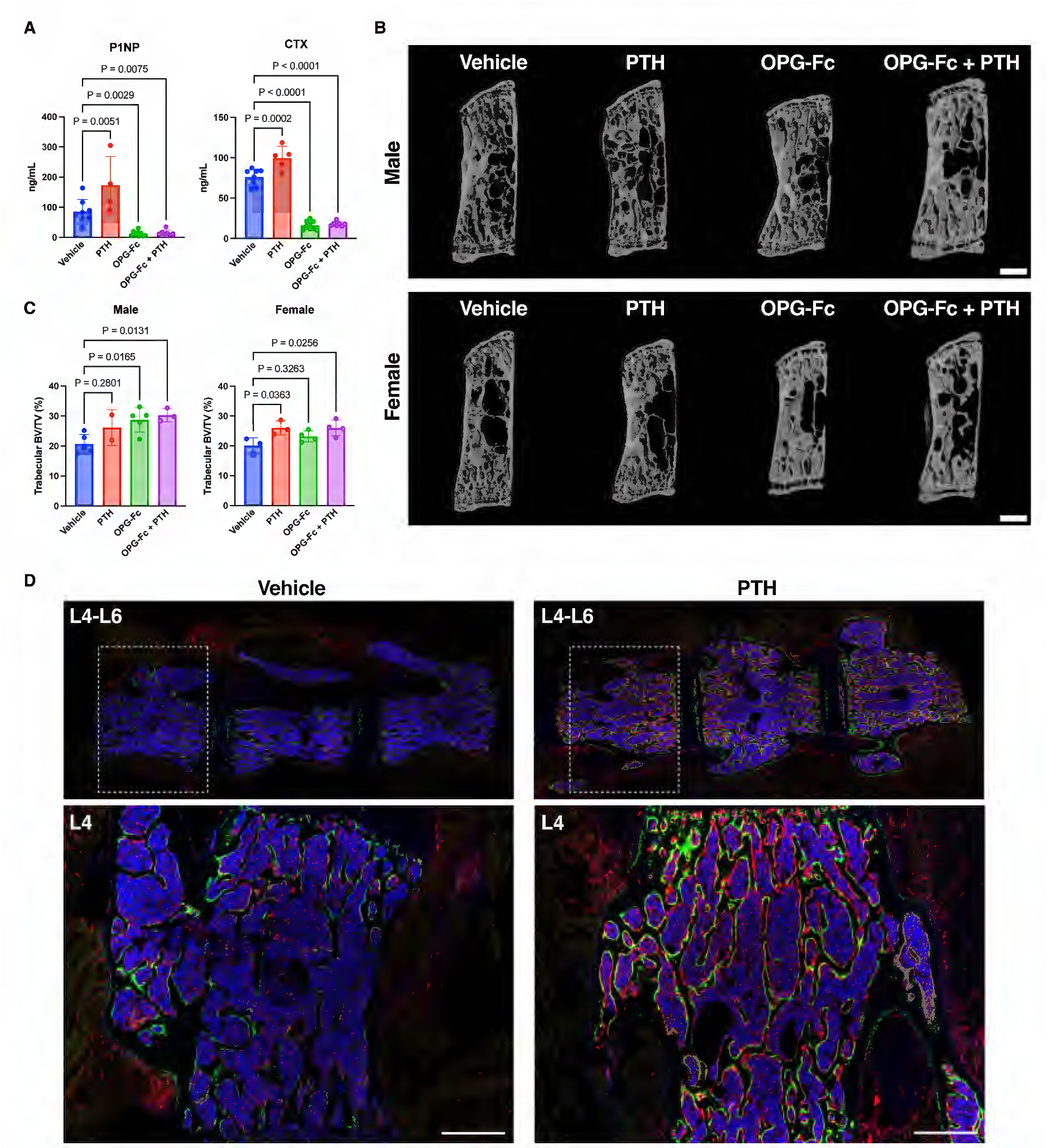
(**A**) Levels of serum bone formation marker, P1NP, and resorption marker, CTX, from 10-week-old Ebf3-CreERT2; Rosa26-LSL-tdTomato mice following 10 days of iPTH, OPG-Fc, combination, or vehicle treatment. (**B**) Representative MicroCT images and (**C**) analysis of the L5 vertebral body after 10 days of iPTH, OPG-Fc, combination, or vehicle treatment. Scale bar = 500 µm. (**D**) Representative fluorescent images of the lumbar spine from Ebf3-CreERT2; Rosa26-LSL-tdTomato mice labeled 10 weeks old, following 10 days of iPTH or vehicle treatment. Scale bar = 500 µm. Statistical test: One-way ANOVA followed by Šidák’s multiple comparisons test (**A and C**). Data is expressed as mean ± SD, n = 5-8 mice/group.

**Supplementary Figure 14.**
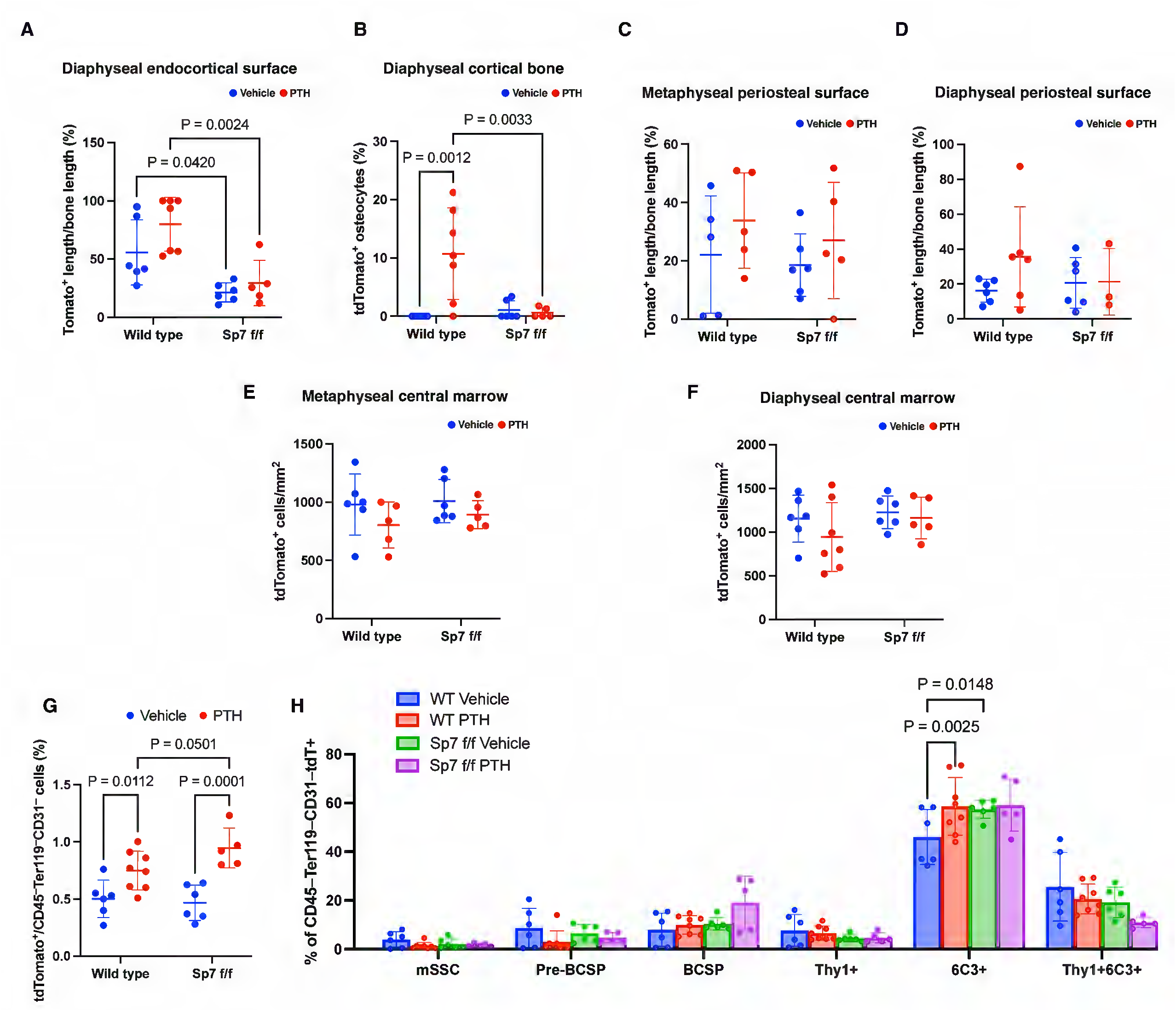
(**A**) Fluorescent image quantification of Ebf3-CreERT2-labeled tdTomato+ cells from wild type and Sp7 f/f mice in the (**A**) diaphyseal endocortical bone surface, (**B**) diaphyseal cortical bone, (**C)** metaphyseal and (**D**) diaphyseal periosteal bone surface, and (**E**) metaphyseal and (**F**) diaphyseal central marrow skeletal regions following 1 month of iPTH or vehicle treatment. (**G**) Flow cytometry analysis of non-hematopoietic Ebf3-CreERT2-tdTomato+ cells (**H**) and their skeletal stem/progenitor cell surface markers (**J**) from wild type and Sp7 f/f mice following 1 month of iPTH or vehicle treatment. Statistical test: Two-way ANOVA with treatment and genotype (**A-G**), or treatment and subpopulations (**H**) as factors followed by Šidák’s multiple comparisons test. Data is expressed as mean ± SD, n = 5-8 mice/group.

**Supplementary Figure 15.**
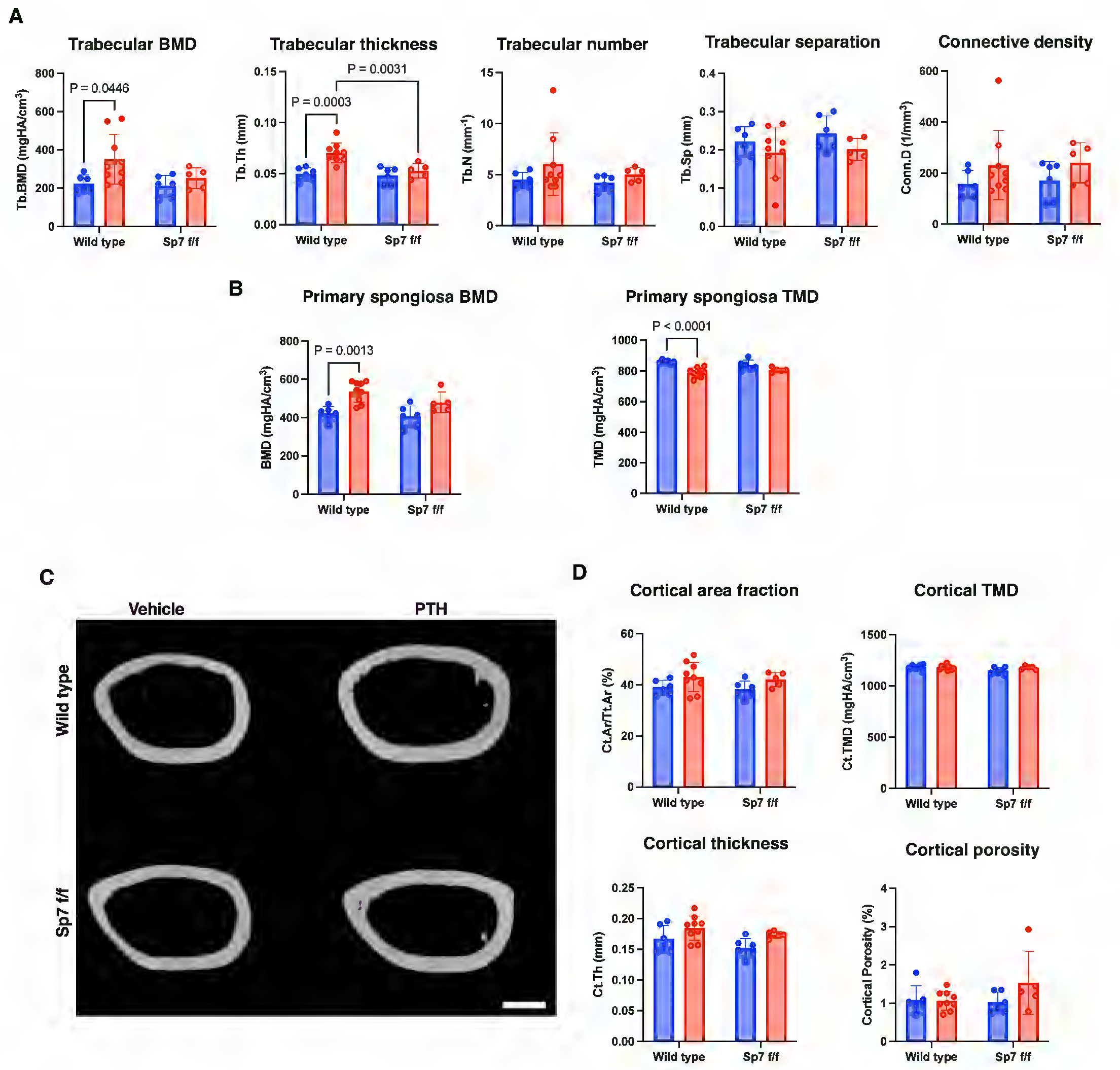
MicroCT analysis of the distal femur metaphysis showing **(A-E)** trabecular, (**F and G**) primary spongiosa bone parameters. (**H**) Representative microCT images of the femur mid-dialysis and analysis of the (**I-M**) cortical bone parameters. Scale bar = 500 µm. Statistical test: Two-way ANOVA with treatment and genotype (**A, B and D**) as factors followed by Šidák’s multiple comparisons test. Data is expressed as mean ± SD, n = 5-8 mice/group.

**Supplementary Figure 16.**
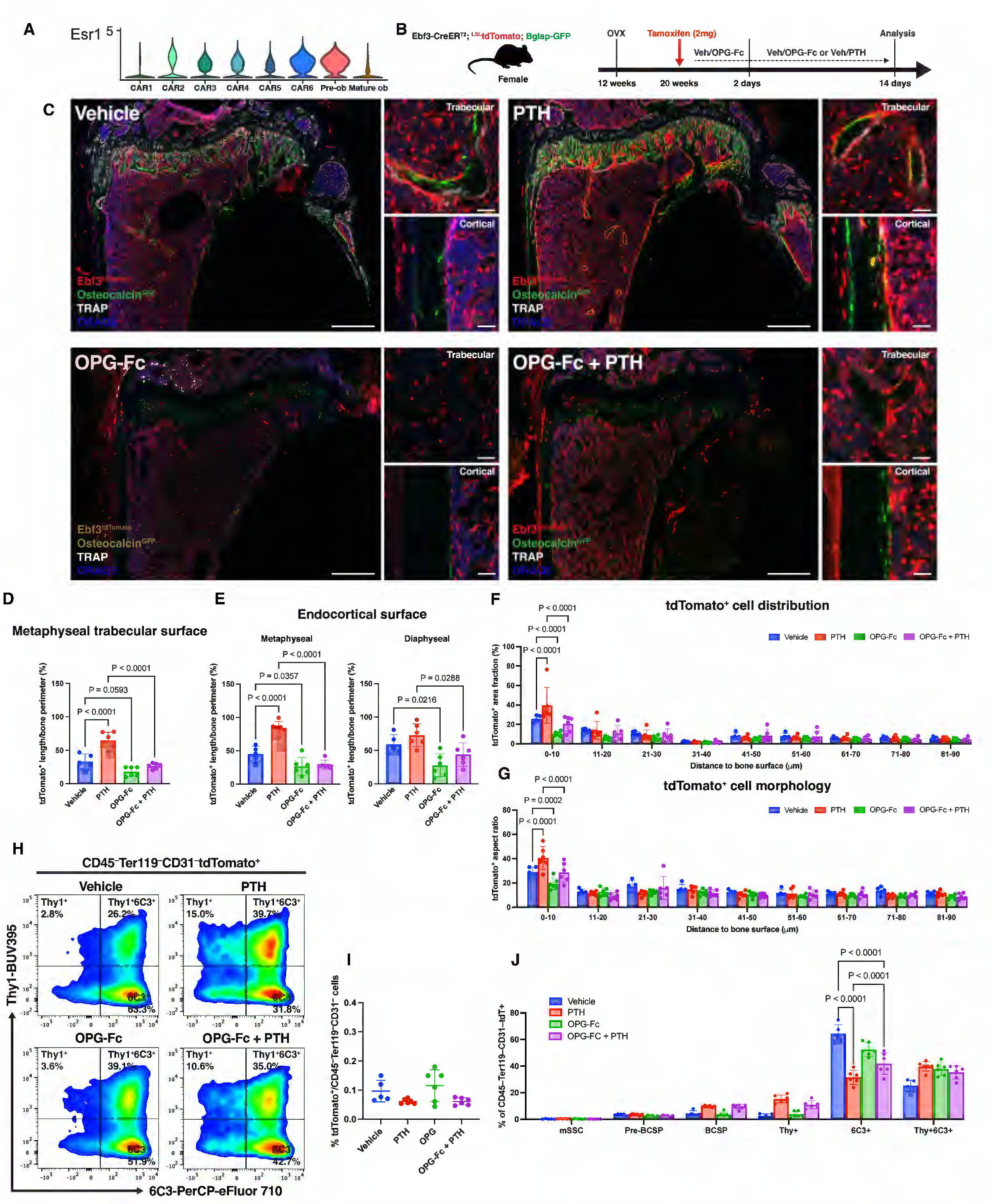
(**A**) Violin plots showing Esr1 expression level across Ebf3-lineage CAR and osteoblast clusters. (**B**) Overview of lineage tracing strategy to study the effects of iPTH, OPG-Fc, or combination treatment on Ebf3-lineage cells under estrogen-deficient setting using female ovariectomized Ebf3-CreERT2; Rosa26-LSL-tdTomato mice. (**C**) Representative fluorescent images of the proximal tibia from ovariectomized Ebf3-CreERT2; Rosa26-LSL-tdTomato; Bglap-GFP mice following 2 weeks of iPTH, OPG-Fc, combination, or vehicle treatment. White = TRAP; Red = Ebf3-CreERT2-tdTomato; green = Bglap-GFP; blue = DAPI. Scale bar = 500 µm (lower magnification) and 50 µm (higher magnification). (**D**) Quantification of tdTomato+ cells on metaphyseal trabecular and (**E**) endocortical bone surfaces following 2 weeks of iPTH, OPG-Fc, combination or vehicle treatment. (**F**) Distribution (area fraction) and (**G**) cell morphology (aspect ratio) analysis of tdTomato+ cells relative to defined distances from bone surfaces in the proximal tibia. (**H**) Representative flow cytometry plots and analysis of (**I**) non-hematopoietic Ebf3-CreERT2-tdTomato+ cells and their (**J**) skeletal stem/progenitor cell surface markers from ovariectomized Ebf3-CreERT2; Rosa26-LSL-tdTomato; Bglap-GFP mice following 2 weeks of iPTH, OPG-Fc, combination, or vehicle treatment. Statistical test: One-way ANOVA followed by by Šidák’s multiple comparisons test (**D, E, and I**), and two-way ANOVA with treatment and specified distance bins (**F and G**), or treatment and subpopulations (**J**) as factors followed by Šidák’s multiple comparisons test. Data is expressed as mean ± SD, n = 5-6 mice/group.

**Supplementary Figure 17.**
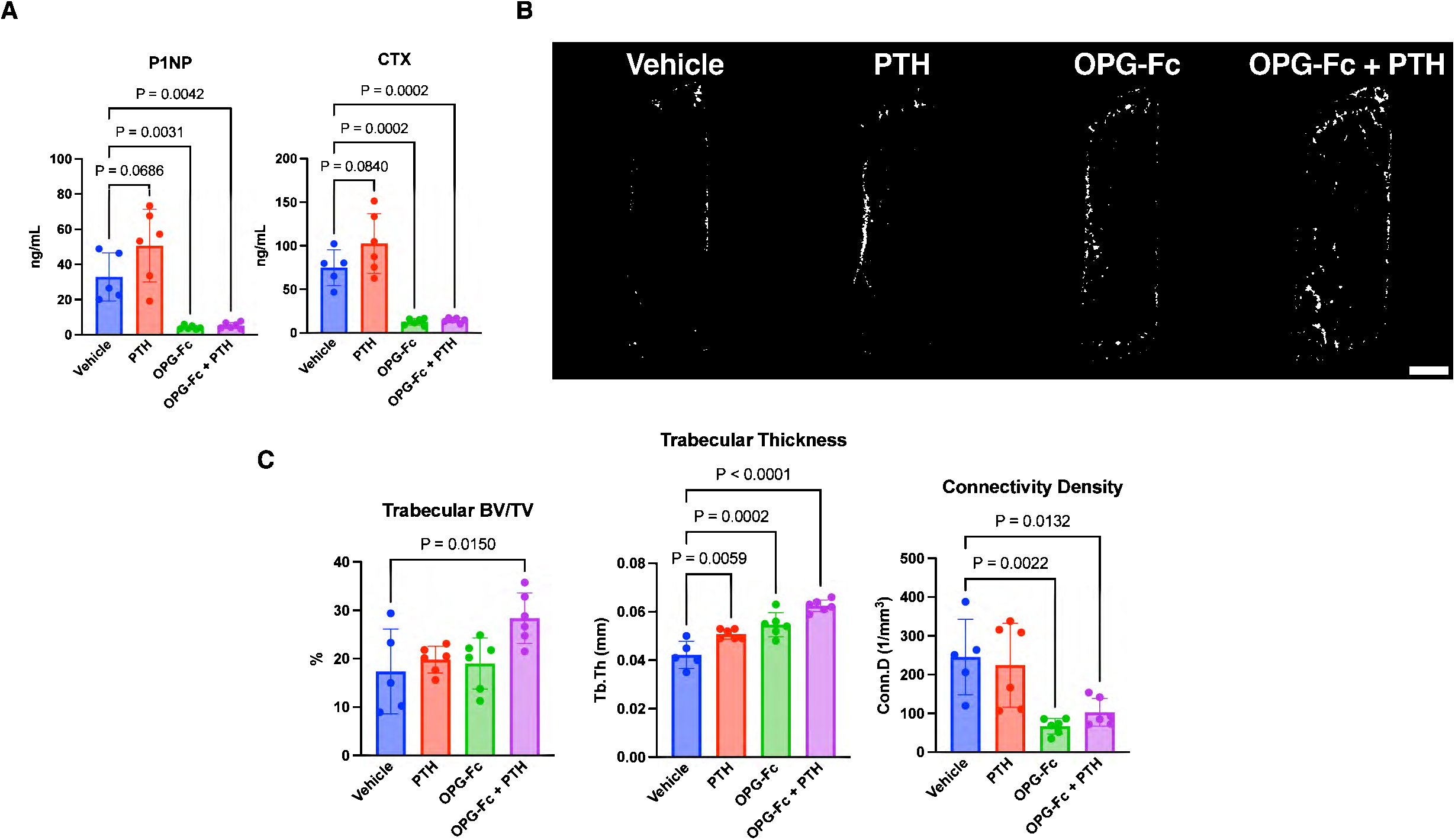
(**A**) Levels of serum bone formation marker, P1NP, and resorption marker, CTX, from 20-weeks-old ovariectomized Ebf3-CreERT2; Rosa26-LSL-tdTomato mice following 2 weeks of iPTH, OPG-Fc, combination, or vehicle treatment. (**B**) Representative MicroCT images and (**C**) analysis of the L5 vertebral body after 10 days of iPTH, OPG-Fc, combination, or vehicle treatment. Statistical test: One-way ANOVA followed by Šidák’s multiple comparisons test (**A, C and D**). Data is expressed as mean ± SD, n = 5-6 mice/group.

## Methods

### Mouse strains

Mice in this study were generated from Ebf3-CreERT2^11^, Sox9-CreERT2^75^, Bglap-GFP^76^, Cxcl12-GFP^77^, Pth1r-floxed^78^, Sp7-floxed^79^, Rosa26-loxP-stop-loxP-tdTomato (Ai14; JAX007914). All mice were backcrossed to C57BL/6J background for at least 10 generations. Mice were genotyped by PCR. Both male and female mice were used in all experiments unless otherwise indicated in the figure legends. Mice were housed in a pathogen-free, temperature- and humidity-controlled facility with husbandry maintained and provided by the Center of Comparative Medicine at the Massachusetts General Hospital. All animal experiments were approved and carried out in accordance with the guidelines issued by the Institutional Animal Care and Use Committees at the Center for Comparative Medicine at Massachusetts General Hospital under approved animal use protocol (2004N000176). At sacrifice, animals were euthanized by CO_2_ inhalation, followed by cervical dislocation.

### Tamoxifen administration

Tamoxifen (Sigma-Aldrich, T5648) was first dissolved in 100% ethanol, then an equal volume of sunflower seed oil (Sigma-Aldrich, S5007) was added and vigorously mixed. The tamoxifen-ethanol-oil mixture was incubated at 65°C until ethanol was completely evaporated and then stored at 37°C until use. Mice were given one dose of tamoxifen (2 mg) via intraperitoneal injection to induce Cre-dependent recombination in labeled cells for lineage tracing experiments, except for experiments in Figure 8, mice were given 3 doses of tamoxifen (2 mg) every other day to increase labelling and Sp7 deletion efficiency. For no tamoxifen controls, the same volume of sunflower oil (without tamoxifen) was injected.

### Parathyroid hormone and OPG-Fc administration

Mice were given daily subcutaneous injections of vehicle (10 mM citric acid, 150 nM NaCl, 0.05% Tween-80) or human PTH 1-34 (100 µg/kg). For OPG-Fc treatment^80^, mice were given once per week subcutaneous injections of vehicle (phosphate buffered saline; PBS) or OPG-Fc (5 mg/kg).

### Histology

Harvested bone samples were immediately fixed with 4% paraformaldehyde for 24 hours at 4°C, then decalcified in 15% EDTA solution (changed every other day) for 14 days. Decalcified samples were cryoprotected in 30% sucrose/PBS solution for 24 hours at 4°C, followed by a 30% sucrose/PBS:OCT (1:1) solution for 24 hours. Samples were embedded in OCT compound (TissueTek, Sakura), and frozen on a block of dry ice. Frozen sample blocks were cryosectioned (Leica CM1850) serially at 10 µm thickness and adhered to positive charged glass slides (Fisherbrand Superfrost plus).

Cell quantification at distinct skeletal compartments of the tibia was performed on fluorescent images. At each skeletal site, a 500 µm x 750 µm region of interest (ROI) was defined. At the proximal metaphysis, the ROI for central marrow and cortical bone analysis was positioned 500 µm below the growth plate. At the midshaft diaphysis, the ROI for central marrow and cortical bone analysis was positioned mid-diaphysis. Cortical bone parameters (endosteal surface/periosteal surface/osteocyte number) were assessed at the compression side of the tibia. Additional details of these regions are depicted in **Supplementary Figure 1C.** Two serial sagittal sections per mouse corresponding to the central sagittal plane of the tibia were quantified by two different scientists blinded to the condition. Mean values per mouse were used in statistical analysis.

### MATLAB image analysis

Fluorescence images of the metaphysis of the tibia were analyzed using a custom pipeline implemented in MATLAB to quantify the spatial distribution of Ebf3-lineage tdTomato+ cells relative to bone surfaces. Multi-channel images were imported and separated into nuclei (DAPI), tdTomato+ signal (distance source), and bone surfaces (distance targets). The nuclei channel was used to generate a binary mask defining the bone marrow space and excluding bone and tissue artifacts. Bone surfaces were identified from the edges of the marrow mask and classified into epiphyseal/growth plate, endocortical bone, and trabecular bone regions. TdTomato+ cells were segmented from the fluorescence channel following background correction and size/morphology filtering. Distances between tdTomato+ cells and endocortical bone surface were computed using Euclidean distance maps constrained to the marrow mask. All spatial measurements were converted to micrometers (1.8872 µm per pixel) based on microscope calibration. Each segmented tdTomato+ cell was assigned to the nearest endocortical bone surface, and assignment quality was evaluated using a clustering-based score. tdTomato+ cell distributions (area fraction) and morphological features (aspect ratio) relative to specified distance bins from the endocortical bone surface were quantified and exported for downstream statistical analysis.

### Microscopy

Multicolor fluorescent images were captured using the Keyence BZ-X700 fluorescent microscope, with preconfigured filter settings for DAPI/FITC/Texas Red/Cy5. Custom filter for ELF 97 (excitation/emission = 350 nm/540 nm) was used to detect fluorescent TRAP staining. Z-stack confocal images were acquired using Zeiss LSM 800 Airyscan confocal microscope with 4 lasers and corresponding filters for DAPI/GFP/ tdTomato/Cy5. Representative images of at least three independent biological samples are shown in the figures. Quantification of fluorescent images was performed using Image J software (National Institute of Health).

### Immunohistochemistry

Immunohistochemical staining of mouse tibia, tissue section underwent antigen retrieval using proteinase K solution (Dako, S3020) for 5 minutes at room temperature, permeabilized with 0.2% TritonX/PBS for 30 minutes at room temperature, blocked with 3% bovine serum albumin (BSA) in 0.2% TritonX/PBS solution for 1 hour at room temperature, followed by overnight incubation at 4°C with rat anti-endomucin (Emcn) monoclonal antibody (1:100, Santa Cruz Biotechnology, sc65495), rat anti-CD31 monoclonal antibody (1:100, Bio-Rad, MCA2388), rabbit anti-perilipin A/B polyclonal antibody (1:100, Sigma-Aldrich, P1873), or rabbit anti-phospho-SMAD2/SMAD3 polyclonal antibody (1:100, ThermoFisher, PA5-110155). Negative controls consisted of isotype-matched primary antibodies at the same protein concentration. The next day, sections were incubated with Alexa Fluor 647-conjugated goat anti-rat (Invitrogen, A-21245) or anti-rabbit (Invitrogen, A-21247) secondary antibodies at room temperature for 3 hours. Nuclei were stained with DAPI (4′,6-diamidino-2-phenylindole, 1 µg/ml, Invitrogen, D1306) for 5 minutes at room temperature, then cover slipped and mounted using Fluoromount (SouthernBiotech, OB100-01).

### Bone single cell isolation

A single-cell suspension of bone cells was prepared using a slightly modified version of previously described protocol^28^. Briefly, femurs and tibias from a single mouse were harvested and stripped of muscle tissue. Whole bones from each mouse were placed in a mortar containing 1 ml of Hank’s Balanced Salt Solution (HBSS, Sigma-Aldrich, H6648) and mechanically homogenized using a pestle. Thereafter, this bone-cell solution was transferred into a single well of a 12-well culture dish containing HBSS and 2 Wünsch units of Liberase^TM^ (Roche, 5401127001) and placed on a shaking incubator (ThermoMixer C, Eppendorf) at 37°C for 20 minutes. After each round of enzymatic digestion, cells were mechanically triturated using a P1000 pipette with tip end cut off, then cell supernatant was collected and filtered through a 70 µm cell strainer (Falcon, 352350) into a 50 mL tube on ice. The bone fragments were digested a second time with Liberase^TM^ and the released cells were collected as described above. After this, the bone fragments were incubated in PBS containing 5 mM EDTA and 0.1% BSA for 20 minutes at 37°C for 20 minutes, and then collected the released cells as described above. Remaining bone fragments were subjected to alternating exposures of either Liberase^TM^ or 5 mM EDTA solution to yield a total of 4 Liberase^TM^ digestions and 3 EDTA incubations. After 7 rounds of digestion, the collected cells were pelleted and resuspended in sorting buffer (2% fetal bovine serum in HBSS; FACS buffer).

### Fluorescence activated cell sorting (FACS)

For FACS isolation of mouse tdTomato+ stromal cells for single cell RNA-sequencing experiments, mouse bone single cell suspension collected after 7 rounds of serial enzymatic digestion underwent magnetic activated cell depletion of hematopoietic cells using mouse lineage depletion kit (Miltenyi Biotec, 130-110-470) combined with CD45 microbeads (Miltenyi Biotec, 130-052-301) according to the manufacturer’s instructions. After magnetic depletion, cells were incubated in the dark for 30 minutes on ice with the prepared primary antibody cocktail. Cells were washed 2 times and resuspended in FACS buffer with DAPI (1 µg/ml, Invitrogen, D1306). FACS was performed using BD FACSAria^TM^ II equipped with 5 lasers (BD bioscience). Stromal tdTomato+ fraction identified as CD45–Ter119–tdTomato+ was collected for 10X single cell RNA-sequencing.

For FACS isolation of human bone marrow stromal cells for single cell RNA-sequencing experiments, human bone marrow cells were incubated with blocking buffer (0.5% BSA in PBS; 1:50, BD bioscience, 564220) for 10 minutes on ice, then incubated in the dark for 30 minutes on ice with primary antibody cocktail with cell viability dye Calcein AM (5 µM; ThemoFisher, C3099). Cells were washed 2 times and resuspended in sorting buffer (0.5% BSA in PBS). FACS was performed using Bigfoot Spectral Cell Sorter equipped with 7 lasers (Invitrogen). Stromal fraction identified as CD45–CD235a–CD14– was collected for 10X single cell RNA-sequencing.

FACS analysis for mouse skeletal stem and progenitor subpopulations was performed on mouse bone single cell suspension collected after 7 rounds of serial enzymatic digestion. Primary antibody dilutions were prepared in Brilliant Stain buffer (BD bioscience, 563794), and cells were incubated in the dark for 30 minutes on ice with the prepared primary antibody cocktail. Cells were washed 2 times and resuspended in FACS buffer with DAPI (1 µg/ml, Invitrogen, D1306). FACS was performed using Bigfoot Spectral Cell Sorter equipped with 7 lasers (Invitrogen). Mouse bone marrow cells and UltraComp eBeads (Invitrogen, 01-333-42) were used to set initial fluorochrome compensation. Fluorescence minus one (FMO) controls were used for additional compensation and to assess background levels for each stain. Gates were drawn as determined by internal FMO controls to define positive and negative populations for each cell surface marker. Per biological sample, 5-8 million events were recorded and analyzed using FlowJo (10.1v). Full details of the FACS gating strategies shown in **Supplementary Figure 4A.**

### FACS antibodies

Antibodies for FACS of mouse samples included CD45 (eBioscience, 17-5921-82; BD bioscience, 568344), Ter119 (eBioscience, 17-0451-82; BD bioscience, 741736), CD31 (BD bioscience, 551262), CD200 (BD bioscience, 565547), CD105 (BioLegend, 120410), Thy1 (BD bioscience, 565257), 6C3 (eBioscience, 46-5891-82), CD51 (BD bioscience, 740946). Antibodies for FACS of human samples included CD235ab (BioLegend, 306614), CD14(BioLegend, 301824), CD45 (BioLegend, 304052), CD31 (BioLegend, 303122).

### Cell proliferation and apoptosis flow cytometric assay

To evaluate cell proliferation, 1.5 mg of 5-ethynyl-2’-deoxyuridine (EdU) (Invitrogen, A10044) was dissolved in PBS and administered via intraperitoneal injection to mice 12 hours before sacrifice. Bone cells were isolated as described above in the bone single cell isolation section. Proliferative cells (EdU+) were detected using the Click-iT^TM^ Alexa Fluor 488 flow cytometry assay kit (Invitrogen, C10425). Apoptotic cells were detected using the FITC Annexin V Apoptosis detection kit (BioLegend, 640922). Both kits were used following the manufacturer’s instructions.

### In vitro culture of Ebf3-lineage cells

For intracellular cAMP quantification, FACS-isolated mouse tdTomato+ stromal cells were seeded in tissue culture 96-well plate at a density of 1 x 10^4^ cells per well, with 3 technical replicates per mouse sample. For phospho-CREB immunocytochemistry, FACS-isolated stromal cells were seeded in an 8-well cell culture chamber slide at a density of 5 x 10^2^ cells per well. For phospho-CREB flow cytometry analysis, FACS-isolated stromal cells per mouse sample were seeded in a 24-well plate at a density of 5 x 10^4^ cells per well. Cells were cultured at 37°C and 5% CO_2_ in low-glucose DMEM with GlutaMAX supplement (Gibco, 10567022) and 10% mesenchymal stem cell-qualified FBS (Gibco, 12662029) for 7 days before PTH stimulation experimentation.

### Intracellular cAMP assay

Prior to stimulation, cells were washed HB solution (1M HEPES, 7.5% BSA in HBSS). Cells were then treated for 30 minutes at room temperature with either vehicle, human PTH 1-34 (100 nM), or forskolin (30 µM) in a IBMX buffer solution (0.2M IBMX and 5N NaOH in HB solution). After, IBMX buffer solution was aspirated and 50 mM HCL solution was added to the cells. Cell solution was used for intracellular cAMP quantification by cAMP Competitive ELISA Assay Kit (Abcam, ab290713) following manufacturer’s protocol. The signal was read on the EnVision plate reader (Perkin Elmer, 2104) at 405 nm.

### Phospho-CREB assays

For phospho-CREB immunocytochemical staining, post-stimulated cells were fixed with 4% paraformaldehyde solution at room temperature for 10 minutes and then washed 3 times with PBS. Cells were permeabilized with 0.2% Triton-X/PBS for 15 minutes, then blocked with 3% BSA in 0.2% TritonX/PBS at room temperature for 1 hour, followed by overnight incubation at 4°C with rabbit anti-phospho-CREB (1:200, Cell Signaling Technology, 9198S). Negative controls consisted of isotype-matched primary antibodies at the same protein concentration. The next day, cells were incubated with Alexa Fluor 488-conjugated goat anti-rabbit (Invitrogen, A-11008) at room temperature for 3 hours. Nuclei were stained with DAPI (4′,6-diamidino-2-phenylindole, 1 µg/ml, Invitrogen, D1306) for 5 minutes at room temperature, then cover slipped and mounted using Fluoromount (SouthernBiotech, OB100-01).

For phospho-CREB flow cytometric analysis, post-stimulated cells were trypsinized and immediately fixed with 4% paraformaldehyde solution at room temperature for 10 minutes. Cells washed with PBS and pelleted, and then permeabilized with ice-cold 90% methanol/PBS on ice for 10 minutes. After, cells were washed with PBS and incubated with anti-phospho-CREB-Alexa 488 conjugated antibody (1:50, Cell Signaling Technology, 9187S) at room temperature for 1 hour. Negative controls consisted of isotype-matched primary antibodies at the same protein concentration. Cells were washed 3 times with PBS and resuspended in PBS for flow cytometry.

### Single cell RNA-sequencing analysis

Cell viability and numbers were assessed using a Luna FX7 fluorescent cell counter (Logos Biosystem) before loading onto the Chromium GEM-X Single Cell 3’ Chip (10X Genomics). Single cell RNA-sequencing cDNA libraries were constructed using the Chromium GEM-X Single Cell 3’ Reagent Kit according to the manufacturer’s instructions, followed by sequencing of the cDNA libraries using the Illumina NextSeq 2000 instrument. Sequencing depth of all libraries ranged from 400-800 million reads to reach a target of 25,000 mean reads per cell. Raw reads obtained from scRNA-seq experiments were demultiplexed, aligned to the custom mouse mm10 genome (with tdTomato gene inserted) using Cellranger toolkit (10X Genomics). Analyses were performed using Seurat^81^ in R and Scanpy^82^ (scVelo^83^ and CellRank^42^) in Python. Cells were filtered out based the following criteria: less than 200 genes and more than 4,000 genes per cell and with more than 20% mitochondrial read content. After quality control filtering of low quality cells from each library, all libraries were then integrated using Harmony^27^ to correct for batch effects, followed by standard single cell processing steps including normalization, identification of highly variable genes across single cells, scaling based on number of Unique Molecular Identifier (UMI), dimensionality reduction (PCA and UMAP), unsupervised clustering, and identifying cluster-specific differentially expressed genes using the nonparametric Wilcoxon rank sum test included in Seurat.

Afterwards, contaminating hematopoietic cell clusters with low or no tdTomato mRNA gene expression were filtered out and the remaining clusters were re-clustered by performing the following single cell processing steps: identifying highly variable genes, scaling, dimensionality reduction, and unsupervised clustering at 0.2 resolution parameter, resulting in the final UMAP shown in the figures. Clustree^84^ was used to assess cluster stability across 0.2 to 1 resolution parameter to guide the selection of a biologically meaningful clustering resolution. Within each cluster, cells from the vehicle-and PTH-treated libraries were randomly down-sampled to equal numbers prior to performing differential gene expression analysis using the MAST hurdle model^74^. The list of significant differentially expressed genes (adjusted P-value < 0.05 and |log2FC| > 0.5) for each cluster was then used for gene set enrichment analysis (GSEA) by Enrichr^85^ (https://maayanlab.cloud/Enrichr/) to identify significantly enriched gene sets (adjusted P-value < 0.05).

For the RNA velocity-based fate probability analysis, spliced and unspliced RNA matrices (loom files) for each library were generated using velocyto^86^ according to the standard protocol (https://velocyto.org/velocyto.py/tutorial/index.html). Next, RNA velocity matrices were then computed using the standard scVelo^83^ workflow with default parameters as described in its documentation (https://scvelo.readthedocs.io/en/stable/). Cell-cell transition matrices and cell fate probabilities were then inferred using the CellRank^42^ framework following the recommended pipeline (https://cellrank.readthedocs.io/en/latest/notebooks/tutorials/estimators/700_fate_probabilities.html).

To assess the transcriptomic effects of PTH on human CAR cells, single cell RNA-sequencing libraries from FACS-isolated human bone marrow stromal cells (described below) were integrated using Harmony with a single cell atlas composed of multiple recently published datasets derived from naïve human bone marrow^60^. Thereafter, the integrated dataset underwent quality control and single cell processing workflow (described above) for downstream analysis.

### RNA isolation and RT-qPCR

Total RNA was isolated from FACS-isolated tdTomato+ cells using PicoPure RNA isolation Kit (Applied Biosystems, KIT0204) according to manufacturer’s protocol. cDNA was prepared with 200 ng RNA and synthesized using the Primescript RT kit (Takara Inc). RT-qPCR was performed using PerfeCTa SYBR Green FastMix (Quanta bio) in the StepOnePlus^TM^ Real-Time PCR system (Applied Biosystems). Glyceraldehyde-3-phosphate dehydrogenase (*Gapdh)* was used as a control housekeeping gene. Relative expression was calculated using the 2^-deltaCT^ method. Primers used for RT-qPCR are listed in **Supplementary Table 1**.

### Tartrate-resistant acid phosphatase (TRAP) staining

Sections were incubated in TRAP buffer (0.92% sodium acetate anhydrous, 1.14% L-(+) tartaric acid, 1% glacial acetic acid, pH 4.1-4.3) at room temperature for 10 minutes, then incubate with ELF97 phosphatase substrate (125 µM, Invitrogen, E6588) in TRAP buffer under UV light at room temperature for 15 minutes. Sections were washed 2 times with PBS, followed by 2 more washes with TRAP washing buffer (20 mM EDTA, 5 mM levamisole hydrochloride, pH 9). Nuclei were stained with DRAQ5 (1:1000 in PBS, Thermo Scientific, 62251) at room temperature for 5 minutes, washed with PBS, then cover slipped and mounted using Fluoromount (SouthernBiotech, OB100-01).

### Ovariectomy

Twelve-week-old female C57/BL6J mice were subjected to either bilateral ovariectomy (OVX) or sham surgery. For analgesia, carprofen (10 mg/kg) was injected subcutaneously before the operation, followed by 1 dose of carprofen every day for 3 days after the operation. Mice were anesthetized and maintained with 3% inhalant isoflurane delivered with 100% O2 (1 liter/min) via face mask. Eye lubricant was used to protect the eyes during surgery. The lower back was shaved and prepared with a triple application of betadine disinfectant alternating with isopropyl alcohol. Mice were then placed on a warm circulating water pad to maintain the body temperature during the operation. A sterile drape was fitted over the entire animal and only surgical site was exposed. The surgery began with a 1 cm dorsal midline skin incision caudal to the posterior border of the ribs. Skin was separated from the muscle layer and a small incision was made on the muscle layer to enter the abdominal cavity. Using sterile forceps, the peri-ovarian fat is gently pulled out to lift and visualize the ovary. The ovary is resected, and the uterine horn returned into the abdomen. The same process is repeated to remove the other ovary. Muscle layer was closed with a continuous suture using 6-0 absorbable Vicryl (Ethicon), and the skin closed with interrupted sutures. Wound clips were applied to reinforce wound closure. Sham operations involved the same incision approach and visualization of the ovary, but without its removal. After surgery, mice were returned to their home cage and allowed unrestricted movement and access to food and water. Eight weeks post-surgery (20 weeks old), mice were euthanized by CO2 inhalation followed by cervical dislocation. Uterine tissues were weighed to confirm successful ovariectomy.

### MicroCT analysis

A high-resolution desktop micro-tomographic imaging system (µCT40, Scanco Medical AG, Brüttisellen, Switzerland) was used to assess the trabecular bone microarchitecture, volume, and mineral density in the primary and secondary spongiosa of the distal femur and the L5 vertebral body of each mouse. Scans were acquired using a 10 µm3 isotropic voxel size, 70 kVp peak x-ray tube intensity, 114 mA x-ray tube current, and 200 ms integration time, and were subjected to Gaussian filtration and segmentation. The primary spongiosa region of interest started at the peak of the distal growth plate and extended proximally for 500 μm (50 transverse slices). The secondary spongiosa region of interest started immediately superior to the primary spongiosa region and extended 1500 μm (150 transverse slices) proximally. In both regions, bone was segmented from soft-tissue using a mineral density threshold of 400 mgHA/cm3. All analyses were performed using the Scanco Evaluation software. Trabecular bone in the endocortical area of the secondary spongiosa region was analyzed for bone volume fraction (Tb.BV/TV, %), trabecular thickness (Tb.Th, mm), trabecular number (Tb.N, mm-1), trabecular separation (Tb.Sp, mm), connectivity density (Conn.D, mm-3), and trabecular bone mineral density (Tb.BMD, mgHA/cm3). The bone in the primary spongiosa was analyzed for bone volume, total volume, bone volume fraction, and bone and tissue mineral densities. Cortical bone architecture and mineral density were analyzed at the femoral mid-diaphysis. Cortical bone was segmented with the mineral density of 700 mgHA/cm3 and then analyzed using the Scanco mid-shaft evaluation script to measure total cross-sectional area (bone + medullary area) (Tt.Ar, mm2), cortical bone area (Ct.Ar, mm2), medullary area (Ma.Ar, mm2), bone area fraction (Ct.Ar/Tt.Ar, %), cortical tissue mineral density (Ct.TMD, mgHA/cm3), cortical thickness (Ct.Th, mm), cortical porosity (%), as well as maximum, minimum and polar moments of inertia (Imax, Imin, and J, mm4). For L5 trabecular analysis, bone was evaluated in a cylindrical region of interest. The cylindrical region was a 2 mm (200 transverse slices) long cylinder (diameter = 750 µm, 75 voxels) that was centralized on the vertebral body. Bone was segmented from soft tissue using a mineral density threshold of 400 mgHA/cm3 and the following architectural parameters were measured using the Scanco Trabecular Bone Morphometry evaluation script: trabecular bone volume fraction (Tb.BV/TV, %), trabecular bone mineral density (Tb. BMD, mgHA/cm^3^), trabecular thickness (Tb.Th, mm), trabecular number (Tb.N, mm^-1^), and trabecular separation (Tb.Sp, mm), and connectivity density (Conn.D, 1/mm^3^).

### Serum bone turnover marker ELISAs

Twelve hour fasting serum was collected from mice just prior to sacrifice by retro-orbital bleed. Serum measurements for P1NP (IDS Immunodiagnostic Systems, AC-33F1) and CTX (IDS Immunodiagnostic Systems, AC-06F1) were performed using commercial bioassay detection kits following the manufacturer’s instructions.

### Human bone marrow stromal cells

DATA-Biopsy study design and participant characteristics are described previously^59,69^. Briefly, bone marrow aspirates were obtained from postmenopausal women receiving teriparatide (20 µg/day) subcutaneously for 3 months. Marrow aspirates were transferred into EDTA-coated tubes (Becton Dickinson, 366643) and placed on a rocker to prevent clotting. Bone marrow stromal cells were extracted by Ficoll-Paque gradient separation and prepared for FACS as described above.

### Statistical analysis

Statistical analyses were performed by GraphPad Prism 10 (GraphPad Software Inc, USA). For comparison between two experimental groups, Student’s two-tailed t tests was used. For more than two groups, one-way or two-way ANOVA followed by Šidák’s multiple comparisons test were used. Data is expressed as mean ± SD. Statistical significance was assigned at P < 0.05 with P values indicated in the figures. The number of mice per experiment is indicated in the figure legend.

## Acknowledgements

We thank Drs. Roland Baron, Francesca Gori, Yi-Hsiang Hsu, Douglas Kiel, Matthew Greenblatt, Christa Maes, Sundeep Khosla, Natalie Sims, Dolores Shoback, and all members of the Wein laboratory and MGH Endocrine Unit for valuable feedback. We thank Dr. Marie Demay for providing guidance and methods for isolating human bone marrow cells. BSKC is supported by Canadian Institutes of Health Research Postdoctoral Fellowship (MFE-201024). MNW acknowledges support from NIAMS (P50AR080596) and NIDDK (R01DK116716) and a Chen Institute MGH Research Scholar award. GP acknowledges support from NIAMS Clinical Investigator Award (K08AR084618). Bone MicroCT work was supported by the NIAMS-funded Center for Musculoskeletal Research (P30AR075042). Flow cytometry work was supported with expert technical assistance by the HSCI-CRM Flow Cytometry facility at Massachusetts General Hospital. Confocal imaging was performed in the Microscopy Core of the Program in Membrane Biology, which is partially supported by a Centre for the Study of Inflammatory Bowel Disease Grant DK043351 and a Boston Area Diabetes and Endocrinology Research Center (BADERC) Award DK135043. The Zeiss confocal system is supported by grant S10OD021577-01.

